# Spatial patterns from a dispersal limitation perspective: revealing biotic interactions

**DOI:** 10.1101/2022.02.10.480007

**Authors:** Michael Kalyuzhny, Jeffrey K. Lake, Annette M. Ostling

## Abstract

Local spatial distributions of populations are often studied in comparison to Complete Spatial Randomness (CSR) and are found to be ubiquitously aggregated, likely due to dispersal limitation. Here we theoretically examine the advantages of comparing observed distributions to simulated populations subject only to drift and Dispersal Limitation (DL). Compared to this DL null, local competition produces overdispersion out to surprisingly large scales—much larger than the scale of competitive interactions. Furthermore, strong overdispersion provides a hallmark that a key requirement of stable coexistence is met, as it can only be observed if intraspecific competition is substantially stronger than interspecific competition. Dispersion compared to CSR is insensitive to competition and as a result unreflective of its presence. Hence, we suggest DL as a complement to CSR since the former focuses on biologically relevant spatial scales and has the potential to detect biotic interactions and habitat specificity.

## Introduction

The spatial distributions of species are considered among their most fundamental properties, begging for mechanistic explanation. Furthermore, most of the processes that shape ecological communities and their diversity and composition operate in a spatial context and leave a spatial mark. Consequently, spatial distributions contain information on the processes that created them and can be used to study these processes if analyses are constructed to properly disentangle them (McIntire & Fajardo 2009; Wiegand & Moloney 2013).

Here we focus on single species distributions at small scales, where individuals are often conceptualized as points in 2D space and the population distribution as a set of points or “(univariate) point pattern” (Wiegand & Moloney 2013). The most basic perspective on such patterns entails drawing a fundamental division between cases when points are close to each other or dense (aggregated), and cases when they are far or sparse (overdispersed). “Close/dense” and “far/sparse” are considered with respect to Complete Spatial Randomness (CSR) or Poisson null model, where points appear randomly, homogenously, and independently throughout the landscape (a random distribution, Wiegand & Moloney 2013). Mechanistically, it has been argued that intraspecific aggregation can be caused by dispersal limitation (Chave *et al*. 2002; Svenning & Skov 2002; Law *et al*. 2003); a spatially clumped habitat (Wiegand *et al*. 2007; Bagchi *et al*. 2011; Brown *et al*. 2016); and by positive interactions (Molofsky *et al*. 2001). Overdispersion, on the other hand, is primarily the result of negative interactions between individuals creating Conspecific Negative Density Dependence (CNDD, Chave *et al*. 2002; Law *et al*. 2003; Begon & Townsend 2020).

Despite its promise, linking spatial patterns and processes proves challenging. First, a given spatial pattern could arise from several different processes (McIntire & Fajardo 2009; Wiegand & Moloney 2013), making it difficult to distinguish between them using the observed pattern alone. Moreover, in forests, the distribution of the vast majority of tree species proves to be aggregated (Pielou 1962; Hubbell 1979; Plotkin *et al*. 2000; Seidler & Plotkin 2006) often at all measured spatial scales (Condit *et al*. 2000). This limits the usefulness of the fundamental division between ‘aggregation’ and ‘overdispersion’, since the latter is very rarely observed, and complicates mechanistic interpretation since aggregation has many possible causes.

The ubiquitous spatial aggregation of adult trees also contradicts the widespread evidence for strong CNDD acting on juveniles (Comita *et al*. 2010; Mangan *et al*. 2010; Johnson *et al*. 2012; Lebrija-Trejos *et al*. 2016), which should generate overdispersion. This evidence for CNDD was obtained by looking at the demography of single life stages (e.g., survival of first-year seedlings) in response to conspecific density. However, CNDD at one life stage could be counteracted by processes at other life stages (Detto *et al*. 2019; Hülsmann *et al*. 2020). Moreover, much of the aforementioned evidence for CNDD potentially suffers from statistical biases (Detto *et al*. 2019). Analyzing the spatial distributions of adults could complement the demographic approach and potentially evaluate the overall cumulative effects of CNDD on the distribution and dynamics of adults. Such effects, however, are rarely found, as there is currently very limited evidence that the effects of CNDD propagate to adults (Mittelbach & McGill 2019), suggesting that CNDD is overwhelmed, diluted, or obscured by the forces creating aggregation (as predicted by Chave *et al*. 2002; Brown *et al*. 2016). Since CNDD and other effects of biotic interactions are of primary importance for population and community dynamics (Hülsmann *et al*. 2020), it is very important to distinguish between the aforementioned scenarios: Does CNDD play no substantial role in the dynamics, or alternatively, do the effects of CNDD on adult distributions need to be teased apart from a backdrop of aggression?

Several approaches were developed for addressing this question. Already Pielou (1962) suggested truncating the distribution of nearest neighbor distances as a solution. In recent years, it was proposed to compare the observed pattern of individuals to a heterogenous Poisson null model, where different parts of the landscape have different expected densities (Wiegand *et al*. 2007). These methods require deciding how to “smooth” the density of individuals or to truncate the distribution of nearest neighbor distances. Furthermore, the mechanistic interpretation of the results is not always clear, since it is not evident what processes were “removed” by the method (Roughgarden 1983). Other works used the locations of juveniles as a null for the expected locations of adults (Condit *et al*. 2000; Bagchi *et al*. 2011), but these works found at most limited evidence for conspecific effects driving adult distribution patterns.

While multiple mechanisms shape community dynamics and spatial patterns, species must first disperse to a location, and only then they can interact with the a-biotic and biotic environment (“dispersal filtering”, e.g. Kraft *et al*. 2015; Cadotte & Tucker 2017). Indeed, models of dispersal assembly, where species disperse and go extinct at random, are often used as null models for community dynamics and as a baseline for detecting biotic and a-biotic interactions (Hubbell 2001; Gotelli & Ulrich 2012; Kalyuzhny *et al*. 2022). However, despite this “preemption” of dispersal, and despite considerable evidence for the importance of dispersal limitation in multiple systems (Hubbell *et al*. 1999; Muller-Landau *et al*. 2008; Losos & Ricklefs 2010), the CSR null model typically used for analyzing spatial patterns (as well as its cluster extensions, Plotkin *et al*. 2000; Wiegand *et al*. 2009), distributes the randomized individuals all across the survey plot – a scale that has no biological relevance. For an uncommon species with limited dispersal, this null may distribute most individuals to many parts of the landscape inaccessible due to dispersal limitation, and in comparison, the observed pattern will seem strongly aggregated. For example, the hypothetical population in Figure 1a, subject to both dispersal limitation and CNDD, is aggregated compared to CSR (Figure 1b). However, it may be considered overdispersed (having fewer neighbors and longer distances to them, Figure 1d-e) compared to a null with only dispersal limitation (Figure 1c). Hence, the CSR null model may create unrealistic and mechanistically unjustifiable low density and long distances to neighbors, potentially obscuring the short-distance repulsion effects of CNDD and masking habitat specificity.

**Figure 1.**
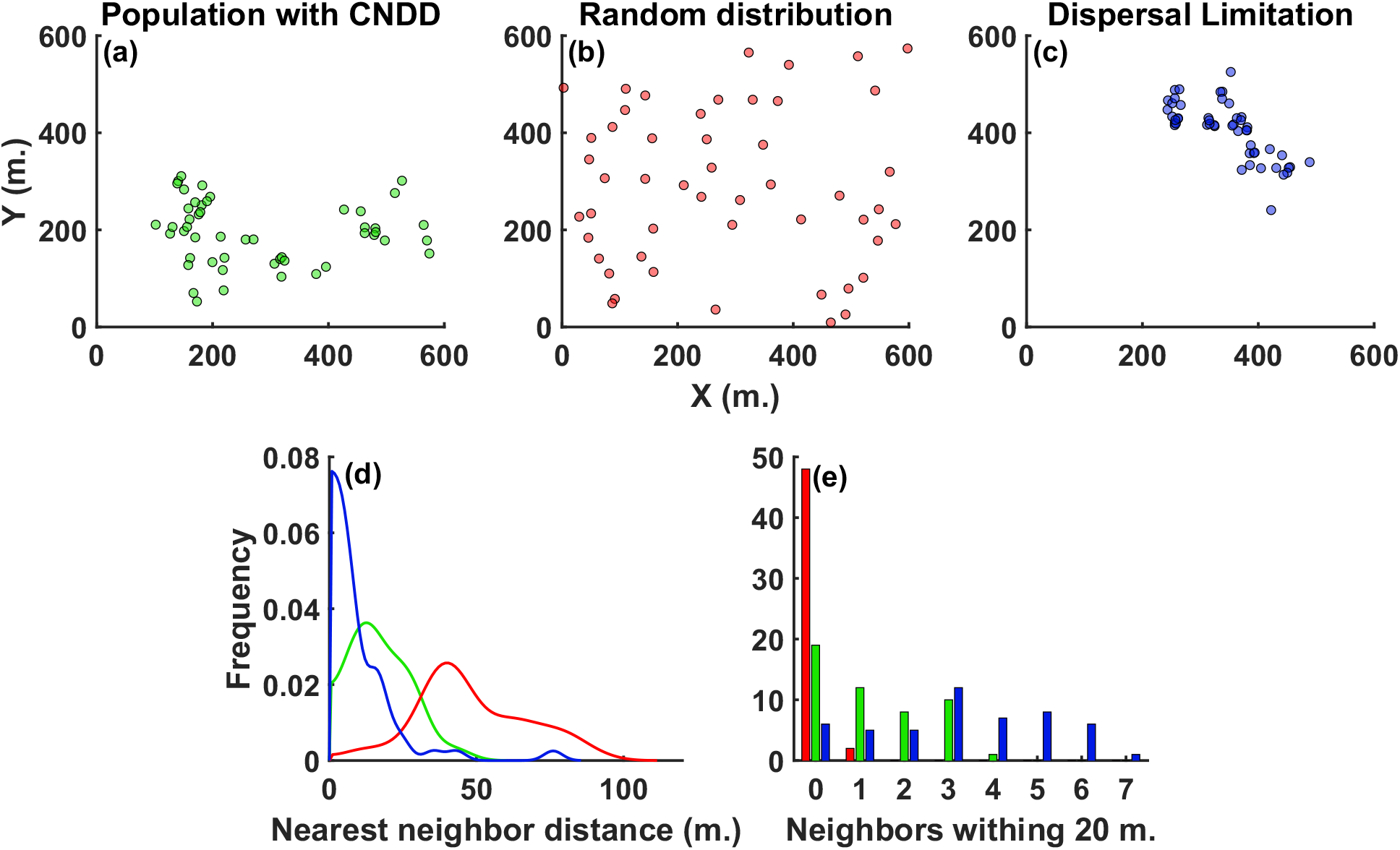
Comparison of the spatial point patterns of (**a**) a simulated population subject to both dispersal limitation and CNDD to a corresponding (**b**) random (CSR) and (**c**) Dispersal Limitation (DL) null models. The simulated population has 50 individuals, dispersal distance of 20 meters, CNDD operating to 7 meters and CNDD strength of *Q*_*C*_ = 5 (see Simulation Models section for details). (**d**) presents the (kernel-smoothed) distribution of nearest neighbor distances and (**e**) presents the distribution of the number of neighbors within 20 meters of each individual for the three cases above. These means of the distributions in (**e**) and (**d**) are used to compute the spatial statistics (see Spatial Statistics section). In all cases, green: simulated population, red: random distribution, blue: DL. The population is more aggregated than random but less than the DL null.

Here we suggest and investigate a new approach for analyzing spatial patterns, namely comparing them with a Dispersal Limitation (DL) reference (null) model. The goal is that this reference will incorporate the aggregation generated by dispersal and hence distribute the individuals at biologically relevant scales, allowing comparisons with it to reveal the effects of biotic interactions and habitat specificity. Such comparisons are now possible due to available estimates of seed dispersal kernels (e.g., Muller-Landau *et al*. 2008; Wright *et al*. 2008).To study the characteristics of this approach and strengthen the link between spatial patterns and processes, we conduct an extensive spatially explicit simulation study of the response of measures of aggregation (with both DL and CSR as nulls) to the magnitude of CNDD and Heterospecific Negative Density Dependence (HNDD) and to the scales of CNDD and dispersal. We hypothesize that density dependence will generate overdispersion (which would increase with its magnitude) compared to DL, while a comparison to CSR will detect aggregation if dispersal is limited. We expect such overdispersion to occur on scales comparable to the scale of CNDD. We further hypothesize that patterns compared with CSR will be mostly sensitive to dispersal distances (Chave *et al*. 2002) while being insensitive to CNDD, while a comparison with DL will show substantial sensitivity to CNDD.

## Simulation models

### The Dispersal Limitation (DL) null model

DL is a spatially explicit, continuous space neutral simulation model aimed at answering: what are the expected species-level spatial patterns in the community if the only processes taking place are dispersal limitation and drift? Hence, it should imitate the data in terms of the landscape size, density of individuals of all species and dispersal, but include no interactions. We recommend using estimates of dispersal kernels obtained using seed trap data or mechanistic models (e.g., Muller-Landau *et al*. 2008; Wright *et al*. 2008). If species differ in their dispersal kernels or size, we recommend running a different simulation for each species with the species-specific dispersal kernel and density of comparable individuals.

Despite focusing the analysis on a single species, we use a multispecies zero-sum community simulation with some low level of immigration from outside. Because the species are neutral, and using a sufficient number of species, the distributions of different species could be considered as independent realizations of the null. Some limited level of immigration must therefore be included in the model to generate diversity and differences between species. Care must be taken not to choose a high immigration level that would influence spatial distributions or make common species unlikely to occur (due to the deterministic negative effect of immigration, Chisholm & O’Dwyer 2014).

For our theoretical analysis, and to facilitate comparison, we generated the DL realizations using the same general simulation model with the same parameters as we used for communities with density dependence, but we set the magnitude of the latter to zero.

Since densities and distances between individuals strongly depend on abundance (Clark & Evans 1954), we recommend comparing them only to samples of the DL simulation (regardless of species identity, as they are neutral) that have abundances similar to the observed. For our analyses, we binned all populations by abundance in bins of increasing width (see Appendix S4) and compared the synthetic populations to null populations of the same abundance bin.

### General simulation model structure

While competition and apparent competition are often (Comita *et al*. 2010; Hülsmann *et al*. 2020) separated into CNDD and HNDD, we believe that it is more biologically justified to assume that all trees exert on all juveniles some General Negative Density Dependence (GNDD, representing competition for shared resources such as light and soil nutrients), while conspecifics also exert an added CNDD effect. Hence, in our simulations we manipulate the level of GNDD, and an added level of CNDD. The level of HNDD is equal to the level of GNDD we assume, while the level of CNDD is equal to the level of GNDD, plus the added CNDD. The situation CNDD > HNDD corresponds in our formulation to added CNDD > 0.

To generate both synthetic communities where CNDD and other forces are at work, and their null expectations, we simulate a community of sessile organisms on a 2D landscape with toroidal boundary conditions, inhabited by a fixed number of *J* individuals (see Appendix S1 for a glossary). Each time step, with probability *m*, a seed arrives from the species pool to a random location, or, with probability 1 – *m*, a random local individual is chosen to reproduce. The offspring is dispersed using the distance distribution *f*(*r*) with mean distance *D*, and upon arrival, the seed (locally produced or immigrant) establishes to become an adult with probability 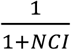 where *NCI* is the Neighborhood Crowding Index.

*NCI* is the summed contribution of the competitive effects of all individuals in the neighborhood *N*:

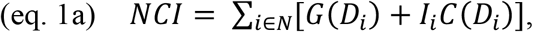

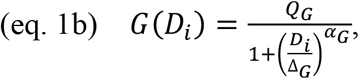

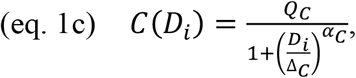

where an individual *i* at distance *D*_*i*_ exerts *GNDD, G*(*D*_*i*_), and added CNDD, *C*(*D*_*i*_) if it is a conspecific, with *I*_*i*_ being an indicator variable for the latter. *Q*_*G*_ and *Q*_*C*_ are the per-capita magnitudes of GNDD and added CNDD, respectively. Δ_*G*_ and Δ_*C*_ are the distances at which GNDD and added CNDD, respectively, fall to half of their magnitude, and *α*_*G*_ and *α*_*C*_ set how rapidly this decay occurs, with higher values meaning the competition force remains high out to distances closer to to Δ_*G*_ and Δ_*C*_, and then declines more quickly with distance from there (see Figure S1 for a visualization of eq. 1b-1c). We always consider the neighborhood *N* to consist of all individuals up to a distance of 2·Δ_*C*_ or 15 meters, whichever is greater, from the seed. Following successful recruitment, a random individual is chosen to die.

### Simulated parameters

#### General settings and dispersal

To roughly resemble a tropical forest plot (where such patterns have been extensively studied), we set the landscape to be a quadrat with edge length *L* = 600 meters and *J* = 5,500 individuals, resembling the density of adult trees in Barro Colorado Island, Panama (Condit *et al*. 2019). Immigration is set to occur with probability *m* = 5·10^−4^ from a regional pool of *S* = 300 species with equal frequency. *m* was chosen to be low to avoid an effect on local spatial patterns, but still high enough to maintain many species with variable abundances.

In line with several empirical works on dispersal, we set *f*(*r*) to be a fat-tailed 2DT with 3 degrees of freedom (Clark *et al*. 1999; Muller-Landau *et al*. 2008). We study a wide regime of parameters to test the robustness of the observed patterns and to examine the sensitivity of different statistics to changes in the parameters.

#### GNDD and CNDD

We set the distance of GNDD to resemble the overall density in tropical forests: Δ_*G*_ = 4.5 meters, representing roughly the crown diameter of adult trees, and *α*_*G*_ = 5, so that at a distance of 20 meters the effect drops to almost zero (∼0.0005Q_G_, Uriarte *et al*. 2005).

Four mean dispersal distances (*D*) and CNDD distances (Δ_*C*_) are used: 7 meters, 20 meters, 60 meters and 3000 meters (effectively infinite distance). For CNDD, we simulated each of these four Δ_*C*_s with a corresponding steepness parameter value (*α*_*C*_): 5, 6, 7, 10, respectively (each chosen to create a relatively quick decline in CNDD at Δ_*C*_, which requires a higher value for higher Δ_*C*_). We assume that CNDD is operating to a larger distance than GNDD because natural enemies, the primary hypothesized cause of CNDD (Hülsmann *et al*. 2020), are capable of dispersal and movement. The dispersal distances are also set to represent realistic variation in forests (Clark *et al*. 1999; Muller-Landau *et al*. 2008).

To study the effect of the magnitude of CNDD and GNDD, we use all combinations of *Q*_*G*_ and *Q*_*C*_ of 0, 0.2, 1 and 5.

To consider scenarios where different species suffer different levels of CNDD (as found by e.g., Comita *et al*. 2010; Lebrija-Trejos *et al*. 2016), we further run simulations where each species has its own value of *Q*_*C*_, *Q*′_*C*_, and these values are distributed between *Q*_*Cmin*_ and *Q*_*Cmin*_ + *Q*_*Crange*_ equidistantly. For these simulations, we used the same dispersal and CNDD distances as above, excluding the infinite distance cases. We used *Q*_*Cmi n*_ = 1 and two levels of *Q*_*Crange*_, 1 and 5. *Q*_*G*_ was assumed to be 0.2 in these simulations.

#### Simulation time and sampling

Each simulation was initiated as a random placement of *J* individuals sampled from the pool and was given 3000 generations (*J* time steps each) to equilibrate. For the simulations with fixed CNDD, four levels of the four parameters (*D*, Δ_*C*_, *Q*_*G*_ and *Q*_*C*_) were run. Following that, 3000 snapshots of each simulation were taken, with 10 generations between snapshots. For the simulations with variable CNDD, three levels of *D* and Δ_*C*_ and two levels of *Q*_*Crange*_ were used. 9 independent runs were used for each regime, each with 1000 snapshots taken after equilibration and 50 generations between snapshots,

In the results we present the mean values of the statistics for all populations with abundance ≥ 5 recorded in a parameter regime, sometimes subdivided by abundance when this is indicated.

## Spatial statistics

The two most common approaches to summarizing univariate point patterns are to examine the distances of individuals to their nearest neighbors (Figure 1d) or the density of neighbors around each individual (Figure 1e) at different distances, which are classically compared to their values under CSR. In particular, the Clark-Evans (*CE*) statistic compares the average distance of an individual to its conspecific nearest neighbor in the observed data, *NND*_*obs*_, to its expectation under CSR, which equals 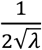, where *λ* is the density of the species (Clark & Evans 1954):

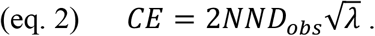

If *CE* > 1 the population is overdispersed compared to CSR, while *CE* < 1 indicates aggregation.

We propose the Excess neighbor Distance (*ED*) statistic, an analogous measure that is symmetric on a logarithmic scale. Distances are compared to the mean nearest neighbor distance in realizations of the DL model that have similar abundance, *NND*_*null*_:

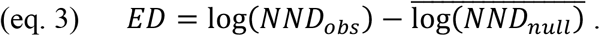

The averaging is done over different realizations of DL. Positive values of *ED* indicate overdispersion with respect to DL while negative values indicate aggregation.

To study the density of neighbors, many works use the Relative Neighborhood Density statistic (*RND*, also known as pair-correlation function; Condit *et al*. 2000; Wiegand & Moloney 2013). *RND*(*r*) compares the observed number of conspecific neighbors that an average individual has between distances *r* and *r* + Δ*r, N*_*obs*_(*r*), to its expectation under CSR, *A*(*r*)*λ*, where *A*(*r*) is the area of the annulus with inner and outer radii *r* and r + Δ*r*, respectively:

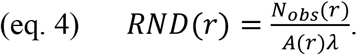

If *RND*(*r*) > 1, the population is aggregated compared to CSR at distances r to r + Δ*r*, while *RND*(*r*) < 1 indicates overdispersion. We propose the analogous Excess neighborhood Abundance (*EA*) statistic,

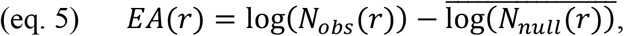

where *N*_*null*_(*r*) is the analog of *N*_*obs*_(*r*) in a comparable realization (in terms of abundance) of the DL null. Positive values of *EA* indicate aggregation with respect to the DL, while negative values indicate overdispersion. In practice, we study *EA* and *RND* at 10-meter distance bins, and we also focus on a circle of 20 meters (*EA*(*20*) and *RND*(20)).

While direct analogs of RND and the Clark-Evans statistics could be devised for use with DL, *ED* and *EA* are constructed on a logarithmic scale, to create symmetry between aggregation and overdispersion to better capture the variation in spatial dispersion (Wiegand *et al*. 2007). Intuitively, if *ED* or *EA*(*r*) have a value of *x*, it indicates that the average nearest neighbor distance or number of conspecific neighbors at distance *r*, respectively, are larger by e^*x*^ then their expectation, which is the geometric mean over realizations of the null. The statistical significance of both *EA* and *ED* can be evaluated by comparing log(*N*_*obs*_(*r*)) or log(*NND*_*obs*_) with the distribution of their analogs under the nulls with a two-sided test.

## Results and discussion

### Overdispersion and its scale

As we expected, dispersion patterns (averaged across species) appear aggregated compared to CSR as long as dispersal is not global. (Figures 2a, S3, S5). Since typical clumps are large, this aggregation extends up to scales ∼ 150 meters (Figure 2a). On the other hand, when compared to DL, Both GNDD and added CNDD, separately and in combination, generate overdispersion relative to DL (Figures 2-4, S2, S4). However, counter to our expectations, this overdispersion occurs at scales much larger than the scales of density dependence or dispersal (Figure 2b), and even GNDD by itself (acting at a distance ∼ 5 meters) can create overdispersion on scales of dozens of meters.

**Figure 2.**
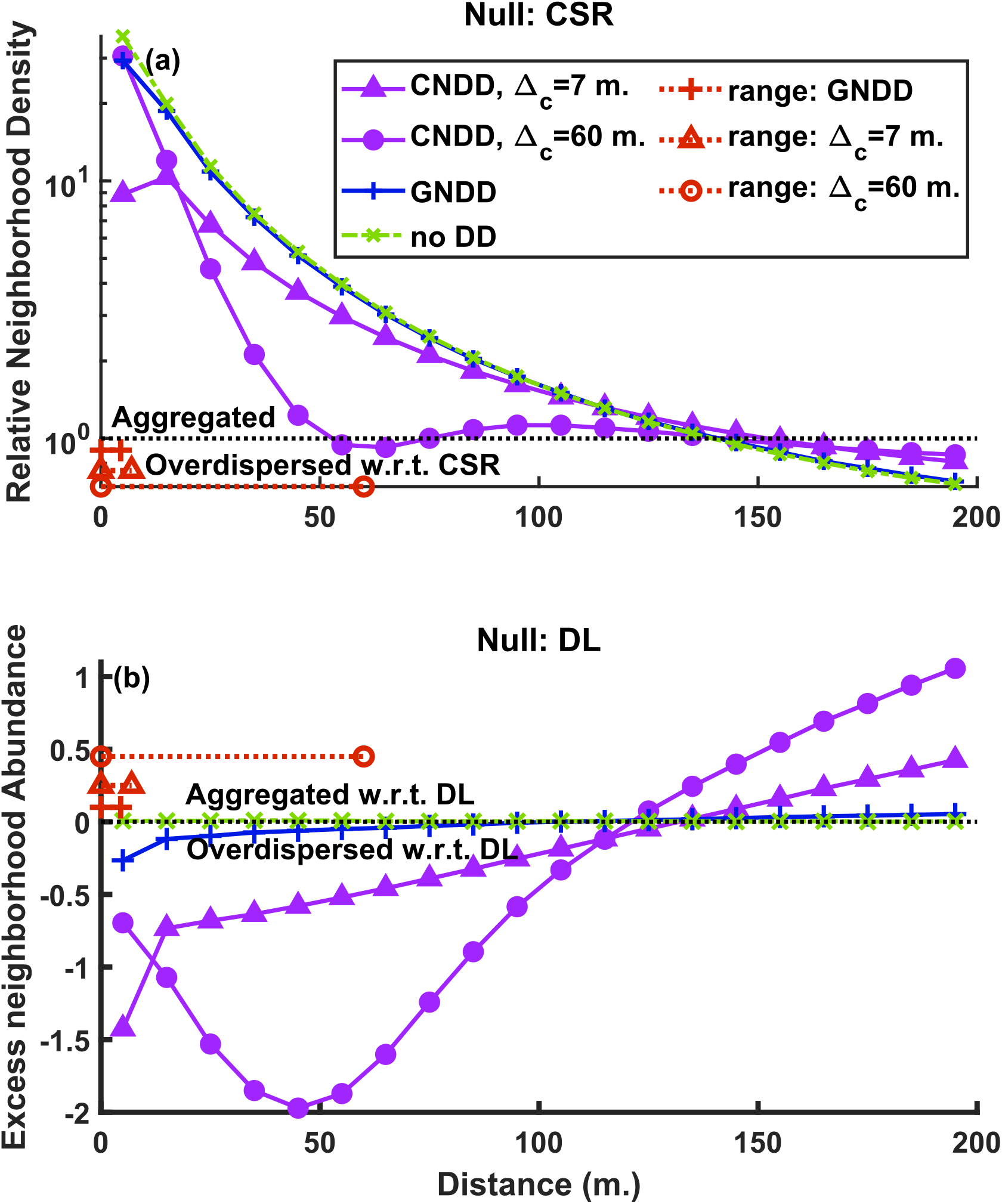
Aggregation and overdispersion of simulated populations at different scales, compared with (**a**) the Complete Spatial Randomness (CSR) null, using Relative Neighborhood Abundance (Eq. 4), and (**b**) the Dispersal Limitation (DL) null, using Excess Abundance (Eq. 5). Mostly aggregation is seen in (**a**), where in (**b**) the overdispersion created by CNDD is made apparent, and it is seen out to large scales. Dispersal distance (*D*) in all cases is 20 meters. Purple curves with triangle and circle symbols are cases with CNDD only (*Q*_*G*_ = 0, *Q*_*G*_ = 5) operating to distances (Δ_*G*_) of 7 and 60 meters, respectively. The blue and green curves represent simulations with only GNDD (*Q*_*G*_ = 5, *Q*_*G*_ = 0) and no density dependence (*Q*_*G*_ = 0, *Q*_*G*_ = 0), respectively. Full correspondence with the null is represented with a grey dotted line. Red lines present the distance scale of density dependence of the corresponding CNDD or GNDD scenario (indicated by the symbol).

We believe this result represents the emergence of large-scale structure from local interactions, which is common in many ecosystems (Levin 1992). The neighbors of each individual tend to be overdispersed, but so are their neighbors, and so forth. Hence, if large-scale overdispersion is observed in nature, it likely stems from CNDD acting on considerably shorter scales.

Another interesting finding is the appearance of a trough in both the *EA*(*r*) and *RND*(*r*) curves when added CNDD operates at long distances (dozens of meters), roughly at the typical scale of CNDD. This is the result of the emergence of clumps of a few individuals that are surrounded by large vacant areas (Figure S6).

Note that the high *EA* and low *RND* at large distances are an unavoidable result: since the overall number of individuals is fixed, having more/less neighbors than expected at short distances necessitates having less/more at longer distances. Still, these patterns are typically interpreted focusing on short distances (Wiegand & Moloney 2013).

See appendix S2 for an analysis of the effects of added CNDD distance scale and dispersal distance scale on spatial statistics.

### Effects of the magnitude of CNDD and GNDD

As we expected, an increase in both GNDD and added CNDD, when they operate separately, increases overdispersion compared to DL (Figures 3, S7a,b). However, the effect of GNDD by itself on overdispersion in the absence of added CNDD is much weaker than the effect of added CNDD by itself (Compare the effect of changes in *Q*_*G*_ when *Q*_*C*_ = 0 to changes in *Q*_*C*_ when *Q*_*G*_ = 0 in Figure 3a,b). Moreover, when both GNDD and added CNDD operate (as is expected in natural systems), their effects counteract each other: if added CNDD is strong, the effect of increasing Q_G_ is *reversed to reducing overdispersion*. If GNDD is strong, increasing Q_C_ still increases overdispersion, but at a slower rate (Figures 3a,b, S7a,b). Overall, the strongest overdispersion can only be observed when added CNDD >> GNDD and therefore strong overdispersion could indicate this scenario (bright yellow colors indicating high overdispersion occur only in the lower right in Figures 3a,b). Finally, using CSR as null, *CE* shows a unimodal shaped dependence or limited dependence on *Q*_*C*_ (Figure 3d), while the behavior of *RND*(20) is consistent with *ED* and *EA*(20) (Figure S7d).

**Figure 3.**
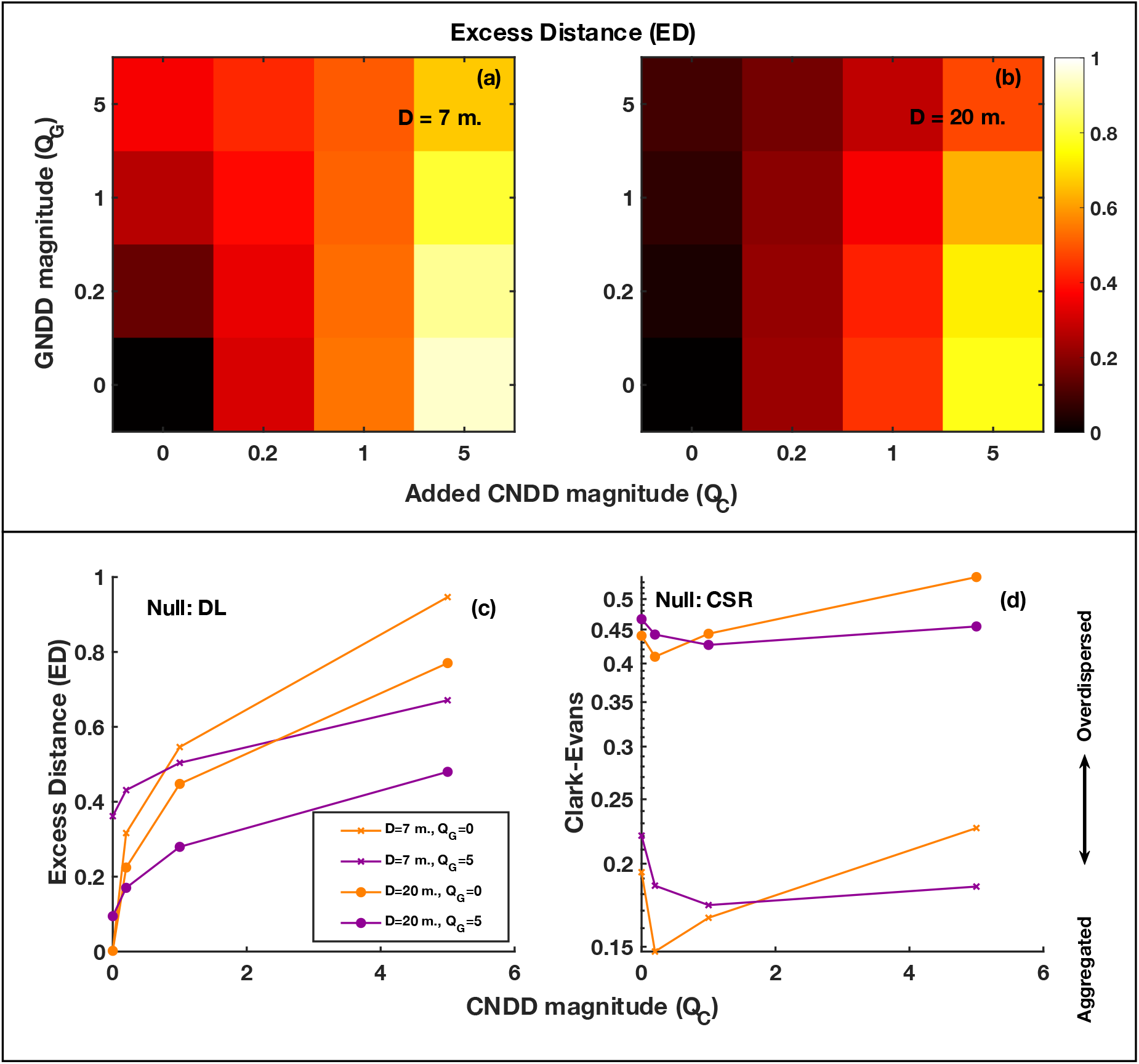
Effect of the magnitude of GNDD (*Q*_*G*_) and added CNDD (*Q*_*C*_) on nearest neighbor statistics. In all cases Δ_*G*_ = 20 meters. The colors of (**a**) and (**b**) represents the Excess nearest neighbor Distance (*ED*, eq. 3) in response to a combination of GNDD and CNDD for dispersal distances (*D*) of 7 meters and 20 meters, respectively. Furthermore, the effect of *Q*_*C*_ on (**c**) *ED* and on (**d**) the Clark-Evans statistic (eq. 2), with the latter using CSR as null, are presented when *Q*_*G*_ = 0 (orange) or *Q*_*G*_ = 5 (purple). GNDD alone can produce much lower overdispersion than added CNDD. GNDD counteracts the effect of added CNDD and the strongest overdispersion is observed when CNDD is maximal and GNDD is minimal.

These results can be understood by considering the spatial uniformity of seed mortality. GNDD is caused by all individuals of all species similarly, causing a relatively spatially uniform mortality across the plot, while added CNDD generates “repulsion” only from conspecifics. Consequently, GNDD cannot cause high species-specific overdispersion, and it can homogenize the non-uniform mortality caused by CNDD, reducing the latter’s effect. Analogously, if CNDD is not distance dependent (Δ_*C*_ = ∞), it will homogenize the local effect of GNDD, and in this situation increasing *Q*_*C*_ may reduce overdispersion (Figure S8). Finally, the “peculiar” behavior of the Clark-Evans statistic, whereas increasing *Q*_*C*_ may increase aggregation, is probably due to the high sensitivity of this statistic to abundance.

### Effects of variation in abundance and CNDD across species

Thus far we analyzed the spatial statistics averaged across species. How do these patterns depend on species’ abundance? From the perspective of CSR, there is a strong effect of reduced aggregation with abundance, as is commonly found in tropical forests (Appendix S3, Condit *et al*. 2000). From the perspective of DL, the relationship is more complex and often unimodal or flat. See appendix S3 for more details and for an analysis of the underlying mechanisms.

Next, we examined the case when species differ in their species-specific susceptibility to CNDD (*Q*_*C*_*’*). We hypothesized that species that have higher *Q*_*C*_*’* would be more overdispersed. However, our simulations show that, from the perspective of statistics using DL as a null, the effect of *Q*_*C*_*’* is unimodal with species of intermediate *Q*_*C*_*’* being the most overdispersed, but sometimes only the increasing or decreasing phase is observed (Figure 4a,b). Qualitatively similar results are observed for lower values of *Q*_*Crange*_ (Figure S9). Surprisingly, statistics that use CSR as a null suggest that species with higher *Q*_*C*_*’* are more *aggregated* (Figure 4c,d).

**Figure 4.**
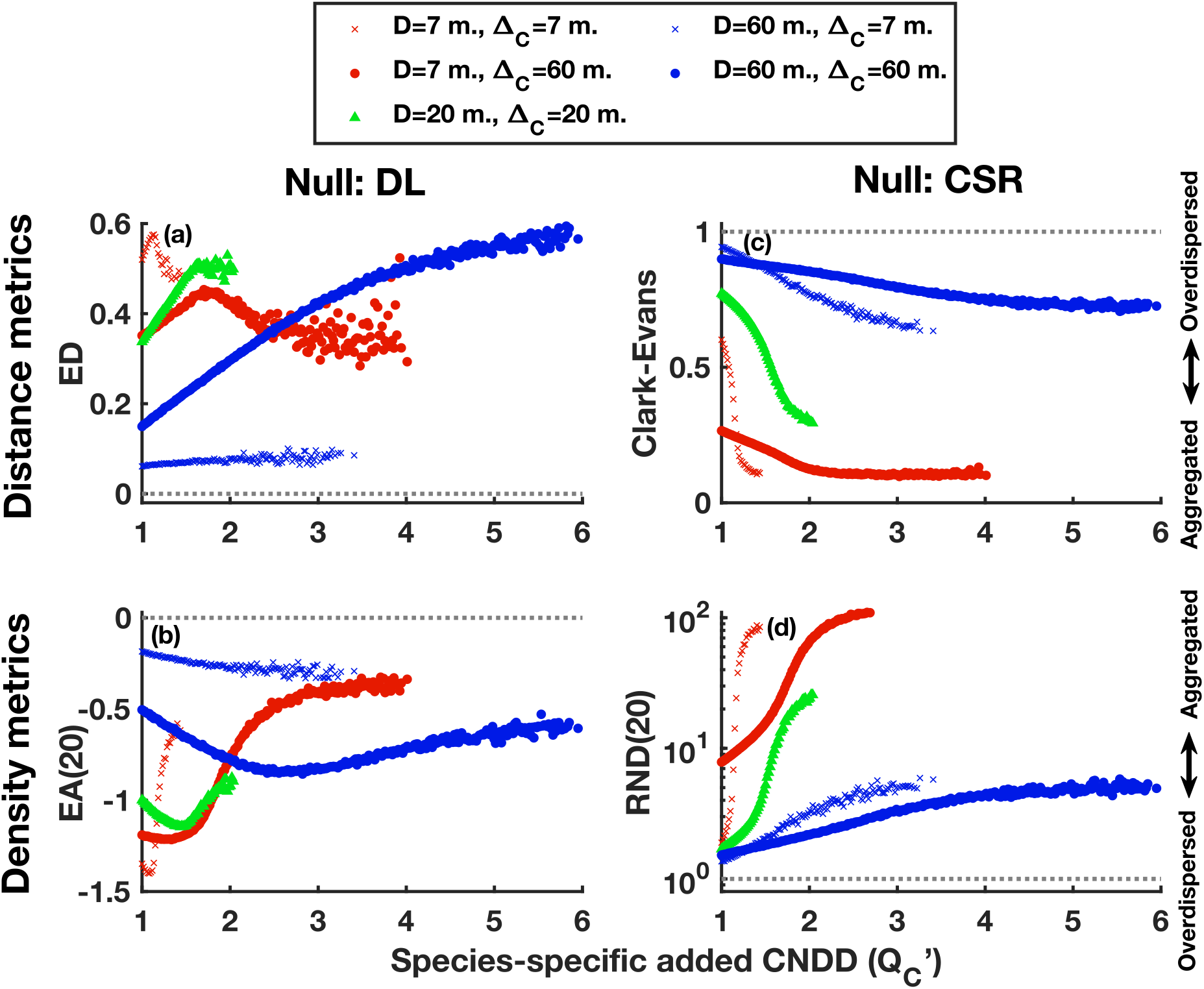
Effect of the species-specific magnitude of CNDD (*Q*_*C*_*’*) on spatial statistics: (**a**) Excess nearest neighbor Distance (*ED*); (**b**) Excess neighborhood Abundance within 20 meters (*EA*(20)); (**c**) the Clark-Evans nearest neighbor statistic and (**d**) Relative Neighborhood Density within 20 meters (*RND*(20)). The top row examines nearest neighbor distance statistics while the bottom row shows neighborhood density. The statistics on the left column have DL as reference while the right column has CSR. Different colors and symbols represent regimes with different levels of dispersal distance (*D*) and CNDD distance (Δ_*G*_), respectively. In all cases, GNDD magnitude (*Q*_*G*_) is 0.2 and CNDD has a minimum value *Q*_*C*min_ = 1 and *Q*_*C*range_ = 5. Species with fewer than 500 observations were discarded, hence, some parameter regimes do not have observations for high *Q*_*C*_*’*. Overdispersion depends unimodally on *Q*_*C*_*’* when using the DL null, while compared to CSR species with higher *Q*_*C*_’ appear more aggregated.

These non-trivial relationships are a result of the effect of *Q*_*C*_*’* on abundance: species with higher *Q*_*C*_*’* are substantially rarer in our model. Consequently, increasing *Q*_*C*_*’* affects dispersion both directly, increases the repulsion effect (as in Figure 3), and indirectly, by making species rarer. With the statistics that use CSR as reference, the reduced abundance causes a sharp increase in aggregation, creating the counter-intuitive increase in aggregation with CNDD. With the statistics that are compared to DL, the typical effect of abundance is unimodal (see Appendix S3), and this adds to the increase in repulsion with stronger *Q*_*C*_*’* to shape the unimodal effect of *Q*_*C*_*’* on overdispersion.

### Sensitivity to different parameters

Figure 5 presents the effect of changing one parameter at a time from an intermediate level to a high level, while keeping the others fixed, across the distributions of the other parameters. First, the statistics that use CSR as null are primarily sensitive to dispersal distance, far beyond their sensitivities to competition (in line with the findings of Chave *et al*. 2002). When using DL as null, the *ED* statistic is sensitive to the different parameters to a comparable degree, while *EA*(20) is primarily sensitive to the magnitude of CNDD, more than to other parameters. Finally, all statistics are weakly affected by GNDD. Hence, we expect the DL null to have high capacity to detect CNDD in empirical data, especially with the *EA*(20) statistic, unlike the CSR null.

**Figure 5.**
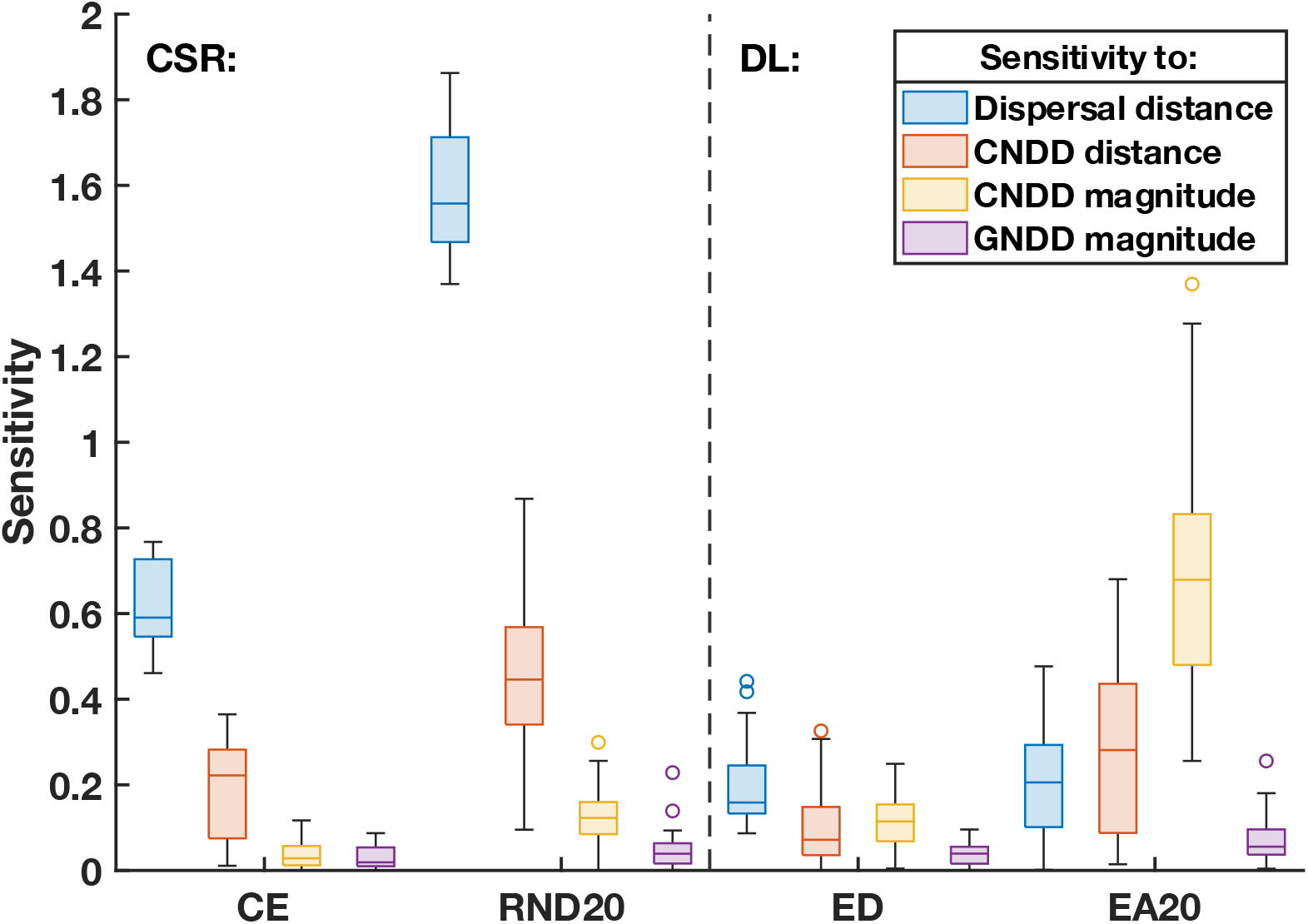
Sensitivity of the statistics using DL and CSR as reference to changes in the simulation parameters. Sensitivities are calculated as the effect on the mean value of the statistic (on a logarithmic scale, hence *RND*(20) and Clark-Evans are log transformed for comparison) of changing one parameter at a time from an intermediate level (D = 20 m. or Δ_*G*_ = 20 m. or *Q*_*C*_ = 0.2 or *Q*_*C*_ = 0.2) to a higher level (D = 60 m., Δ_*G*_ = 60 m., *Q*_*C*_ = 1, *Q*_*C*_ = 1, respectively). The distributions of these sensitivities, each calculated by comparing a pair of simulations differing in the parameter of interest, is presented across all the values of the other parameters. Statistics based on CSR have low sensitivity to CNDD and high sensitivity to dispersal distance, while statistics based on DL are much more sensitive to CNDD.

## Conclusions and overview

Ecologists have long sought an approach to gaining insight into process from spatial patterns within species, but this goal has been elusive. Our theoretical analysis demonstrates the usefulness of spatial patterns for understanding the cumulative effect of density dependent interactions on adult individuals in natural communities, once they are compared with predictions based on dispersal limitation, rather than complete spatial randomness. First, it is of fundamental importance to evaluate the magnitude of added CNDD, or, equivalently, the degree to which CNDD > HNDD, as this is the requirement for stabilizing niche differences, which may lead to coexistence if fitness differences are sufficiently small (Hülsmann *et al*. 2020). Our analysis reveals that high levels of overdispersion compared to DL are only possible if added CNDD is substantial and can be used to detect this ecologically important situation. This stems from the combination of two factors: first, in the absence of added CNDD (when CNDD = HNDD = GNDD, left column of Figures 3a,b), GNDD is only capable of generating limited values of overdispersion, at least for dispersal distances of 20 meters (typical of tropical forests, Muller-Landau *et al*. 2008). On the other hand, much higher levels of overdispersion are enabled by added CNDD. Second, GNDD counteracts added CNDD, so if both are present, overdispersion will be reduced compared to a situation when GNDD is weak. Hence, strong GNDD and weak added CNDD are inconsistent with high overdispersion.

Among the information that can be obtained from spatial patterns are the relative scales of CNDD and dispersal. If dispersal acts on longer distances than CNDD, as is believed (Janzen 1970), we show that *EA*(*r*) should decrease with *r* monotonically. Moreover, it is possible to correlate measures of CNDD magnitude (e.g., Comita *et al*. 2010; Lebrija-Trejos *et al*. 2016) with values of *ED* and *EA* to test whether specific measures of CNDD are associated with adult distributions.

Another factor that can have a substantial effect on species’ distributions is habitat specificity. We did not include this important factor in our analyses since there are multiple established methods to quantify habitat specificity using spatial distributions (e.g., Harms *et al*. 2001; Wiegand *et al*. 2007; Flugge *et al*. 2014). We hypothesize that if the habitat is patchy, seeds would be lost to unsuitable habitats and this will reduce overdispersion and potentially even create aggregation compared to DL. Such aggregation may further be reduced at short distances owing to CNDD. We view such aggregated patterns as much more informative than aggregation compared to CSR. Overall, the analysis of spatial patterns of adults can complement analyses of demography for understanding community dynamics. Specifically, explaining the variation in *ED* and *EA*(20) among species can detect the relative importance of factors shaping species’ distributions, and our work suggests that such analyses should include abundance and dispersal distance as covariates, since they have a substantial effect on these statistics.

Nevertheless, our results emphasize that spatial patterns observed in nature are a complex emergent phenomenon. While we initially expected that the magnitude and scale of overdispersion could serve as a measure of the magnitude and scale of CNDD, our results show this is not straightforward: even with all else being equal, we find that the relationship between CNDD and overdispersion may not be monotonic, so that species subject to stronger CNDD may be *less* overdispersed (Figure 4), let alone if they differ by other factors such as dispersal distance. The scale of emerging population overdispersion is also found to exceed the scale of CNDD caused by individuals. Finally, overdispersion compared to DL is the combined product of GNDD and added CNDD that counteract each other, up to the point that each could reduce overdispersion under specific conditions.

A limitation of our approach is its dependence on reliable estimates of dispersal distance that are difficult to obtain. We recommend that empirical analyses comparing spatial patterns to DL should always test the robustness of results by using multiple dispersal kernels, incorporating this uncertainty. Assuming too long dispersal distance is expected to bias estimates of overdispersion downwards, which could be more conservative if the goal is to detect CNDD. Finally, it is possible to analyze what dispersal distance is required to obtain the observed patterns, and if that distance is unrealistically large (e.g., 200 meters for trees) this could serve as evidence for CNDD.

The use of CSR as a reference for dispersion patterns is the simplest analysis of spatial patterns, that has been done for over a hundred years (Gleason 1920), often with the aim of detecting the signals of habitat specificity and biotic interactions (Condit *et al*. 2000; Plotkin *et al*. 2000; Bagchi *et al*. 2011). However, previous analyses (Chave *et al*. 2002) as well as this work, show that dispersal limitation will almost inevitably obscure the signal of these factors when using CSR. Moreover, we show that the expected dependence of spatial patterns compared to CSR on species’ properties such as the magnitude of CNDD makes little sense in some situations, likely due to their sensitivity to abundance. All this is due to considering the reference at a biologically irrelevant scale – the scale of the survey plot, over which the individuals are randomly distributed with no biological justification. Conversely, with our approach, distributions are compared to their expectations if only dispersal and drift were operating. Hence, we believe that DL sets the biologically relevant and interpretable reference for spatial patterns rooted in ecological theory, offering a complimentary definition of “aggregation” and “overdispersion”. we hope this work will facilitate its application to natural populations.

## Acknowledgements

We thank S.J. Wright, Ronen Kadmon and members of the Ostling lab for helpful comments on the manuscript. This research was supported in part through computational resources and services provided by Advanced Research Computing at the University of Michigan, Ann Arbor. MK was supported by a Michigan Life Sciences Fellowship, the Zuckerman STEM Leadership Program, funds from MCubed and the Associate Professor Support Fund at the University of Michigan. JKL acknowledges Adrian College and the University of Michigan for sabbatical support.

## Statement of authorship

MK conceptualized the study, all authors planned the study, MK performed the analysis and wrote the first draft of the manuscript and all authors contributed substantially to interpreting results and manuscript revisions.

## Data accessibility statement

this work does not use any data. All code used to obtain the results is available in Appendix S4.

