## Appendix S4 for "Spatial patterns from a dispersal limitation perspective: revealing biotic interactions": Analyze_var_CNDD2.m

```
%%%%%%%%%%%%%%%%%%%%%%%%%%%%%%%%%%%%%%%%%%%%%%%%%%%%%%%%%
%%% Analyze the variable CNDD simulations
%%%%%%%%%%%%%%%%%%%%%%%%%%%%%%%%%%%%%%%%%%%%%%%%%%%%%%%%%
%
% Date created: 15/12/2021
% Date last modified: 28/01/2022
%
% Author: Michael Kalyuzhny
%
% Merges the statistics from different itterations
%
%% 
dists = {'7' '20' '60' '3000'}; %the distances used (in m.)

load('ext3_results/inp_all_var_CNDD.mat');
abd_bin_edges = [1 5:2:51 55:5:100 110:10:200 250:50:600 700:100:1000 1250 1500 5500];

si = size(inp_all); %remove last dimension
ittr = si(end); 
si = si(1:end-1);
regs = prod(si); %the total number of regimes

%full data:
JC_all = cell(si);
CE_all = cell(si);
ED_all = cell(si);
EA20_all = cell(si);
RND20_all = cell(si);
sp_abd_all = cell(si);

%species averages:
ED_sp_mean = cell(si);
EA20_sp_mean = cell(si);
CE_sp_mean = cell(si);
RND20_sp_mean = cell(si);
ED_sp_samp_size = cell(si);
EA20_sp_samp_size = cell(si);
CE_sp_samp_size = cell(si);
RND20_sp_samp_size = cell(si);

% Tables of abdxQ_C
samp_size_2D = cell(si);
EA20_2Dbin = cell(si); %FIRST DIM.: Abundance, Second DIM.: Q_C
ED_2Dbin = cell(si);
CE_2Dbin = cell(si);
RND20_2Dbin = cell(si);
Abd_2Dbin_centers = cell(si);
Q_C_2Dbin_centers = cell(si);


parfor rr = 1:regs
    
    rr
    %load reference data:
    ref = load(['imm_range_' num2str(round(inp_all(rr).a*sqrt(pi)*gamma(inp_all(rr).b-1.5)/(2*gamma(inp_all(rr).b-1)))) '_neut_analysis_results.mat']);
    
   
    %Compute CNDDs:
    if inp_all(rr).Q_CNDD_min == inp_all(rr).Q_CNDD_max %no variability...
        JC_all{rr} = inp_all(rr).Q_CNDD_min*ones(1,inp_all(rr).S_reg);
    else
        inc = (inp_all(rr).Q_CNDD_max-inp_all(rr).Q_CNDD_min)/(inp_all(rr).S_reg-1); %increment
        JC_all{rr} = ( inp_all(rr).Q_CNDD_min : inc : inp_all(rr).Q_CNDD_max ); %first species has minimal JC, last has maximal, evenly distributed.    
    end
    
    one_over_prop_area20 = (inp_all(rr).L^2)./(pi*400);
    
    %Load and analyze current regime:
    mean_dist = [];
    N20 = [];
    bins = [];

    for ii = 1:ittr %run over itterations of the same regime
        
        dat_now = load(['ext3_results/' inp_all(rr).output_file(1:end-1) num2str(ii) '.mat']);
        [mean_dist_now, N20_now, CE_now, bins_now, sp_abd_now, abd_bin_centers] = Stats_regime_tables(dat_now.com_samp, abd_bin_edges, 5, 600, nan);
        
        %add to previous results:
        mean_dist = [mean_dist ; mean_dist_now];
        N20 = [N20 ; N20_now];
        CE_all{rr} = [CE_all{rr} ; CE_now];
        bins = [bins ; bins_now];
        sp_abd_all{rr} = [sp_abd_all{rr} ; sp_abd_now];
    end
    
    
    %Prepare variables:
    ED_all{rr} = nan(size(mean_dist));
    EA20_all{rr} = nan(size(mean_dist));
    RND20_all{rr} = nan(size(mean_dist));

    for yy = 1:size(mean_dist,1)
        for sp = 1:size(mean_dist,2)
            bin_now = bins(yy,sp); %abundance bin
            if isnan(bin_now)
                bin_now = 1;
            end
            ED_all{rr}(yy,sp) = log(mean_dist(yy,sp)) - ref.mean_log_dist(bin_now); 
            EA20_all{rr}(yy,sp) = max(log(N20(yy,sp)), log(1/sp_abd_all{rr}(yy,sp))) - ref.log_N20(bin_now);
            RND20_all{rr}(yy,sp) = N20(yy,sp)*one_over_prop_area20/sp_abd_all{rr}(yy,sp);
        end
    end
    
    % divide into bins in 2D:
    % USED IN EARLIER VERSIONS
    
    JC_table = ones(size(CE_all{rr},1),1)*(JC_all{rr}(:)');
    tab_all = [ED_all{rr}(:)  EA20_all{rr}(:)  log(sp_abd_all{rr}(:)) CE_all{rr}(:) RND20_all{rr}(:) JC_table(:) ];
    to_rem = (isnan(tab_all(:,1))) | (~isfinite(tab_all(:,3))) ;
    tab_all(to_rem,:) = [];

    tab_all = array2table(tab_all,'VariableNames',{'ED', 'EA20', 'abd', 'CE', 'RND20', 'JC'});

    % Divide into bins:
    [N,Xedges,Yedges,binX,binY] = histcounts2(tab_all{:,'abd'}, tab_all{:,'JC'},15); % X: log abd.; Y: JC

    EA20_2Dbin{rr} = nan(size(N)); %FIRST DIM.: Abundance, Second DIM.: Q_C
    ED_2Dbin{rr} = nan(size(N));
    CE_2Dbin{rr} = nan(size(N));
    RND20_2Dbin{rr} = nan(size(N));
    
    % IMPORTANT! In each of these matrices, different rows have different abundance bins and different columnts have
    % different JC bins!
    % Hence, the correct way to plot this would be:
    % surf(Q_C_2Dbin_centers{rr}, Abd_2Dbin_centers{rr},ea)

    for xx = 1:length(Xedges)-1
        for yy = 1:length(Yedges)-1
            EA20_2Dbin{rr}(xx,yy) = mean(tab_all{(binX == xx) & (binY == yy), 'EA20'});
            ED_2Dbin{rr}(xx,yy) = mean(tab_all{(binX == xx) & (binY == yy), 'ED'});
            CE_2Dbin{rr}(xx,yy) = mean(tab_all{(binX == xx) & (binY == yy), 'CE'});
            RND20_2Dbin{rr}(xx,yy) = mean(tab_all{(binX == xx) & (binY == yy), 'RND20'});
        end
    end

    Abd_2Dbin_centers{rr} = Xedges(1:end-1)+diff(Xedges)/2;
    Q_C_2Dbin_centers{rr} = Yedges(1:end-1)+diff(Yedges)/2;
    
    samp_size_2D{rr} = N;
    
    %compute average (to make the data easier to load!)
    ED_sp_mean{rr} = nanmean(ED_all{rr});
    EA20_sp_mean{rr} = nanmean(EA20_all{rr});
    CE_sp_mean{rr} = nanmean(CE_all{rr});
    RND20_sp_mean{rr} = nanmean(RND20_all{rr});
    
    ED_sp_samp_size{rr} = sum(~isnan(ED_all{rr}));
    EA20_sp_samp_size{rr} = sum(~isnan(EA20_all{rr}));
    CE_sp_samp_size{rr} = sum(~isnan(CE_all{rr}));
    RND20_sp_samp_size{rr} = sum(~isnan(CE_all{rr}));
    
    
end

% WE ARE NOT SAVING THE HUGE TABLES LIKE ED_all

%minimalist version:
save('var_CNDD_all_analyses_results2.mat', 'Abd_2Dbin_centers', 'Q_C_2Dbin_centers', 'samp_size_2D', 'EA20_2Dbin', 'ED_2Dbin', 'CE_2Dbin', 'RND20_2Dbin', 'ED_sp_samp_size', 'EA20_sp_samp_size', 'CE_sp_samp_size', 'ED_sp_mean', 'EA20_sp_mean', 'RND20_sp_mean', 'RND20_sp_samp_size', 'CE_sp_mean',  'inp_all', 'JC_all');
```
