## Appendix S4 for "Spatial patterns from a dispersal limitation perspective: revealing biotic interactions": Compute_survival_curve.m

```
%%%%%%%%%%%%%%%%%%%%%%%%%%%%%%%%%%%%%%%%%%%%%%%%%%%%%%%%%%%%
%%% Analyzing the survival at different distances
%%%%%%%%%%%%%%%%%%%%%%%%%%%%%%%%%%%%%%%%%%%%%%%%%%%%%%%%%%%%


% each row of the matrix is a parameter regime with order:
% [dispersal distance , CNDD distance , CNDD magnitude , GNDD
% magnitude]

function [surv_r] = Compute_survival_curve(dat,ranges, num_seeds, inp)

% Sample input:
%regs_use = [1 1 4 4];% ; 1 3 4 1 ; 3 1 4 1 ; 3 3 4 1 ; 1 1 1 4 ; 3 3 1 4];
%num_seeds = 1000;
%ranges = 0.5:0.5:30;
% load('inp_all.mat')
% inp = inp_all(regs_use(1),regs_use(2), regs_use(3),regs_use(4));
% dat = load(['ext1_results/' inp.output_file '.mat']);

%% Precalculations (many of them resembling the simulation):

% For analysis:
num_ranges = length(ranges);
surv_r = zeros(1,num_ranges);

%Compute CNDDs:
if inp.Q_CNDD_min == inp.Q_CNDD_max %no variability...
    JCs = inp.Q_CNDD_min*ones(1,inp.S_reg);
else
    inc = (inp.Q_CNDD_max-inp.Q_CNDD_min)/(inp.S_reg-1); %increment
    JCs = ( inp.Q_CNDD_min : inc : inp.Q_CNDD_max ); %first species has minimal JC, last has maximal, evenly distributed.    
end

%precalculating distance and contributions to NCIs:
dists_discretized = single([0.01 0.1:0.1:(1.5*inp.L)]);
gndd = single(inp.Q_gen./(1+(1/inp.G_dist^inp.G_steep)*dists_discretized.^inp.G_steep)); %magnitude of gndd at the discretized distances
cndd = single(1./(1+(1/inp.CNDD_dist^inp.CNDD_steep)*dists_discretized.^inp.CNDD_steep)); % same for CNDD

%% %disperse the seeds at the different ranges:

year_now = unidrnd(size(dat.com_samp,3),num_seeds, num_ranges); %choose which years to sample

for rr = 1:num_ranges
    for ss = 1:num_seeds
        
        %disperse the seed:
        seed = dat.com_samp(randi(inp.J),1:3,year_now(ss,rr));
        theta = rand*2*pi; %direction
        seed(1) = seed(1) + ranges(rr)*cos(theta); %perform seed displacement - x coord.
        seed(2) = seed(2) + ranges(rr)*sin(theta); %perform seed displacement - y coord.

        % Enforce torus boundaries:
        seed(1:2) = rem(seed(1:2),inp.L); %for coordinates > L - performs wrapping, also shortens negative values to be > -L
        seed(seed<0) = inp.L + seed(seed<0); %for negative values

        %compute competitive effect of neighborhood:

        xdiff = abs(seed(1) - dat.com_samp(:,1,year_now(ss,rr))); %distance of seed to every tree along x axis
        ydiff = abs(seed(2) - dat.com_samp(:,2,year_now(ss,rr)));
        xdiff = min(xdiff, inp.L - xdiff); % TORUS
        ydiff = min(ydiff, inp.L - ydiff); % TORUS

        %now find trees neighbordhood ('quad_nei') and compute the distances to them and the NCI:

        quad_nei = (xdiff<inp.D) & (ydiff<inp.D);
        dists = sqrt(xdiff(quad_nei).^2 + ydiff(quad_nei).^2); %seed-tree distnces

        is_consp_here = (dat.com_samp(quad_nei,3,year_now(ss,rr)) == seed(3))'; %conspecific trees AMONG NEIGHBORS
        prealoc_ind = 1+round(10*dists); %find the indexes of the distances in the preallocated tables
        NCI = sum( gndd(prealoc_ind) + (JCs(seed(3)).*is_consp_here).*cndd(prealoc_ind), 2); %sum neighborhood crowding

        % add survival probability:
        surv_r(rr) = surv_r(rr) + (1/(1+NCI));
    end
end

surv_r = surv_r/num_seeds;

end
```
