## Appendix S4 for "Spatial patterns from a dispersal limitation perspective: revealing biotic interactions": compose_pieces.m

```
% compose the pieces of the simulations that haven't ended

%% Find all directories that have to be analyzed:

abd_bin_edges = [1 5:2:51 55:5:100 110:10:200 250:50:600 700:100:1000 1250 1500 5500];
dist_bin_edges = 0:10:200;

%assumption: if it's a folder it hasn't been deleted because the simulation hasn't finished running
D = dir;
D = D(3:end);

incomplete = find([D.isdir]); %indexes of incomplete simulations

parfor fff = 1:length(incomplete)
    ff = incomplete(fff);
    
    reg_now = D(ff).name;
    D2 = dir([reg_now '/']);
    
    if length(D2)>2 %there are any samples
        D2 = D2(3:end);
        com_samp = nan(5500,3,length(D2),'single'); %the composed community samples
        for ss = 1:length(D2)
            l = load([reg_now '/s_' num2str(ss) '_' reg_now '.mat']);
            com_samp(:,:,ss) = l.com;
        end
        save_com_samp(com_samp, [reg_now '.mat']);
        %save([reg_now '.mat'],'com_samp');
        [~, ~, ~, ~, ~, ~, ~] = Stats_regime6(com_samp, abd_bin_edges, dist_bin_edges, 0.25, 5, 600, 20, [reg_now '_analysis_results.mat']); %run analysis for this regime and save result separately

    end
end

%%
% for ss = 1:length(partial_sims)
%     ss
%     D = dir(['run_results/' partial_sims{ss}]); %content of the folder
%     D = D(3:end);
%     
%     com_samp = nan(5500,3,length(D),'single'); %the composed community samples
%     for dd = 1:length(D) %every sample
%         l = load(['run_results/' partial_sims{ss} '/s_' num2str(dd) '_' partial_sims{ss} '.mat']);
%         com_samp(:,:,dd) = l.com_s;
%     end
%     
%     save(['run_results/' partial_sims{ss} '.mat'],'com_samp');
% end

function save_com_samp(com_samp, filename)
save(filename,'com_samp');
end
```
