## Appendix S4 for "Spatial patterns from a dispersal limitation perspective: revealing biotic interactions": Extensive_run_analyze.m

```
 %%%%%%%%%%%%%%%%%%%%%%%%%%%%%%%%%%%%%%%%%%%%%%%%%%%%%%%%%%%%%%%%%%%
%%% Run simulations analyzing how 4 parameters affect spatial patterns
%%%%%%%%%%%%%%%%%%%%%%%%%%%%%%%%%%%%%%%%%%%%%%%%%%%%%%%%%%%%%%%%%%%
%
% Author: Michael Kalyuzhny
%
% Date created: 11/09/2020
% Date last modified: 27/01/2022
%
%%%%%%%%%%%%%%%%%%%%%%%%%%%%%%%%%%%%%%%%%%%%%%%%%%%%%%%%%%%%%%%%%%%
% Description:
%
% We explore the effect of four four levels of four parameters on NND and relative mark-correlation function:
% 1. Dispersal distance (7 m., 20 m., 60 m., 3000 m.)
% 2. CNDD distance (7 m., 20 m., 60 m., 3000 m.)
% 3. CNDD magnitude (0, 0.2, 1, 5) - Conspecific Negative Density Dependece
% 4. GNDD magnitude (0, 0.2, 1, 5) - General Negative Density Dependence
%
% These are the order of the 4 dimensions of the results
%
%% Run simulations 

% Generic regime:

inp_all = struct;
inp_all.J = 5500; %number of individuals in the forest
inp_all.L = 600; %landscape edge (meters)
inp_all.S_reg = 300; %number of species in regional pool
inp_all.b = 2; %'b' parameter of the 2DT kernel (sensu Nathan 2012), = p+1 (sensu Clark 1999), representing heavy tailedness
inp_all.a = 12.73; %second parameter of the 2DT
inp_all.D = 600; %quadrat edge
inp_all.Q_gen = 5; %magnitude of broad spectrum competitive effect
inp_all.Q_CNDD_min = 5; %minimal value of CNDD
inp_all.Q_CNDD_max = 5;
inp_all.CNDD_dist = 2000; %spatial scale at which CNDD drops
inp_all.CNDD_steep = 50; %steepness of CNDD drop with distance
inp_all.G_dist = 4.5; % GNDD scale and steepness
inp_all.G_steep = 5; % GNDD scale and steepness  
inp_all.alpha = 1; %total competitive strength - to transform NCI to seed establishment. 1/alpha = NCI leading to survival = 0.5
inp_all.imm_prob = 0.0005; %number of immigrating seeds to add

% Time and sampling parameters (time in sweeps):
inp_all.samps_tot = 3000; %how many samples to take?
inp_all.samp_freq = 10; %how many sweeps between samples?
inp_all.first_samp = 3000; %after how many sweeps to take first sample?
inp_all.print_freq = 1000;
inp_all.output_file = 'nan';

% Set levels of parameters:

range_vals = [7 20 60 3000]; %dispersal and CNDD distance values 
magnitude_vals = [0 0.2 1 5]; %CNDD and GNDD values
CNDD_steep_vals = [5 6 7 10]; %steepness
D_vals = [15 40 120 600]; %size of qudrat in which the neighborhood is computed

% Assign the values along the four dimensions

inp_all = repmat(inp_all,4,4,4,4);
for dd=1:4 %dispersal distance
    for jd = 1:4 % CNDD distance
        for jm = 1:4 %CNDD magnitude
            for hm = 1:4 %GNDD magnitude
                inp_all(dd,jd,jm,hm).a = range_vals(dd)*2*gamma(1)/(sqrt(pi)*gamma(0.5)); %set disperdal distance, formula derived from Nathan et al 2012
                inp_all(dd,jd,jm,hm).CNDD_dist = range_vals(jd);
                inp_all(dd,jd,jm,hm).CNDD_steep = CNDD_steep_vals(jd);
                inp_all(dd,jd,jm,hm).Q_CNDD_min = magnitude_vals(jm);
                inp_all(dd,jd,jm,hm).Q_CNDD_max = magnitude_vals(jm);
                inp_all(dd,jd,jm,hm).Q_gen = magnitude_vals(hm);
                inp_all(dd,jd,jm,hm).D = D_vals(jd);
                
                inp_all(dd,jd,jm,hm).output_file = ['ext1_dd_' sprintf('%0.4g',range_vals(dd)) '_jd_' sprintf('%0.4g',range_vals(jd)) '_jm_' sprintf('%0.4g',magnitude_vals(jm)) '_hm_' sprintf('%0.4g',magnitude_vals(hm))];
            end
        end
    end
end

% Set analysis parameters:
abd_bin_edges = [1 5:2:51 55:5:100 110:10:200 250:50:600 700:100:1000 1250 1500 5500];
dist_bin_edges = 10:10:200;

save('../inp_all.mat', 'inp_all')

%% Set regime:
regime_now = str2num(getenv('SLURM_ARRAY_TASK_ID'));
%% Run:

inp = inp_all(regime_now)
tmp = rand(1,regime_now); %this is meant to create some disparity in the seed of simulations beginning exactly at the same time

[com_samp, ~, ~] = sim2_1(inp);

[~,~,~, ~, ~, ~, ~, ~, ~] = Stats_regime6(com_samp, abd_bin_edges, dist_bin_edges, 0.25, 5, inp.L, 20, [inp.output_file '_analysis_results.mat']); %run analysis for this regime and save result separately
```
