## Appendix S4 for "Spatial patterns from a dispersal limitation perspective: revealing biotic interactions": sim2_1.m

```
%%%%%%%%%%%%%%%%%%%%%%%%%%%%%%%%%%%%%%%%%%%%%%%
% Forest simulator version 2.1 - unparallelized version
%%%%%%%%%%%%%%%%%%%%%%%%%%%%%%%%%%%%%%%%%%%%%%%
%
% Version comments: specialist ASYMMETRIC JC with each tree and seed in a quadrat.
% Time is continous. in this version time is printed. Torus Boundary conditions
%
% Author: Michael Kalyuzhny
%
% First written: 02/06/2020
% Last updated: 30/12/2020
%
%% Description
%  Main community matrix - 3 columns: x,y,species ID. Seeds are generated
% (with same attributes) and dispersed with torus BC. immigrants are
%  added###. 
%
%
% All time is in sweeps. Fixed number of samples is taken, after equilibration. 

function [com_samp, time_samp, runtime] = sim2_1(inp)

% %% Input patameters - example
% 
% inp.J = 40000; %number of individuals in the forest
% inp.L = 1000; %landscape edge (meters)
% inp.S_reg = 300; %number of species in regional pool
% inp.b = 2; %'b' parameter of the 2DT kernel (sensu Nathan 2012), = p+1 (sensu Clark 1999), representing heavy tailedness
% inp.a = 25; %second parameter of the 2DT
% inp.D = 40; %quadrat edge
% inp.Q_gen = 1; %magnitude of broad spectrum competitive effect
% inp.Q_CNDD_min = 0.1; %minimal value of JC
% inp.Q_CNDD_max = 0.2;
% inp.CNDD_dist = 15; %spatial scale at which JC drops
% inp.CNDD_steep = 7; %steepness of JC drop with distance
% inp.H_dist = 3;
% inp.H_steep = 6;    
% inp.alpha = 1; %total competitive strength - to transform NCI to seed establishment. 1/alpha = NCI leading to survival = 0.5
% inp.imm_prob = 0.01; %number of immigrating seeds to add
% 
% % Time and sampling parameters (time in sweeps):
% inp.samps_tot = 1; %how many samples to take?
% inp.samp_freq = 1; %how many sweeps between samples?
% inp.first_samp = 1; %after how many sweeps to take first sample?
% inp.print_freq = 1;
% inp.output_file = 'sim2.mat';

%% Initialization

%all time is in sweeps (J events)

%Compute CNDDs:
if inp.Q_CNDD_min == inp.Q_CNDD_max %no variability...
    JCs = inp.Q_CNDD_min*ones(1,inp.S_reg);
else
    inc = (inp.Q_CNDD_max-inp.Q_CNDD_min)/(inp.S_reg-1); %increment
    JCs = ( inp.Q_CNDD_min : inc : inp.Q_CNDD_max ); %first species has minimal JC, last has maximal, evenly distributed.    
end

%precalculating distance and contributions to NCIs:
dists_discretized = single([0.01 0.1:0.1:(1.5*inp.L)]);
gndd = single(inp.Q_gen./(1+(1/inp.G_dist^inp.G_steep)*dists_discretized.^inp.G_steep)); %magnitude of gndd at the discretized distances
cndd = single(1./(1+(1/inp.CNDD_dist^inp.CNDD_steep)*dists_discretized.^inp.CNDD_steep)); % same for CNDD

% Time initialization
t_now = 0;
next_samp_time = inp.first_samp;
time_samp = NaN(1,inp.samps_tot,'single'); %when were the samples taken?
next_samp_ind = 1; %index of next sample
stop_samp_ind = inp.samps_tot + 1; %when this is the next sample - stop the run
next_print = inp.print_freq;
mkdir(inp.output_file) % create a temporary directory for the samples
time_add = 1/inp.J; %the amount of time (in sweeps) that = a time step


% Generate initial community
com = zeros(inp.J,3,'single'); %this is the community composition, main dynamic variable. first two columns are coordinates. 3rd is species ID, 4th is quadrat (using linear indexing)
com(:,1:2) = rand(inp.J,2,'single')*inp.L; %set uniform random coordinates
com(:,3) = randi(inp.S_reg,inp.J,1,'single'); %draw species from uniform dist.

%% Main Loop:
tic

while next_samp_ind < stop_samp_ind

    %generate seed:
    if rand > inp.imm_prob % No immigration: generate and disperse seed:
        seed = com(randi(inp.J),1:3);
        u = rand; %temporary variable, to be transformed into distance:
        r = sqrt(((1-u)./(inp.a.^(2*inp.b-2))).^(1./(1-inp.b)) - inp.a^2 ); %transform to 2DT distance distribution
        theta = rand*2*pi; %direction
        seed(1) = seed(1) + r*cos(theta); %perform seed displacement - x coord.
        seed(2) = seed(2) + r*sin(theta); %perform seed displacement - y coord.

        % Enforce torus boundaries:
        seed(1:2) = rem(seed(1:2),inp.L); %for coordinates > L - performs wrapping, also shortens negative values to be > -L
        seed(seed<0) = inp.L + seed(seed<0); %for negative values
    else % Immigration:
        seed = [rand(1,2,'single')*inp.L randi(inp.S_reg,'single')]; %generate immigrating seeds
    end
    
    %compute competitive effect of neighborhood:
    
    xdiff = abs(seed(1) - com(:,1)); %distance of seed to every tree along x axis
    ydiff = abs(seed(2) - com(:,2));
    xdiff = min(xdiff, inp.L - xdiff); % TORUS
    ydiff = min(ydiff, inp.L - ydiff); % TORUS
    
    %now find trees neighbordhood ('quad_nei') and compute the distances to them and the NCI:
    
    quad_nei = (xdiff<inp.D) & (ydiff<inp.D);
    dists = sqrt(xdiff(quad_nei).^2 + ydiff(quad_nei).^2); %seed-tree distnces

    is_consp_here = (com(quad_nei,3) == seed(3))'; %conspecific trees AMONG NEIGHBORS
    prealoc_ind = 1+round(10*dists); %find the indexes of the distances in the preallocated tables
    NCI = sum( gndd(prealoc_ind) + (JCs(seed(3)).*is_consp_here).*cndd(prealoc_ind), 2); %sum neighborhood crowding
  
    %Check if establishment is sucessful, if so - do all the things, otherwise - skip to next itteration (do nothing):
    if rand < (1/(1 + inp.alpha*NCI))
        com(randi(inp.J,'single'),:) = seed; % replace a random tree with the seed

        %print time:
        if t_now >= next_print
            disp(['Regime: ' inp.output_file ', Runtime: ' sprintf('%0.8g',toc) ' sec., Time: ' sprintf('%0.8g',t_now) ' gen.'])
            next_print = next_print + inp.print_freq;
        end

        if t_now >= next_samp_time %if it's time to sample
            time_samp(next_samp_ind) = t_now; %record time

            save([inp.output_file '/s_' num2str(next_samp_ind) '_' inp.output_file '.mat'],'com');

            %next sampling:
            next_samp_time = next_samp_time + inp.samp_freq;
            next_samp_ind = next_samp_ind + 1;
        end

        t_now = t_now + time_add; %progress time
    end

end

runtime = toc;

% Compose the pieces and save:

com_samp = NaN(inp.J,3,inp.samps_tot,'single');

for ss = 1:inp.samps_tot
    l = load([inp.output_file '/s_' num2str(ss) '_' inp.output_file '.mat'],'com');
    com_samp(:,:,ss) = l.com;
end

save([inp.output_file '.mat'])
rmdir(inp.output_file, 's') % delete temporary directory for the samples


disp(['Finished! Runtime: ' sprintf('%0.8g',toc) ' sec., Time: ' sprintf('%0.8g',t_now) ' gen.'])
end
```
