## Appendix S4 for "Spatial patterns from a dispersal limitation perspective: revealing biotic interactions": Full_analysis.m

```
%%%%%%%%%%%%%%%%%%%%%%%%%%%%%%%%%%%%%%%%%%%%%%%%%%%%%%%%%%%%%%%%%%%%%%%%%%%
% RUN (MOST OF THE) FULL ANALYSIS
%%%%%%%%%%%%%%%%%%%%%%%%%%%%%%%%%%%%%%%%%%%%%%%%%%%%%%%%%%%%%%%%%%%%%%%%%%%
%
% Date created: 27/01/2022
% Date last modified: 29/01/2022
%
% Author: Michael Kalyuzhny
%
% Prerequisite: run the batch files calling the simulations in the folders
% 'ext1_results' and 'ext3_results'.
%
%% Finalize nescesaty computations:

Null_run_analyze; %generate the null expectations
Analyze_var_CNDD2; %perform the combined analysis of variable CNDD

%% Generate figures:

plot_kernels2; %SI Fig. 1
generate_fig1_2; %Fig 1
generate_fig_EA_3; %Fig 2
generate_fig_DD_magnitude4; %Fig 3 and SI figs.
generate_fig_var_Qc_ver4.m; %Fig 4
generate_fig_sensitivity; % Fig 5
generate_fig_all_regs; % Figures S2-5, showing the four statistics with ALL parameters
generate_fig_abd3; % effect of abudnance on spatial patterns
generate_fig_DD_distance1; %effect of CNDD and dispersal distances on spatial patterns
```
