## Appendix S4 for "Spatial patterns from a dispersal limitation perspective: revealing biotic interactions": generate_fig1_2.m

```
%%%%%%%%%%%%%%%%%%%%%%%%%%%%%%%%%%%%%%%%%%%%%%%%%%%%%%%%%%%
% GENERATE FIG. 1 EXAMPLIFYING THE NULL MODELS AND ANALYSIS
%%%%%%%%%%%%%%%%%%%%%%%%%%%%%%%%%%%%%%%%%%%%%%%%%%%%%%%%%%%

% Author: Michael Kalyuzhny
% Created: 13/07/21
% Last modified: 24/01/22

% THIS SCRIPT REQUIRES THE RAW SIMULATION RESULTS FILES:
% 'ext1_results/ext1_dd_20_jd_7_jm_5_hm_0.mat'
% 'imm_range_20_neut.mat'
% The original import is commented out. Instead, the precalculated data is
% loaded
%% Set samples and import:

dd = 2; %level of dispersal distance
jd = 1; %level of CNDD distance
jm = 4; %level of CNDD magnitude
gm = 1; %level of GNDD magnitude
L = 600; %landscape size

abd_study = 50; %what abundance should we focus on?
sim_take = 1; %which sample (with abundance 'abd_study') should be taken from the community simulation?
null_take = 7; %which sample (with abundance 'abd_study') should be taken from the null simulation?

dists = {'7' '20' '60' '3000'}; %the distances used (in m.)

% %%%%%%%%%%%%%%%%%%%%%%%%%%%% ORIGINAL CODE: #######
% %% Load all data:
% %this takes a little time
% 
% load('inp_all.mat'); %the list of all regimes
% 
% %find appropriate sample:
% l = load(['ext1_results/' inp_all(dd,jd,jm,gm).output_file '.mat']); %load simulation data
% com_samp_sim = sum_pop(l.com_samp); %change to year by abudnance matrix
% 
% null = load(['imm_range_' dists{dd} '_neut.mat']); %null
% com_samp_null = sum_pop(null.com_samp); %change to year by abudnance matrix
% 
% %% get sample from simulation and CSR:
% [y,s] = find(com_samp_sim == abd_study); %find all cases when abundance = abd_study
% 
% %extract point pattern of appropriate sample:
% points_inds = l.com_samp(:,3,y(sim_take)) == s(sim_take);
% points_sim = l.com_samp(points_inds,1:2,y(sim_take));
% 
% % Point pattern under CSR:
% points_CSR = L*rand(abd_study,2);
% 
% %% Get sample from DL null::
% 
% [y,s] = find(com_samp_null == abd_study); %find all cases when abundance = abd_study
% 
% %extract point pattern of appropriate sample:
% points_null_inds = null.com_samp(:,3,y(null_take)) == s(null_take);
% points_null = null.com_samp(points_null_inds,1:2,y(null_take));
% 
% save('points_Fig1_precalculated.mat','points_sim', 'points_CSR', 'points_null')
% %%%%%%%%%%%%%%%%%%%%%%%%%%%%%%%%%%%%%%%%%%%%%%%%%%%%%%%%%%%%%%%%%%%%%%%

%% Import pre-calulated data:

load('points_Fig1_precalculated.mat','points_sim', 'points_CSR', 'points_null')
%% Perform analyses for the histograms:

[dists_sim, neighbors20_sim] = analysis1(points_sim, L);
[dists_CSR, neighbors20_CSR] = analysis1(points_CSR, L);
[dists_null, neighbors20_null] = analysis1(points_null, L);

%smooth them:
[f_dists_sim,x_dists_sim] = ksdensity(dists_sim ,'Support','positive','BoundaryCorrection','reflection');
[f_dists_CSR,x_dists_CSR] = ksdensity(dists_CSR,'Support','positive','BoundaryCorrection','reflection');
[f_dists_null,x_dists_null] = ksdensity(dists_null,'Support','positive','BoundaryCorrection','reflection');

%count neighbors:
max_neighbors = max([neighbors20_sim neighbors20_CSR neighbors20_null]);
neighbors_bins_sim = histcounts(neighbors20_sim, (0:(max_neighbors+1))-0.01);
neighbors_bins_CSR = histcounts(neighbors20_CSR, (0:(max_neighbors+1))-0.01);
neighbors_bins_null = histcounts(neighbors20_null, (0:(max_neighbors+1))-0.01);

%% Plot:

f = figure();

% Plot simulation

s1 = subplot(2,3,1);
hold on
set(s1,'FontSize',16,'FontWeight','bold','LineWidth',1.5);

scatter(points_sim(:,1), points_sim(:,2), 24, ...
    'MarkerEdgeColor',[0 0 0], 'MarkerFaceColor',[0.1 .9 0], 'MarkerFaceAlpha', .5)
xlim([0 600])
ylim([0 600])
xticks([0 200 400 600])
yticks([0 200 400 600])
text(10,570,'(a)','FontSize',16,'FontWeight','bold')
title('Population')
ylabel('Y (m.)')

% Plot CSR

s1 = subplot(2,3,2);
hold on
set(s1,'FontSize',16,'FontWeight','bold','LineWidth',1.5);

scatter(points_CSR(:,1), points_CSR(:,2), 24, ...
    'MarkerEdgeColor',[0 0 0], 'MarkerFaceColor',[1 0 0], 'MarkerFaceAlpha', .5)
xlim([0 600])
ylim([0 600])
xticks([0 200 400 600])
yticks([0 200 400 600])
text(10,570,'(b)','FontSize',16,'FontWeight','bold')
title('Random distribution')
xlabel('X (m.)')

% Plot DL_null

s1 = subplot(2,3,3);
hold on
set(s1,'FontSize',16,'FontWeight','bold','LineWidth',1.5);

scatter(points_null(:,1), points_null(:,2), 24, ...
    'MarkerEdgeColor',[0 0 0], 'MarkerFaceColor',[0 .1 .9], 'MarkerFaceAlpha', .5)
xlim([0 600])
ylim([0 600])
xticks([0 200 400 600])
yticks([0 200 400 600])
text(10,570,'(c)','FontSize',16,'FontWeight','bold')
title('Dispersal Limitation')

% plot distribution of NND:

s1 = subplot(2,3,4.5);
hold on
set(s1,'FontSize',16,'FontWeight','bold','LineWidth',1.5);

plot(x_dists_sim, f_dists_sim, 'LineWidth',1.5, 'Color', [0.1 .9 0])
plot(x_dists_CSR, f_dists_CSR, 'LineWidth',1.5, 'Color', [1 0 0])
plot(x_dists_null, f_dists_null, 'LineWidth',1.5, 'Color', [0 .1 .9])
xlim([0 120])
xlabel('Nearest neighbor distance (m.)')
ylabel('Frequency')
text(5,0.076,'(d)','FontSize',16,'FontWeight','bold')

% Plot number of neighbors:

s1 = subplot(2,3,5.5);
hold on
set(s1,'FontSize',16,'FontWeight','bold','LineWidth',1.5);
b = bar([neighbors_bins_CSR; neighbors_bins_sim; neighbors_bins_null]','BaseValue',-.25, 'FaceColor','flat');
b(1).CData = [1 0 0];
b(2).CData = [0.1 .9 0];
b(3).CData = [0 .1 .9];
ylim([0 50]);
xlabel('Neighbors withing 20 m.')
xticks(1:(max_neighbors+1))
xticklabels(string(0:max_neighbors))
text(0.6,47,'(e)','FontSize',16,'FontWeight','bold')


%% Export:

set(f,'PaperSize',[12 8]); %set the paper size to what you want
print(f,'Fig1','-dpdf')

%%

function [dists, neighbors20] = analysis1(pp, L)
%recives the point pattern pp and returns the distributions of nearest
%neighbor distances and the number of neighbors within 20 meters

this_abd = size(pp,1);    
xdists = abs(pp(:,1)-pp(:,1)'); %distance between every pair of points on a torus along x axis
xdists = min(xdists,L-xdists); %torus!
ydists = abs(pp(:,2)-pp(:,2)'); %distance between every pair of points on a torus along y axis
ydists = min(ydists,L-ydists); %torus!
dd = sqrt(xdists.^2+ydists.^2); %distances of first 'num_samples' individuals to all 

%replace diagonal with Inf (so that your distance to yourseld does not get counted in any distance bin):
dd(1:(this_abd+1):(this_abd^2)) = Inf;

dists = min(dd,[],2); %find minimum in every row:
neighbors20 = sum(dd<20);
    
end
```
