## Appendix S4 for "Spatial patterns from a dispersal limitation perspective: revealing biotic interactions": generate_fig_abd3.m

```
%%%%%%%%%%%%%%%%%%%%%%%%%%%%%%%%%%%%%%%%%%%%%%%%%%%%%%%%%%%%%%%%%%%%
%%% Generate Figures of the effect of abundance 
%%%%%%%%%%%%%%%%%%%%%%%%%%%%%%%%%%%%%%%%%%%%%%%%%%%%%%%%%%%%%%%%%%%%

% Author: Michael Kalyuzhny

% First creted: 09/08/21
% Last updated: 09/08/21

%% Set up:

% each row of the matrix is a parameter regime with order:
% [dispersal distance , CNDD distance , CNDD magnitude , GNDD
% magnitude]

regs = [1 1 1 1 ; 3 1 1 1 ; 4 1 1 1 ; 1 1 3 1 ; 1 1 4 1 ; 3 1 4 1 ; 1 3 4 1 ; 3 3 4 1];
min_sample_size = 100;

% import_null:
refs = cell(1,4);
dists = {'7' '20' '60' '3000'}; %the distances used (in m.)
for dd = 1:length(dists)    
    refs{dd} = load(['imm_range_' dists{dd} '_neut_analysis_results.mat']); %null
end

% Load data and calculate ED and EA:
load('inp_all.mat');
dat = cell(1,size(regs,1));

for rr = 1:size(regs,1)

inp = inp_all(regs(rr,1), regs(rr,2), regs(rr,3), regs(rr,4));
dat{rr} = load(['ext1_results/' inp.output_file '_analysis_results.mat']);
dat{rr}.ED = dat{rr}.mean_log_dist - refs{regs(rr,1)}.mean_log_dist;
dat{rr}.EA = dat{rr}.log_N20 - refs{regs(rr,1)}.log_N20;

% Delete cases when sample size is too small:
too_few_samps = dat{rr}.samps_per_bin < min_sample_size;

dat{rr}.ED(too_few_samps) = nan;
dat{rr}.EA(too_few_samps) = nan;
dat{rr}.CE(too_few_samps) = nan;
dat{rr}.RND20(too_few_samps) = nan;
dat{rr}.N20_p90(too_few_samps) = nan;

end

%% Plot Statistics:

f = figure()

% [dispersal distance , CNDD distance , CNDD magnitude , GNDD
% magnitude]
%[1 1 1 1 ; 3 1 1 1 ; ##4 1 1 1## ; 1 1 3 1 ; 1 1 4 1 ; 3 1 4 1 ; 1 3 4 1 ; 3 3 4 1];
% thin line: Q_C = 0, intermediate = 1, thick = 5
% solid: delta_C = 7 m., dash: delta_C = 60 m.
% blue: dispreal = 60 m. red: dierpsal = 7 m.
legend_entries_regs = {'Q_C=0, D=7 m.', ...
    'Q_C=0, D=60 m.', ...
    'Q_C=0, D=\infty (CSR)', ...
    'Q_C=1, \Delta_C=7 m., D=7 m.', ...
    'Q_C=5, \Delta_C=7 m., D=7 m.', ...
    'Q_C=5, \Delta_C=7 m., D=60 m.', ...
    'Q_C=5, \Delta_C=60 m., D=7 m.', ...
    'Q_C=5, \Delta_C=60 m., D=60 m.', ...
    'Square grid'};

syms = {'-', '-', ':', '-', '-', '-', '--', '--'};
cols = [.9 .1 0 ; 0 .1 .9 ; 0 0 0 ; .9 .1 0 ; .9 .1 0 ; 0 .1 .9 ; .9 .1 0 ; 0 .1 .9];
widths = [.5 .5 1 1 2.5 2.5 2.5 2.5];

regs_use = [4 5 6 7 8];

s1 = subplot(2,3,1);
hold on
set(s1,'FontSize',16,'FontWeight','bold','LineWidth',1.5, 'XScale','log');
for rr = regs_use
    plot(dat{rr}.abd_bin_centers ,dat{rr}.EA, syms{rr}, 'color',cols(rr,:),'MarkerFaceColor',cols(rr,:),'LineWidth',widths(rr))
end

ylabel('Excess Abundance (EA(20))')
text(500,-.1,'(a)','FontSize',16,'FontWeight','bold')
xlim([5 1000])
xticks([10 100 1000])

regs_use = [1 2 4 5 6 7 8];

s1 = subplot(2,3,2);
hold on
set(s1,'FontSize',16,'FontWeight','bold','LineWidth',1.5, 'XScale','log', 'YScale','log');
for rr = regs_use
    plot(dat{rr}.abd_bin_centers ,dat{rr}.RND20, syms{rr}, 'color',cols(rr,:),'MarkerFaceColor',cols(rr,:),'LineWidth',widths(rr))
end

ylabel('Relative density (RND(20))')
xlim([5 1000])
xticks([10 100 1000])
text(500,130,'(b)','FontSize',16,'FontWeight','bold')

%% Plot explanatory patterns:

regs_use = 1:8;
%[1 1 1 1 ; 3 1 1 1 ; 4 1 1 1 ; 1 1 3 1 ; 1 1 4 1 ; 3 1 4 1 ; 1 3 4 1 ; 3 3 4 1];

s1 = subplot(2,3,4);
hold on
set(s1,'FontSize',16,'FontWeight','bold','LineWidth',1.5, 'XScale','log');
for rr = regs_use
    plot(dat{rr}.abd_bin_centers ,dat{rr}.mean_log_dist, syms{rr}, 'color',cols(rr,:),'MarkerFaceColor',cols(rr,:),'LineWidth',widths(rr));
end
plot(dat{rr}.abd_bin_centers, log(inp.L./sqrt(dat{rr}.abd_bin_centers)), 'color',[.88 .05 .92]','LineWidth',1.5) 
xlim([5 1000])
xticks([10 100 1000])
legend(legend_entries_regs)
ylabel('Log(nearest neighbor dist. (m))')
text(500, 5.8,'(c)','FontSize',16,'FontWeight','bold')

regs_use = [1 3 4 5 7];

s1 = subplot(2,3,5);
hold on
set(s1,'FontSize',16,'FontWeight','bold','LineWidth',1.5, 'XScale', 'log');
for rr = regs_use
    plot(dat{rr}.abd_bin_centers ,dat{rr}.occupancy, syms{rr}, 'color',cols(rr,:),'MarkerFaceColor',cols(rr,:),'LineWidth',widths(rr));
end
xlim([5 1000])
xticks([10 100 1000])
xlabel('Abundance')
ylabel('20X20 m. cells occupied')
text(500, .77,'(d)','FontSize',16,'FontWeight','bold')

s1 = subplot(2,3,6);
hold on
set(s1,'FontSize',16,'FontWeight','bold','LineWidth',1.5, 'XScale','log');
for rr = regs_use
    plot(dat{rr}.abd_bin_centers ,dat{rr}.N20_p90, syms{rr}, 'color',cols(rr,:),'MarkerFaceColor',cols(rr,:),'LineWidth',widths(rr));
end
xlim([5 1000])
xticks([10 100 1000])
annotation('doublearrow',[.15 .15],[.15 .25],'LineWidth',2.5,'color',[.2 .2 .2])
annotation('doublearrow',[.18 .18],[.15 .3],'LineWidth',2.5,'color',[.2 .2 .2])
annotation('textbox','color',[.2 .2 .2],'String','Maximal overdispersion','FontSize',14,'LineStyle','none','FontWeight','bold')
ylabel('90th percentile of conspecific neighbors')
text(500, .96,'(e)','FontSize',16,'FontWeight','bold')

%% Export

set(f,'PaperSize',[13 9]); %set the paper size to what you want
print(f,'Fig_abd_v3','-dpdf')

%% Generate SI figuere with additional statistics:

f = figure()

regs_use = [4 5 6 7 8];

s1 = subplot(1,3,1);
hold on
set(s1,'FontSize',16,'FontWeight','bold','LineWidth',1.5, 'XScale','log');
for rr = regs_use
    plot(dat{rr}.abd_bin_centers ,dat{rr}.ED, syms{rr}, 'color',cols(rr,:),'MarkerFaceColor',cols(rr,:),'LineWidth',widths(rr))
end

ylabel('Excess Distance (ED)')
text(500,.96,'(a)','FontSize',16,'FontWeight','bold')
xlim([5 1000])
xticks([10 100 1000])

regs_use = [1 2 4 5 6 7 8];

s1 = subplot(1,3,2);
hold on
set(s1,'FontSize',16,'FontWeight','bold','LineWidth',1.5, 'XScale','log', 'YScale','log');
for rr = regs_use
    plot(dat{rr}.abd_bin_centers ,dat{rr}.CE, syms{rr}, 'color',cols(rr,:),'MarkerFaceColor',cols(rr,:),'LineWidth',widths(rr))
end

ylabel('Clark-Evans')
xlim([5 1000])
xticks([10 100 1000])
text(500,.95,'(b)','FontSize',16,'FontWeight','bold')
xlabel('Abundance')
legend(legend_entries_regs)

%% Export

set(f,'PaperSize',[13 5]); %set the paper size to what you want
print(f,'Fig_abd_SI','-dpdf')

%%%%%%%%%%%%%%%%%%%%%%%%%%%%%%%%%%%%%%%%%%%%%%%%%%%%%%%%%%%%%%%%%%%%%%%%%%%%%%%%%%%%%%%%%%%%%%%%%
% %% Re-Run the needed regimes to compute extra statistics:
% 
% % each row of the matrix is a parameter regime with order:
% % [dispersal distance , CNDD distance , CNDD magnitude , GNDD
% % magnitude]
% 
% regs_abd = [1 1 1 1 ; 3 1 1 1 ; 4 1 1 1 ; 1 1 3 1 ; 1 1 4 1 ; 3 1 4 1 ; 1 3 4 1 ; 3 3 4 1];
% 
% abd_bin_edges = [1 5:2:51 55:5:100 110:10:200 250:50:600 500:100:1000 1250 1500 5500];
% dist_bin_edges = 0:10:200;
% load('../inp_all.mat')
% 
% parfor rr = 1:size(regs_abd,1)
% 
% inp = inp_all(regs_abd(rr,1), regs_abd(rr,2), regs_abd(rr,3), regs_abd(rr,4));
% tmp = load([inp.output_file '.mat'],'com_samp');
% com_samp = tmp.com_samp;
% 
% [~, ~, ~, ~, ~, ~, ~] = Stats_regime4(com_samp, abd_bin_edges, dist_bin_edges, 0.25, 5, inp.L, 20, [inp.output_file '_analysis_results.mat']); %run analysis for this regime and save result separately
% 
% end
```
