## Appendix S4 for "Spatial patterns from a dispersal limitation perspective: revealing biotic interactions": generate_fig_all_regs.m

```
%%%%%%%%%%%%%%%%%%%%%%%%%%%%%%%%%%%%%%%%%%%%%%%%%%%%%%%%%%%%%%%%%%
% Generate the figures of all 256 combinations of parameters
%%%%%%%%%%%%%%%%%%%%%%%%%%%%%%%%%%%%%%%%%%%%%%%%%%%%%%%%%%%%%%%%%%

% Author: Michael Kalyuhny
%
% First Written: 03/08/21
% Last modified: 28/01/22

%% Calculate all statistics:

% Regime indexing:
% [dispersal distance , CNDD distance , CNDD magnitude , GNDD
% magnitude]

CE_all = nan(4,4,4,4); %Clark-Evans for all regimes
ED_all = nan(4,4,4,4); %Excess distance for all regimes
RND20_all = nan(4,4,4,4); %relative neighborhood density at 20 meters for all regimes
EA20_all = nan(4,4,4,4); %Excess neighborhood abundance at 20 meters for all regimes

load('inp_all.mat'); %the list of all regimes
dists = {'7' '20' '60' '3000'}; %the distances used (in m.)

for dd=1:4 %dispersal distance
    ref = load(['imm_range_' dists{dd} '_neut_analysis_results.mat']); %null
    for cd = 1:4 %CNDD (Janzen-Connell) distance
        for cm = 1:4 %CNDD (Janzen-Connell) magnitude
            for gm = 1:4 %GNDD magnitude
                reg_now = load(['ext1_results/' inp_all(dd,cd,cm,gm).output_file '_analysis_results.mat']);

                %compute weighted means:
                CE_all(dd,cd,cm,gm) = nansum(reg_now.CE.*reg_now.samps_per_bin)/sum(reg_now.samps_per_bin);
                ED_all(dd,cd,cm,gm) = nansum((reg_now.mean_log_dist - ref.mean_log_dist).*reg_now.samps_per_bin)/sum(reg_now.samps_per_bin);
                EA20_all(dd,cd,cm,gm) = nansum((reg_now.log_N20 - ref.log_N20).*reg_now.samps_per_bin)/sum(reg_now.samps_per_bin);
                RND20_all(dd,cd,cm,gm) = nansum(reg_now.RND20.*reg_now.samps_per_bin)/sum(reg_now.samps_per_bin);

            end
        end
    end
end


%% Plot stats:

% RECOMMENDATION: UNCOMMENT A FIGURE EACH TIME

dists_title = {'7 m.', '20 m.', '60 m.', '\infty'};

% Plot ED:
% f1 = plot_all(ED_all, [-0.05 1], [4 1], 'Excess Distance (ED)', dists_title, false)
% set(f1,'PaperSize',[12 13]); %set the paper size to what you want
% print(f1,'Fig_SI_ED_all','-dpdf')

%f2 = plot_all(CE_all, [0 1.3], [4 1], 'Clark-Evans', dists_title, false)
% set(f2,'PaperSize',[11 13]); %set the paper size to what you want
% print('-painters',f2,'Fig_SI_CE_all','-dpdf')

% f3 = plot_all(EA20_all, [-1.6 0.05], [4 1], 'Excess neighborhood Abundance (EA(20))', dists_title, false)
% set(f3,'PaperSize',[11 13]); %set the paper size to what you want
% print('-painters',f3,'Fig_SI_EA20_all','-dpdf')

f4 = plot_all(RND20_all, [0.2 140], [4 1], 'Relative Neighborhood Density (RND(20))', dists_title, true)
set(f4,'PaperSize',[11 13]); %set the paper size to what you want
print(f4,'Fig_SI_RND20_all','-dpdf')
%% The main plotting function:

function [f] = plot_all(stat_now, ylim_now, subplot_legend, yaxis_name, dists, Y_log)

f = figure();

for dd=1:4
    for cd = 1:4
        sp = subplot(4,4,4*(dd-1)+cd);
        set(sp,'FontSize',20,'FontWeight','bold','LineWidth',1.5);
         
        st = (reshape(stat_now(dd,cd,:,:),4,4));
        
        %plot:
        col_regimes = categorical({'Q_C = 0', 'Q_C = 0.2', 'Q_C = 1', 'Q_C = 5'});
        col_regimes = reordercats(col_regimes, {'Q_C = 0', 'Q_C = 0.2', 'Q_C = 1', 'Q_C = 5'});
        h = bar((col_regimes), st);
        h(1).FaceColor = 'b';
        h(2).FaceColor = 'g';
        h(3).FaceColor = 'y';
        h(4).FaceColor = 'r';
        ylim(ylim_now)
        set(sp,'YGrid','on')
        title(['\Delta_C = ' dists{cd} ', D = '  dists{dd}])
        
        if Y_log == true
            set(sp,'YScale','log')
        end
            
        if (dd==subplot_legend(1)) && (cd==subplot_legend(2))
            legend({'Q_G = 0', 'Q_G = 0.2', 'Q_G = 1', 'Q_G = 5'})
            annotation('arrow',[.4 .6],[.98 .98],'LineWidth',2.5)
            t = text(2,2,'CNDD Distance (\Delta_C)', 'FontSize', 16', 'FontWeight', 'bold');
            annotation('arrow',[.02 .02],[.6 .4],'LineWidth',2.5)
            t = text(0,2,'Dispersal distance (D)', 'FontSize', 16', 'FontWeight', 'bold');
            set(t,'Rotation',90)
            ylabel(yaxis_name, 'FontSize', 16', 'FontWeight', 'bold')
            annotation('doublearrow',[.95 .95],[.3 .7],'LineWidth',2.5)
            t = text(6,1,'Overdispersed', 'FontSize', 16', 'FontWeight', 'bold');
            set(t,'Rotation',90)
            t = text(8,1,'Aggregated', 'FontSize', 16', 'FontWeight', 'bold');
            set(t,'Rotation',90)
        end
    end
end

end
```
