## Appendix S4 for "Spatial patterns from a dispersal limitation perspective: revealing biotic interactions": generate_fig_DD_distance1.m

```
 %%%%%%%%%%%%%%%%%%%%%%%%%%%%%%%%%%%%%%%%%%%%%%%%%%%%%%%%%%%%%%%%%%%%
% Generate figures of the effect of CNDD and dispersal distance on the spatial statistics 
%%%%%%%%%%%%%%%%%%%%%%%%%%%%%%%%%%%%%%%%%%%%%%%%%%%%%%%%%%%%%%%%%%%%

% Author: Michael Kalyuhny
%
% First Written: 01/08/21
% Last modified: 03/08/21

%%%%%%%%%%%%%%%%%%%%%%%%%%%%%%%%%%%%%%%%%%%%%%%%%%%%%%%%%%%%%%%%%%%%%%%%%%
%% PART ONE: EFFECT OF CNDD DISTANCE
%%%%%%%%%%%%%%%%%%%%%%%%%%%%%%%%%%%%%%%%%%%%%%%%%%%%%%%%%%%%%%%%%%%%%%%%%%

%% Set up parameters to use:

% each row of the matrix is a parameter regime with order:
% [dispersal distance , CNDD distance , CNDD magnitude , GNDD
% magnitude]
% here the CNDD distance parameter is written but all levels are used

regs_CNDD = [1 1 4 2 ; 2 1 4 2 ; 3 1 4 2 ; 1 1 2 2 ];
CNDD_levels = 4;

reg_num = size(regs_CNDD,1);
%% import_null:

refs = cell(1,4);
dists = {'7' '20' '60' '3000'}; %the distances used (in m.)

for dd = 1:length(dists)    
    refs{dd} = load(['imm_range_' dists{dd} '_neut_analysis_results.mat']); %null
end

%% import and analyze data for CNDD:

load('inp_all.mat')

% in the data matrices for plotting, each row is a regime and each column is a CNDD level

EA20s_CNDD = nan(reg_num, CNDD_levels);
EDs_CNDD = nan(reg_num, CNDD_levels);
CEs_CNDD = nan(reg_num, CNDD_levels);
RN20_CNDD = nan(reg_num, CNDD_levels);

for rr = 1:reg_num %run over the different regimes
    for cc = 1:CNDD_levels
        
        dat = load(['ext1_results/' inp_all(regs_CNDD(rr,1), cc, regs_CNDD(rr,3), regs_CNDD(rr,4)).output_file '_analysis_results.mat']);

        EA20s_CNDD(rr,cc) = nansum(dat.samps_per_bin.*(dat.log_N20 - refs{regs_CNDD(rr,1)}.log_N20))/nansum(dat.samps_per_bin);
        EDs_CNDD(rr,cc) = nansum(dat.samps_per_bin.*(dat.mean_log_dist - refs{regs_CNDD(rr,1)}.mean_log_dist))/nansum(dat.samps_per_bin);
        CEs_CNDD(rr,cc) = nansum(dat.samps_per_bin.*(dat.CE))/nansum(dat.samps_per_bin);
        RN20_CNDD(rr,cc) = nansum(dat.samps_per_bin.*(dat.RND20))/nansum(dat.samps_per_bin);

    end
end

%% plot ED and CE (a + b):

cols = [0 .1 .9 ; 0 .9 .1 ; .9 0 .1 ; 0 .1 .9  ];
syms = {'-o', '-o', '-o', '-x'};
legend_entries_regs = {'disp.: 7 m., Q_C = 5', 'disp.: 20 m., Q_C = 5', 'disp.: 60 m., Q_C = 5', 'disp.: 7 m., Q_C = 0.2'};

xvalues = [7 20 60 180];
xtick_text = {'7', '20', '60', '\infty'};

% o: Qc = 5 , x: 0.2.
% blue: dispersal distance = 7 m, green: 20 m.., red: inf

% [dispersal distance , CNDD distance , CNDD magnitude , GNDD magnitude]
%regs_CNDD = [1 1 4 2 ; 2 1 4 2 ; 3 1 4 2 ; 1 1 2 2 ];

figure();

s1 = subplot(4,2,3);
hold on
set(s1,'FontSize',16,'FontWeight','bold','LineWidth',1.5, 'XScale','log');
for rr = 1:reg_num
    plot(xvalues ,EDs_CNDD(rr,:),syms{rr},'color',cols(rr,:),'MarkerFaceColor',cols(rr,:),'LineWidth',1.5)
end

xticks(xvalues);
xticklabels(xtick_text)
legend(legend_entries_regs,'FontSize',12)
xlabel('CNDD distance (\Delta_C (m.))')
ylabel('Excess Distance (ED)')
text(130,0.92,'(a)','FontSize',16,'FontWeight','bold')

s1 = subplot(4,2,5);
hold on
set(s1,'FontSize',16,'FontWeight','bold','LineWidth',1.5, 'YScale','log', 'XScale','log');
for rr = 1:reg_num
    plot(xvalues ,CEs_CNDD(rr,:),syms{rr},'color',cols(rr,:),'MarkerFaceColor',cols(rr,:),'LineWidth',1.5)
end

xticks(xvalues);
xticklabels(xtick_text)
xlabel('CNDD distance (\Delta_C (m.))')
ylabel('Clark-Evans')
text(130,0.85,'(b)','FontSize',16,'FontWeight','bold')

% NEXT SECTION TAKES TIME SO IS COMMENTED OUR BY DEFAULT - use it with raw
% simulation results to compute the survival curves

% %% Compute the survival curves for several of the regimes above:
% 
% regs_CNDD_dist2 = [1 1 4 2 ; 3 1 4 2 ]; %for the sake of the survival curves
% percentiles_examine = 0.01:0.01:0.99;
% levels_CNDD_dist2 = [1 2 3 4]; %what levels of CNDD distance should be examined
% num_seeds = 500;
% 
% num_regs_CNDD_dist2 = size(regs_CNDD_dist2,1);
% num_levels_CNDD_dist2 = length(levels_CNDD_dist2);
% load('inp_all.mat')
% 
% %In all this section, rows are regimes of other parameters, columnts are levels of GNDD
% surv_curves = cell(num_regs_CNDD_dist2,num_levels_CNDD_dist2);
% dists_examine = cell(num_regs_CNDD_dist2,num_levels_CNDD_dist2); %save the dists
% 
% parfor ii = 1:(num_regs_CNDD_dist2*num_levels_CNDD_dist2)
%     [rr, cc] = ind2sub([num_regs_CNDD_dist2 num_levels_CNDD_dist2],ii); 
%     
%     %load data:
%     inp = inp_all(regs_CNDD_dist2(rr,1), levels_CNDD_dist2(cc), regs_CNDD_dist2(rr,3), regs_CNDD_dist2(rr,4));
%     dat = load(['ext1_results/' inp.output_file '.mat']);
% 
%     % Define a few functions and calculae the CDF of dispersal distances
%     dists_examine{ii} = Percentiles_to_dists(percentiles_examine, 0.1:0.1:200, inp.a, inp.b);
%     
%     surv_curves{ii} = Compute_survival_curve(dat,dists_examine{ii}, num_seeds, inp);
% end
% 
% save('CNDD_dist_reg_survival_curves.mat', 'percentiles_examine', 'regs_CNDD_dist2', 'levels_CNDD_dist2', 'num_regs_CNDD_dist2', 'num_levels_CNDD_dist2', 'num_seeds', 'dists_examine', 'surv_curves')

%% Plot the survival curves for several the regimes above:

load('CNDD_dist_reg_survival_curves.mat')

cols = [0 .1 .9 ; 0.9 0 .1 ]; %different for every regime
syms = {':', '--', '-', '-'};
lws = [1.5 1.5 1.5 2.5];

s1 = subplot(4,2,7);
hold on
set(s1,'FontSize',16,'FontWeight','bold','LineWidth',1.5, 'YScale','log');
xlabel('Percentile of dispersal distance distribution')
ylabel('Survival probability')
text(0.9,0.8,'(c)','FontSize',16,'FontWeight','bold')

for rr = 1:num_regs_CNDD_dist2

    for cc = 1:num_levels_CNDD_dist2
        
        plot(percentiles_examine, surv_curves{rr, cc}, syms{cc},'color',cols(rr,:),'LineWidth',lws(cc))

    end
end

text(0.02, 0.02, 'Disp=7 m.', 'Color',[0 .1 .9], 'FontSize',14)
text(0.32, 0.02, 'Disp=60 m.', 'Color',[0.9 0 .1], 'FontSize',14)
text(0.62, 0.017, 'Dash:  \Delta_C',  'FontSize',14)


%%%%%%%%%%%%%%%%%%%%%%%%%%%%%%%%%%%%%%%%%%%%%%%%%%%%%%%%%%%%%%%%%%%%%%%%%%
%% PART TWO: EFFECT OF Dispersal distance
%%%%%%%%%%%%%%%%%%%%%%%%%%%%%%%%%%%%%%%%%%%%%%%%%%%%%%%%%%%%%%%%%%%%%%%%%%

%% Set  up parameters to use:

% each row of the matrix is a parameter regime with order:
% [dispersal distance , CNDD distance , CNDD magnitude , GNDD
% magnitude]
% here the dispersal parameter is written but all levels are used

regs_disp = [1 1 4 2 ; 1 2 4 2 ; 1 3 4 2 ; 1 1 2 2 ];
disp_levels = 4;

reg_num = size(regs_disp,1);
%% import_null:

refs = cell(1,4);
dists = {'7' '20' '60' '3000'}; %the distances used (in m.)

for dd = 1:length(dists)    
    refs{dd} = load(['imm_range_' dists{dd} '_neut_analysis_results.mat']); %null
end

%% import and analyze data for CNDD:

load('inp_all.mat')

% in the data matrices for plotting, each row is a regime and each column is a CNDD level

EA20s_disp = nan(reg_num, disp_levels);
EDs_disp = nan(reg_num, disp_levels);
CEs_disp = nan(reg_num, disp_levels);
RND20_disp = nan(reg_num, disp_levels);

for rr = 1:reg_num %run over the different regimes
    for gg = 1:disp_levels
        
        dat = load(['ext1_results/' inp_all(gg, regs_disp(rr,2), regs_disp(rr,3), regs_disp(rr,4)).output_file '_analysis_results.mat']);

        EA20s_disp(rr,gg) = nansum(dat.samps_per_bin.*(dat.log_N20 - refs{gg}.log_N20))/nansum(dat.samps_per_bin);
        EDs_disp(rr,gg) = nansum(dat.samps_per_bin.*(dat.mean_log_dist - refs{gg}.mean_log_dist))/nansum(dat.samps_per_bin);
        CEs_disp(rr,gg) = nansum(dat.samps_per_bin.*(dat.CE))/nansum(dat.samps_per_bin);
        RND20_disp(rr,gg) = nansum(dat.samps_per_bin.*(dat.RND20))/nansum(dat.samps_per_bin);

    end
end

%% plot ED and CE (d + e):

cols = [0 .1 .9 ; 0 .9 .1 ; .9 0 .1 ; 0 .1 .9 ];
syms = {'-o', '-o', '-o', '-x'};
legend_entries_regs = {'\Delta_c = 7 m., Q_C = 5', '\Delta_c = 20 m., Q_C = 5', '\Delta_c = 60 m., Q_C = 5', '\Delta_c = 7 m., Q_C = 0.2'};
xvalues = [7 20 60 180];
xtick_text = {'7', '20', '60', '\infty'};

% o: Q_c = 5, x: Q_c = .2.
% blue: disp = 7, green: disp 20, disp: 60

% [dispersal distance , CNDD distance , CNDD magnitude , GNDD magnitude]
%regs_disp = [1 1 4 2 ; 1 2 4 2 ; 1 3 4 2 ; 1 1 2 2 ];

s1 = subplot(4,2,4);
hold on
set(s1,'FontSize',16,'FontWeight','bold','LineWidth',1.5, 'XScale','log');
for rr = 1:reg_num
    plot(xvalues ,EDs_disp(rr,:),syms{rr},'color',cols(rr,:),'MarkerFaceColor',cols(rr,:),'LineWidth',1.5)
end

xticks(xvalues);
xticklabels(xtick_text)
legend(legend_entries_regs,'FontSize',12)
xlabel('Dispersal distance (m.)')
%ylabel('Excess Distance (ED)')
%xlim([0 6])
text(130,0.9,'(d)','FontSize',16,'FontWeight','bold')

s1 = subplot(4,2,6);
hold on
set(s1,'FontSize',16,'FontWeight','bold','LineWidth',1.5, 'XScale','log');
for rr = 1:reg_num
    plot(xvalues ,CEs_disp(rr,:),syms{rr},'color',cols(rr,:),'MarkerFaceColor',cols(rr,:),'LineWidth',1.5)
end

xticks(xvalues);
xticklabels(xtick_text)
xlabel('Dispersal distance (m.)')
%ylabel('Clark-Evans')
%xlim([0 6])
text(60,1.1,'(e)','FontSize',16,'FontWeight','bold')

annotation('doublearrow',[.95 .95],[.3 .6],'LineWidth',2.5)
t = text(250,2,'Overdispersed', 'FontSize', 16', 'FontWeight', 'bold');
set(t,'Rotation',90)
t = text(250,0,'Aggregated', 'FontSize', 16', 'FontWeight', 'bold');
set(t,'Rotation',90)

% %% Compute the survival curves for several of the regimes above:
% 
% regs_disp2 = [1 1 4 2 ; 1 3 4 2 ]; %for the sake of the survival curves
% levels_CNDD_dist2 = [1 2 3 4]; %what levels of dispersal should be examined
% percentiles_examine = 0.01:0.01:0.99;
% num_seeds = 50000;
% 
% num_regs_disp2 = size(regs_disp2,1);
% num_levels_CNDD_dist2 = length(levels_CNDD_dist2);
% load('inp_all.mat')
% 
% %In all this section, rows are regimes of other parameters, columnts are levels of GNDD
% surv_curves = cell(num_regs_disp2,num_levels_CNDD_dist2);
% dists_examine = cell(num_regs_disp2,num_levels_CNDD_dist2); %save the dists
% 
% parfor ii = 1:(num_regs_disp2*num_levels_CNDD_dist2)
%     [rr, gg] = ind2sub([num_regs_disp2 num_levels_CNDD_dist2],ii); 
%     
%     %load data:
%     inp = inp_all(levels_CNDD_dist2(gg), regs_disp2(rr,2), regs_disp2(rr,3), regs_disp2(rr,4));
%     dat = load(['ext1_results/' inp.output_file '.mat']);
% 
%     % Define a few functions and calculae the CDF of dispersal distances
%     dists_examine{ii} = Percentiles_to_dists(percentiles_examine, 0.1:0.1:200, inp.a, inp.b);
%     
%     surv_curves{ii} = Compute_survival_curve(dat,dists_examine{ii}, num_seeds, inp);
% end
% 
% save('disp_reg_survival_curves.mat', 'percentiles_examine', 'regs_disp2', 'levels_CNDD_dist2', 'num_regs_disp2', 'num_levels_CNDD_dist2', 'num_seeds', 'dists_examine', 'surv_curves')

%% Plot the survival curves for several the regimes above:

load('disp_reg_survival_curves.mat')

cols = [0 .1 .9 ; 0.9 0 .1 ]; %different for every regime
syms = {':', '--', '-', '-'};
lws = [1.5 1.5 1.5 2.5];

s1 = subplot(4,2,8);
hold on
set(s1,'FontSize',16,'FontWeight','bold','LineWidth',1.5, 'YScale','log');
%xlabel('Percentile of dispersal distribution')
%ylabel('Survival probability')
text(0.91,0.89,'(f)','FontSize',16,'FontWeight','bold')
ylim([.025 1]);

for rr = 1:num_regs_disp2

    for gg = 1:num_levels_CNDD_dist2
        
        plot(percentiles_examine, surv_curves{rr, gg}, syms{gg},'color',cols(rr,:),'LineWidth', lws(gg))

    end
end

text(0.02, 0.035, '\Delta_C=7 m.', 'Color',[0 .1 .9], 'FontSize',14)
text(0.32, 0.035, '\Delta_C=60 m.', 'Color',[0.9 0 .1], 'FontSize',14)
text(0.65, 0.037, 'Dash: disp.',  'FontSize',14)

%%%%%%%%%%%%%%%%%%%%%%%%%%%%%%%%%%%%%%%%%%%%%%%%%%%%%%%%%%%%%%%%%%%%%%%%
%% Part three (second figure): CNDD with Density statistics (second order)
%%%%%%%%%%%%%%%%%%%%%%%%%%%%%%%%%%%%%%%%%%%%%%%%%%%%%%%%%%%%%%%%%%%%%%%%

%% plot EA and PCF20 (a + b):

cols = [0 .1 .9 ; 0 .9 .1 ; .9 0 .1 ; 0 .1 .9  ];
syms = {'-o', '-o', '-o', '-x'};
legend_entries_regs = {'disp.: 7 m., Q_C=5', 'disp.: 20 m., Q_C=5', 'disp.: 60 m., Q_C=5', 'disp.: 7 m., Q_C=0.2'};

xvalues = [7 20 60 180];
xtick_text = {'7', '20', '60', '\infty'};

% o: Qc = 5 , x: 0.2.
% blue: dispersal distance = 7 m, green: 20 m.., red: inf

% [dispersal distance , CNDD distance , CNDD magnitude , GNDD magnitude]
%regs_CNDD = [1 1 4 2 ; 2 1 4 2 ; 3 1 4 2 ; 1 1 2 2 ];

figure();

s1 = subplot(3,2,3);
hold on
set(s1,'FontSize',16,'FontWeight','bold','LineWidth',1.5, 'XScale','log');
for rr = 1:reg_num
    plot(xvalues ,EA20s_CNDD(rr,:),syms{rr},'color',cols(rr,:),'MarkerFaceColor',cols(rr,:),'LineWidth',1.5)
end

xticks(xvalues);
xticklabels(xtick_text)
legend(legend_entries_regs,'FontSize',12)
xlabel('CNDD distance (\Delta_C (m.))')
ylabel('Excess Abundance (EA(20))')
text(8,-0.1,'(a)','FontSize',16,'FontWeight','bold')

s1 = subplot(3,2,5);
hold on
set(s1,'FontSize',16,'FontWeight','bold','LineWidth',1.5, 'YScale','log', 'XScale','log');
for rr = 1:reg_num
    plot(xvalues ,RN20_CNDD(rr,:),syms{rr},'color',cols(rr,:),'MarkerFaceColor',cols(rr,:),'LineWidth',1.5)
end

xticks(xvalues);
xticklabels(xtick_text)
xlabel('CNDD distance (\Delta_C (m.))')
ylabel('Relative Density (RND(20))')
text(8,100,'(b)','FontSize',16,'FontWeight','bold')


%         EA20s_CNDD(rr,cc) = nansum(dat.samps_per_bin.*(dat.log_N20 - refs{regs_CNDD(rr,1)}.log_N20))/nansum(dat.samps_per_bin);
%         EDs_CNDD(rr,cc) = nansum(dat.samps_per_bin.*(dat.mean_log_dist - refs{regs_CNDD(rr,1)}.mean_log_dist))/nansum(dat.samps_per_bin);
%         CEs_CNDD(rr,cc) = nansum(dat.samps_per_bin.*(dat.CE))/nansum(dat.samps_per_bin);
%         RN20_CNDD(rr,cc)


%%%%%%%%%%%%%%%%%%%%%%%%%%%%%%%%%%%%%%%%%%%%%%%%%%%%%%%%%%%%%%%%%%%%%%%%
%% Part Four (second figure): CNDD with Density statistics (second order)
%%%%%%%%%%%%%%%%%%%%%%%%%%%%%%%%%%%%%%%%%%%%%%%%%%%%%%%%%%%%%%%%%%%%%%%%

%% plot EA and PCF20 (a + b):


cols = [0 .1 .9 ; 0 .9 .1 ; .9 0 .1 ; 0 .1 .9 ];
syms = {'-o', '-o', '-o', '-x'};
legend_entries_regs = {'\Delta_c = 7 m., Q_C = 5', '\Delta_c = 20 m., Q_C = 5', '\Delta_c = 60 m., Q_C = 5', '\Delta_c = 7 m., Q_C = 0.2'};
xvalues = [7 20 60 180];
xtick_text = {'7', '20', '60', '\infty'};

% o: Q_c = 5, x: Q_c = .2.
% blue: disp = 7, green: disp 20, disp: 60

% [dispersal distance , CNDD distance , CNDD magnitude , GNDD magnitude]
%regs_disp = [1 1 4 2 ; 1 2 4 2 ; 1 3 4 2 ; 1 1 2 2 ];

s1 = subplot(3,2,4);
hold on
set(s1,'FontSize',16,'FontWeight','bold','LineWidth',1.5, 'XScale','log');
for rr = 1:reg_num
    plot(xvalues ,EA20s_disp(rr,:),syms{rr},'color',cols(rr,:),'MarkerFaceColor',cols(rr,:),'LineWidth',1.5)
end

xticks(xvalues);
xticklabels(xtick_text)
legend(legend_entries_regs,'FontSize',12)
xlabel('Dispersal distance (m.)')
%ylabel('Excess Distance (ED)')
%xlim([0 6])
text(8,-0.1,'(c)','FontSize',16,'FontWeight','bold')
annotation('doublearrow',[.95 .95],[.3 .6],'LineWidth',2.5)
t = text(250,2,'Overdispersed', 'FontSize', 16', 'FontWeight', 'bold');
set(t,'Rotation',90)
t = text(250,0,'Aggregated', 'FontSize', 16', 'FontWeight', 'bold');
set(t,'Rotation',90)

s1 = subplot(3,2,6);
hold on
set(s1,'FontSize',16,'FontWeight','bold','LineWidth',1.5, 'XScale','log', 'YScale', 'log');
for rr = 1:reg_num
    plot(xvalues ,RND20_disp(rr,:),syms{rr},'color',cols(rr,:),'MarkerFaceColor',cols(rr,:),'LineWidth',1.5)
end

xticks(xvalues);
xticklabels(xtick_text)
xlabel('Dispersal distance (m.)')
%ylabel('Clark-Evans')
%xlim([0 6])
text(8,80,'(d)','FontSize',16,'FontWeight','bold')
```
