## Appendix S4 for "Spatial patterns from a dispersal limitation perspective: revealing biotic interactions": generate_fig_DD_magnitude4.m

```
 %%%%%%%%%%%%%%%%%%%%%%%%%%%%%%%%%%%%%%%%%%%%%%%%%%%%%%%%%%%%%%%%%%%%
% Generate figures of the effect of DD magnitude on the spatial statistics 
%%%%%%%%%%%%%%%%%%%%%%%%%%%%%%%%%%%%%%%%%%%%%%%%%%%%%%%%%%%%%%%%%%%%

% Author: Michael Kalyuhny
%
% First Written: 22/10/21
% Last modified: 29/01/22

%%%%%%%%%%%%%%%%%%%%%%%%%%%%%%%%%%%%%%%%%%%%%%%%%%%%%%%%%%%%%%%%%%%%%%%%%%
%% PART ONE: EFFECT OF GNDD with CNDD:
%%%%%%%%%%%%%%%%%%%%%%%%%%%%%%%%%%%%%%%%%%%%%%%%%%%%%%%%%%%%%%%%%%%%%%%%%%

%% Set  up parameters to use:

% each row of the matrix is a parameter regime with order:
% [dispersal distance , CNDD distance , CNDD magnitude , GNDD
% magnitude]
% here the GNDD magnitude parameter is written but all levels are used

regs_GC = cell(4,4,2); %1st dim: GNDD; 2nd dim: CNDD; 3rd dim: dispersal
for gg = 1:4
    for cc=1:4
        for dd=1:2
            regs_GC{gg,cc,dd} = [dd,2,cc,gg]; %the indexes of the regimes to consider
        end
    end
end


%% import_null:

refs = cell(1,4);
dists = {'7' '20' '60' '3000'}; %the distances used (in m.)

for dd = 1:length(dists)    
    refs{dd} = load(['imm_range_' dists{dd} '_neut_analysis_results.mat']); %null
end

%% import and analyze data for GNDD:

load('inp_all.mat')

% in the data matrices for plotting, each row is a regime and each column is a CNDD level

EA20s_GC = nan(4,4,2);
EDs_GC = nan(4,4,2);
CEs_GC = nan(4,4,2);
RN20_GC = nan(4,4,2);

for gg = 1:4
    for cc=1:4
        for dd=1:2
            dat = load(['ext1_results/' inp_all(regs_GC{gg,cc,dd}(1), regs_GC{gg,cc,dd}(2), regs_GC{gg,cc,dd}(3), regs_GC{gg,cc,dd}(4)).output_file '_analysis_results.mat']);

            EA20s_GC(gg,cc,dd) = nansum(dat.samps_per_bin.*(dat.log_N20 - refs{dd}.log_N20))/nansum(dat.samps_per_bin);
            EDs_GC(gg,cc,dd) = nansum(dat.samps_per_bin.*(dat.mean_log_dist - refs{dd}.mean_log_dist))/nansum(dat.samps_per_bin);
            CEs_GC(gg,cc,dd) = nansum(dat.samps_per_bin.*(dat.CE))/nansum(dat.samps_per_bin);
            RN20_GC(gg,cc,dd) = nansum(dat.samps_per_bin.*(dat.RND20))/nansum(dat.samps_per_bin);

        end
    end
end

%% plot ED for GNDD and CNDD:

f = figure()

s1 = subplot(2,2,1);
imagesc(EDs_GC(4:-1:1,:,1),[0 1])
colormap hot
set(s1,'FontSize',16,'FontWeight','bold','LineWidth',1.5);
ylabel('GNDD magnitude (Q_G)')
xlabel('CNDD magnitude (Q_C)')
xticklabels({'0','0.2','1','5'})
xticks([1 2 3 4])
yticklabels({'5','1','0.2','0'})
title('Excess Distance (ED)')
text(4.2,0.7,'(a)','FontSize',16,'FontWeight','bold')
text(3.6,1.1,'D = 7 m.','FontSize',16,'FontWeight','bold')

s1 = subplot(2,2,2);
imagesc(EDs_GC(4:-1:1,:,2),[0 1])
colormap hot
set(s1,'FontSize',16,'FontWeight','bold','LineWidth',1.5);
xticklabels({'0','0.2','1','5'})
xticks([1 2 3 4])
yticklabels({'5','1','0.2','0'})
text(4.2,0.7,'(b)','FontSize',16,'FontWeight','bold')
text(3.6,1.1,'D = 20 m.','FontSize',16,'FontWeight','bold')
colorbar

%%%%%%%%%%%%%%%%%%%%%%%%%%%%%%%%%%%%%%%%%%%%%%%%%%%%%%%%%%%%%%%%%%%%%%%%%%
%% PART TWO: EFFECT OF CNDD MAGNITUDE - compare with CSR
%%%%%%%%%%%%%%%%%%%%%%%%%%%%%%%%%%%%%%%%%%%%%%%%%%%%%%%%%%%%%%%%%%%%%%%%%%

%% Set  up parameters to use:

% each row of the matrix is a parameter regime with order:
% [dispersal distance , CNDD distance , CNDD magnitude , GNDD
% magnitude]
% here the CNDD magnitude parameter is written but all levels are used

regs_CNDD = [1 2 1 1 ; 1 2 1 4 ; 2 2 1 1 ; 2 2 1 4];
CNDD_levels = 4;

reg_num = size(regs_CNDD,1);
%% import_null:

refs = cell(1,4);
dists = {'7' '20' '60' '3000'}; %the distances used (in m.)

for dd = 1:length(dists)    
    refs{dd} = load(['imm_range_' dists{dd} '_neut_analysis_results.mat']); %null
end

%% import and analyze data for CNDD:

load('inp_all.mat')

% in the data matrices for plotting, each row is a regime and each column is a CNDD level

EA20s_CNDD = nan(reg_num, CNDD_levels);
EDs_CNDD = nan(reg_num, CNDD_levels);
CEs_CNDD = nan(reg_num, CNDD_levels);
RND20_CNDD = nan(reg_num, CNDD_levels);

for rr = 1:reg_num %run over the different regimes
    for cc = 1:CNDD_levels
        
        dat = load(['ext1_results/' inp_all(regs_CNDD(rr,1), regs_CNDD(rr,2), cc, regs_CNDD(rr,4)).output_file '_analysis_results.mat']);

        EA20s_CNDD(rr,cc) = nansum(dat.samps_per_bin.*(dat.log_N20 - refs{regs_CNDD(rr,1)}.log_N20))/nansum(dat.samps_per_bin);
        EDs_CNDD(rr,cc) = nansum(dat.samps_per_bin.*(dat.mean_log_dist - refs{regs_CNDD(rr,1)}.mean_log_dist))/nansum(dat.samps_per_bin);
        CEs_CNDD(rr,cc) = nansum(dat.samps_per_bin.*(dat.CE))/nansum(dat.samps_per_bin);
        RND20_CNDD(rr,cc) = nansum(dat.samps_per_bin.*(dat.RND20))/nansum(dat.samps_per_bin);

    end
end

%% plot ED for CNDD:

cols = [ 1 .5 0 ; .57 0 .58 ; 1 .5 0 ; .57 0 .58];
syms = {'-x', '-x', '-o', '-o'};
legend_entries_regs = {'D=7 m., Q_G=0', 'D=7 m., Q_G=5', 'D=20 m., Q_G=0', 'D=20 m., Q_G=5'};

% [dispersal distance , CNDD distance , CNDD magnitude , GNDD magnitude]

s1 = subplot(2,2,3);
hold on
set(s1,'FontSize',16,'FontWeight','bold','LineWidth',1.5);
for rr = 1:reg_num
    plot([0 .2 1 5] ,EDs_CNDD(rr,:),syms{rr},'color',cols(rr,:),'MarkerFaceColor',cols(rr,:),'LineWidth',1.5)
end


xlabel('CNDD magnitude (Q_C)')
ylabel('Excess Distance (ED)')
xlim([0 6])
ylim([0 1])
text(5.35,0.92,'(c)','FontSize',16,'FontWeight','bold')
text(0.5,0.92,'Null: DL','FontSize',16,'FontWeight','bold')
legend(legend_entries_regs,'FontSize',10)

% 
s1 = subplot(2,2,4);
hold on
set(s1,'FontSize',16,'FontWeight','bold','LineWidth',1.5, 'YScale','log');
for rr = 1:reg_num
    plot([0 .2 1 5] ,CEs_CNDD(rr,:),syms{rr},'color',cols(rr,:),'MarkerFaceColor',cols(rr,:),'LineWidth',1.5)
end

ylabel('Clark-Evans')
xlim([0 6])
text(5.35,0.5,'(d)','FontSize',16,'FontWeight','bold')

annotation('doublearrow',[.95 .95],[.2 .3],'LineWidth',2.5)
t = text(7,0.5,'Overdispersed', 'FontSize', 14', 'FontWeight', 'bold');
set(t,'Rotation',90)
t = text(7,.1,'Aggregated', 'FontSize', 14', 'FontWeight', 'bold');
set(t,'Rotation',90)
text(0.2,0.5,'Null: CSR','FontSize',16,'FontWeight','bold')


%%

set(f,'PaperSize',[11 11]); %set the paper size to what you want
print('-painters','-dpdf',f,'Figure3_QC_QG_magnitude')

%%%%%%%%%%%%%%%%%%%%%%%%%%%%%%%%%%%%%%%%%%%%%%%%%%%%%%%%%%%%%%%%%%%%%%%%%%
%% PART FIVE: SAME FIGURE WITH DENSITY METRICS:
%%%%%%%%%%%%%%%%%%%%%%%%%%%%%%%%%%%%%%%%%%%%%%%%%%%%%%%%%%%%%%%%%%%%%%%%%%

f2 = figure();

s1 = subplot(2,2,1);
imagesc(EA20s_GC(4:-1:1,:,1),[-1.6 0])
colormap hot
set(s1,'FontSize',16,'FontWeight','bold','LineWidth',1.5);
ylabel('GNDD magnitude (Q_G)')
xlabel('CNDD magnitude (Q_C)')
xticklabels({'0','0.2','1','5'})
xticks([1 2 3 4])
yticklabels({'5','1','0.2','0'})
title('Excess neighborhood Abundance (EA(20))')
text(4.2,0.7,'(a)','FontSize',16,'FontWeight','bold')
text(3.6,1.1,'D = 7 m.','FontSize',16,'FontWeight','bold')

s1 = subplot(2,2,2);
imagesc(EA20s_GC(4:-1:1,:,2),[-1.6 0])
colormap hot
set(s1,'FontSize',16,'FontWeight','bold','LineWidth',1.5);
xticklabels({'0','0.2','1','5'})
xticks([1 2 3 4])
yticklabels({'5','1','0.2','0'})
text(4.2,0.7,'(b)','FontSize',16,'FontWeight','bold')
text(3.6,1.1,'D = 20 m.','FontSize',16,'FontWeight','bold')
colorbar

%% plot EA for CNDD:

cols = [ 1 .5 0 ; .57 0 .58 ; 1 .5 0 ; .57 0 .58];
syms = {'-x', '-x', '-o', '-o'};
legend_entries_regs = {'D=7 m., Q_G=0', 'D=7 m., Q_G=5', 'D=20 m., Q_G=0', 'D=20 m., Q_G=5'};

% [dispersal distance , CNDD distance , CNDD magnitude , GNDD magnitude]

s1 = subplot(2,2,3);
hold on
set(s1,'FontSize',16,'FontWeight','bold','LineWidth',1.5);
for rr = 1:reg_num
    plot([0 .2 1 5] ,EA20s_CNDD(rr,:),syms{rr},'color',cols(rr,:),'MarkerFaceColor',cols(rr,:),'LineWidth',1.5)
end


xlabel('CNDD magnitude (Q_C)')
ylabel('EA(20)')
xlim([0 6])
%ylim([0 1])
text(5.35,-.1,'(c)','FontSize',16,'FontWeight','bold')
text(0.5,-.1,'Null: DL','FontSize',16,'FontWeight','bold')
legend(legend_entries_regs,'FontSize',10)

% 
s1 = subplot(2,2,4);
hold on
set(s1,'FontSize',16,'FontWeight','bold','LineWidth',1.5, 'YScale','log');
for rr = 1:reg_num
    plot([0 .2 1 5] ,RND20_CNDD(rr,:),syms{rr},'color',cols(rr,:),'MarkerFaceColor',cols(rr,:),'LineWidth',1.5)
end

ylabel('RND(20)')
xlim([0 6])
text(5.35,90,'(d)','FontSize',16,'FontWeight','bold')

annotation('doublearrow',[.95 .95],[.2 .3],'LineWidth',2.5)
t = text(7,5,'Overdispersed', 'FontSize', 14', 'FontWeight', 'bold');
set(t,'Rotation',90)
t = text(7,100,'Aggregated', 'FontSize', 14', 'FontWeight', 'bold');
set(t,'Rotation',90)
text(0.2,90,'Null: CSR','FontSize',16,'FontWeight','bold')

set(f,'PaperSize',[11 11]); %set the paper size to what you want
print('-painters','-dpdf',f,'Figure3_QC_QG_magnitude')

%%%%%%%%%%%%%%%%%%%%%%%%%%%%%%%%%%%%%%%%%%%%%%%%%%%%%%%%%%%%%%%%%%%%%%%%%%%
%% Generate SI Figure showing CNDD can reduce overdispersion:
%%%%%%%%%%%%%%%%%%%%%%%%%%%%%%%%%%%%%%%%%%%%%%%%%%%%%%%%%%%%%%%%%%%%%%%%%%%

regs_CNDD = [1 1 1 3 ; 1 3 1 3 ; 1 4 1 3];
CNDD_levels = 4;

reg_num = size(regs_CNDD,1);

%% import_null:

refs = cell(1,4);
dists = {'7' '20' '60' '3000'}; %the distances used (in m.)

for dd = 1:length(dists)    
    refs{dd} = load(['imm_range_' dists{dd} '_neut_analysis_results.mat']); %null
end

%% import and analyze data for CNDD:

load('inp_all.mat')

% in the data matrices for plotting, each row is a regime and each column is a CNDD level

EA20s_CNDD3 = nan(reg_num, CNDD_levels);
EDs_CNDD3 = nan(reg_num, CNDD_levels);
CEs_CNDD3 = nan(reg_num, CNDD_levels);
RND20s_CNDD3 = nan(reg_num, CNDD_levels);

for rr = 1:reg_num %run over the different regimes
    for cc = 1:CNDD_levels
        
        dat = load(['ext1_results/' inp_all(regs_CNDD(rr,1), regs_CNDD(rr,2), cc, regs_CNDD(rr,4)).output_file '_analysis_results.mat']);

        EA20s_CNDD3(rr,cc) = nansum(dat.samps_per_bin.*(dat.log_N20 - refs{regs_CNDD(rr,1)}.log_N20))/nansum(dat.samps_per_bin);
        EDs_CNDD3(rr,cc) = nansum(dat.samps_per_bin.*(dat.mean_log_dist - refs{regs_CNDD(rr,1)}.mean_log_dist))/nansum(dat.samps_per_bin);
        CEs_CNDD3(rr,cc) = nansum(dat.samps_per_bin.*(dat.CE))/nansum(dat.samps_per_bin);
        RND20s_CNDD3(rr,cc) = nansum(dat.samps_per_bin.*(dat.RND20))/nansum(dat.samps_per_bin);

    end
end

%% plot ED and CE (a + b):

cols = [0 .1 .9 ;  0 .9 .1 ; 0.9 0 .1 ];
syms = {'-o', '-o', '-o'};
legend_entries_regs = {'D=7 m., \Delta_C=7 m.', 'D=7 m., \Delta_C=60 m.', 'D=7 m., \Delta_C=\infty'};

% blue: CNDD = 7 m, green: 60 m., red: inf

% [dispersal distance , CNDD distance , CNDD magnitude , GNDD magnitude]
%regs_CNDD = [1 1 1 3 ; 1 3 1 3 ; 1 4 1 3 ; 3 1 1 3];
f3 = figure();

s1 = subplot(2,2,1);
hold on
set(s1,'FontSize',16,'FontWeight','bold','LineWidth',1.5);
for rr = 1:reg_num
    plot([0 .2 1 5] ,EDs_CNDD3(rr,:),syms{rr},'color',cols(rr,:),'MarkerFaceColor',cols(rr,:),'LineWidth',1.5)
end

ylabel('ED')
xlim([0 6])
text(5.3,0.7,'(a)','FontSize',16,'FontWeight','bold')

%%%%%%%%%%%%%%%%%%%%%%%%%%%%%%%%%%%%%%%%%%%%%%%%%%%%%%%%%%%%%%%%%%%%%%%%%%%%

s1 = subplot(2,2,2);
hold on
set(s1,'FontSize',16,'FontWeight','bold','LineWidth',1.5);
for rr = 1:reg_num
    plot([0 .2 1 5] ,EA20s_CNDD3(rr,:),syms{rr},'color',cols(rr,:),'MarkerFaceColor',cols(rr,:),'LineWidth',1.5)
end

ylabel('EA(20)')
xlim([0 6])
text(5.3,-.1,'(b)','FontSize',16,'FontWeight','bold')

%%%%%%%%%%%%%%%%%%%%%%%%%%%%%%%%%%%%%%%%%%%%%%%%%%%%%%%%%%%%%%%%%%%%%%%%%%%%

s1 = subplot(2,2,3);
hold on
set(s1,'FontSize',16,'FontWeight','bold','LineWidth',1.5);
for rr = 1:reg_num
    plot([0 .2 1 5] ,CEs_CNDD3(rr,:),syms{rr},'color',cols(rr,:),'MarkerFaceColor',cols(rr,:),'LineWidth',1.5)
end

xlabel('CNDD magnitude (Q_C)')
ylabel('Clark-Evans')
xlim([0 6])
text(5.3,0.21,'(c)','FontSize',16,'FontWeight','bold')

%%%%%%%%%%%%%%%%%%%%%%%%%%%%%%%%%%%%%%%%%%%%%%%%%%%%%%%%%%%%%%%%%%%%%%%%%%%%

s1 = subplot(2,2,4);
hold on
set(s1,'FontSize',16,'FontWeight','bold','LineWidth',1.5, 'YScale','log');
for rr = 1:reg_num
    plot([0 .2 1 5] ,RND20s_CNDD3(rr,:),syms{rr},'color',cols(rr,:),'MarkerFaceColor',cols(rr,:),'LineWidth',1.5)
end

%legend(legend_entries_regs,'FontSize',12)
legend(legend_entries_regs,'FontSize',10)
ylabel('RND(20)')
xlim([0 6])
text(5.3,110,'(d)','FontSize',16,'FontWeight','bold')

%%

set(f3,'PaperSize',[9 7]); %set the paper size to what you want
print(f3,'Fig_QC_delta_to_inf','-dpdf')
```
