## Appendix S4 for "Spatial patterns from a dispersal limitation perspective: revealing biotic interactions": generate_fig_EA_3.m

```
 %%%%%%%%%%%%%%%%%%%%%%%%%%%%%%%%%%%%%%%%%%%%%%%%%%%%%%%%%%%%%%%%%%%%
% Generate figures of EA(r) and RND(r)
%%%%%%%%%%%%%%%%%%%%%%%%%%%%%%%%%%%%%%%%%%%%%%%%%%%%%%%%%%%%%%%%%%%%

% Author: Michael Kalyuhny
%
% First Written: 17/07/21
% Last modified: 28/01/22

%% Set  up parameters to use:

% each row of the matrix is a parameter regime with order:
% [dispersal distance , CNDD distance , CNDD magnitude , GNDD
% magnitude]

regs_use = [2 1 4 1 ; 2 3 4 1 ; 2 3 1 4 ; 2 3 1 1];
reg_num = size(regs_use,1);

%% import_null:

refs = cell(1,4);
dists = {'7' '20' '60' '3000'}; %the distances used (in m.)

for dd = 1:length(dists)    
    refs{dd} = load(['imm_range_' dists{dd} '_neut_analysis_results.mat']); %null
end

%% import data:

EA = cell(1,reg_num);
PCF_dat = cell(1,reg_num);
load('inp_all.mat')

for rr = 1:reg_num %run over the different regimes
    dat = load(['ext1_results/' inp_all(regs_use(rr,1), regs_use(rr,2), regs_use(rr,3), regs_use(rr,4)).output_file '_analysis_results.mat']);
    
    %EA{rr} = dat.log_ND - refs{regs_use(rr,1)}.log_ND;
    EA{rr} = nansum( (dat.log_ND - refs{regs_use(rr,1)}.log_ND).*(dat.samps_per_bin') ) / sum(dat.samps_per_bin);
    PCF_dat{rr} = dat.PCF;
end

dist_axis = dat.dist_bin_centers;

%% plot data:

cols = [.63 .15 1 ; .63 .15 1 ; 0 .1 .9 ; .5 .85 0 ];
syms = {'-^', '-o', '-+','-.x'};
legend_entries = {'CNDD, \Delta_c=7 m.', 'CNDD, \Delta_c=60 m.', 'GNDD','no DD', 'range: GNDD', 'range: \Delta_c=7 m.', 'range: \Delta_c=60 m.'};
marker_sizes = [8 8 10 10];

% triangle: CNDD = 7 m, circle: 60 m., x: only GNDD, -.: neutal

% [dispersal distance , CNDD distance , CNDD magnitude , GNDD magnitude]
% regs_use = [1 1 4 1 ; 1 3 4 1 ; 3 1 4 1 ; 3 3 4 1 ; 1 1 1 4 ; 3 3 1 4];

f = figure();

s1 = subplot(2,1,1);
hold on
set(s1,'FontSize',14,'FontWeight','bold','LineWidth',1.5, 'YScale','log');
for rr = 1:reg_num
    plot(dist_axis,PCF_dat{rr},syms{rr},'color',cols(rr,:),'MarkerFaceColor',cols(rr,:),'LineWidth',2,'MarkerSize',8)
end

plot([0 4.5],[0.9 0.9], ':+', 'color', [.85 .15 .0], 'LineWidth', 2.5,'MarkerSize',10); 
plot([0 7],[0.75 0.75], ':^', 'color', [.85 .15 .0], 'LineWidth', 2.5,'MarkerSize',8); 
plot([0 60],[0.65 0.65], ':o', 'color', [.85 .15 .0], 'LineWidth', 2.5,'MarkerSize',8); 
line([0 200],[1 1], 'LineStyle', ':', 'color', 'black', 'LineWidth', 2)
ylabel('Relative Neighborhood Density')
legend(legend_entries,'NumColumns',2, 'FontSize', 14)
text(10,30,'(a)','FontSize',14,'FontWeight','bold')
%text(50,30,'Null: CSR','FontSize',16,'FontWeight','bold')
title('Null: CSR')
text(10,1.2,'Aggregated','FontSize',14,'FontWeight','bold')
text(10,0.8,'Overdispersed w.r.t. CSR','FontSize',14,'FontWeight','bold')


s1 = subplot(2,1,2);
hold on
set(s1,'FontSize',14,'FontWeight','bold','LineWidth',2.5);
for rr = 1:reg_num
    plot(dist_axis,EA{rr},syms{rr},'color',cols(rr,:),'MarkerFaceColor',cols(rr,:),'LineWidth',2,'MarkerSize',marker_sizes(rr))
end

plot([0 4.5],[0.1 0.1], ':+', 'color', [.85 .15 .0], 'LineWidth', 2.5,'MarkerSize',10); 
plot([0 7],[0.25 0.25], ':^', 'color', [.85 .15 .0], 'LineWidth', 2.5,'MarkerSize',8); 
plot([0 60],[0.45 0.45], ':o', 'color', [.85 .15 .0], 'LineWidth', 2.5,'MarkerSize',8); 
line([0 200],[0 0], 'LineStyle', ':', 'color', 'black', 'LineWidth', 2)
ylabel('Excess neighborhood Abundance')
text(2,1,'(b)','FontSize',14,'FontWeight','bold')
xlabel('Distance (m.)')
ylim([-2 1.1])
text(20,0.2,'Aggregated w.r.t. DL','FontSize',14,'FontWeight','bold')
text(20,-0.2,'Overdispersed w.r.t. DL','FontSize',14,'FontWeight','bold')
title('Null: DL')


%%

set(f,'PaperSize',[10 12]); %set the paper size to what you want
print(f,'Figure2','-dpdf')

%%%%%%%%%%%%%%%%%%%%%%%%%%%%%%%%%%%%%%%%%%%%%%%%%%%%%
%% plot a few distribution maps of regime with small clusters:
%%%%%%%%%%%%%%%%%%%%%%%%%%%%%%%%%%%%%%%%%%%%%%%%%%%%%
%% Load:
rr = 2;

%
com_samp = load(['ext1_results/' inp_all(regs_use(rr,1), regs_use(rr,2), regs_use(rr,3), regs_use(rr,4)).output_file '.mat'], 'com_samp');
%com_samp = load(['ext1_results/' inp_all(1, 2, 4, 1).output_file '.mat'], 'com_samp');
com_samp = com_samp.com_samp;

composition_ts = sum_pop(com_samp); % generate a matrix of abundances in samples (rows) by species (columns)

% THIS REQUIRES RUNNING THE SIMULAION
%%

samps_plot = [unidrnd(size(composition_ts,1),9,1) unidrnd(size(composition_ts,2),9,1)];

ff = figure()

for ii = 1:9
    

%find samples:
    points_inds = com_samp(:,3,samps_plot(ii,1)) == samps_plot(ii,2);
    points = com_samp(points_inds,1:2,samps_plot(ii,1));

    s1 = subplot(3,3,ii);
    hold on
    set(s1,'FontSize',16,'FontWeight','bold','LineWidth',1.5);

    scatter(points(:,1), points(:,2), 24, ...
        'MarkerEdgeColor',[0 0 0], 'MarkerFaceColor',[0.1 .9 0], 'MarkerFaceAlpha', .5)
    xlim([0 600])
    ylim([0 600])
    xticks([0 200 400 600])
    yticks([0 200 400 600])
    
    if ii == 4
        ylabel('Y (m.)')
    end
    
    if ii == 8
        xlabel('X (m.)')
    end
end
```
