## Appendix S4 for "Spatial patterns from a dispersal limitation perspective: revealing biotic interactions": generate_fig_sensitivity.m

```
%%%%%%%%%%%%%%%%%%%%%%%%%%%%%%%%%%%%%%%%%%%%%%%%%%%%%%%%%%%%%%%%%%%%%%%%%
%%% Compute and plot sensitivity to different parameters
%%%%%%%%%%%%%%%%%%%%%%%%%%%%%%%%%%%%%%%%%%%%%%%%%%%%%%%%%%%%%%%%%%%%%%%%%
%
% Author: Michael Kalyuzhny
%
% Date created: 14/10/21
% Date last modified: 28/01/22
%
%% Calculate statistics:

% use only subset without very long distances

% Regime indexing:
% [dispersal distance , CNDD distance , CNDD magnitude , GNDD
% magnitude]

CE_all = nan(3,3,4,4); %Clark-Evans for all regimes
ED_all = nan(3,3,4,4); %Excess distance for all regimes
RND20_all = nan(3,3,4,4); %relative neighborhood density at 20 meters for all regimes
EA20_all = nan(3,3,4,4); %Excess neighborhood abundance at 20 meters for all regimes

load('inp_all.mat'); %the list of all regimes
dists = {'7' '20' '60' '3000'}; %the distances used (in m.)

for dd=1:3 %dispersal distance
    ref = load(['imm_range_' dists{dd} '_neut_analysis_results.mat']); %null
    for cd = 1:3 %CNDD (Janzen-Connell) distance
        for cm = 1:4 %CNDD (Janzen-Connell) magnitude
            for gm = 1:4 %GNDD magnitude
                reg_now = load(['ext1_results/' inp_all(dd,cd,cm,gm).output_file '_analysis_results.mat']);
                
                %compute weighted means:
                CE_all(dd,cd,cm,gm) = nansum(reg_now.CE.*reg_now.samps_per_bin)/sum(reg_now.samps_per_bin);
                ED_all(dd,cd,cm,gm) = nansum((reg_now.mean_log_dist - ref.mean_log_dist).*reg_now.samps_per_bin)/sum(reg_now.samps_per_bin);
                EA20_all(dd,cd,cm,gm) = nansum((reg_now.log_N20 - ref.log_N20).*reg_now.samps_per_bin)/sum(reg_now.samps_per_bin);
                RND20_all(dd,cd,cm,gm) = nansum(reg_now.RND20.*reg_now.samps_per_bin)/sum(reg_now.samps_per_bin);

            end
        end
    end
end

%% Compute sensitivities:

%which values to compare?
ref_ind = 2; %low level (BUT NON-ZERO)
comp_ind = 3; %higher level

% Setup:
samp_size = 4*(3*3*4 + 3*3*4 + 3*3*4 + 3*3*3); %based on what's below
sens = nan(samp_size,1); %sensitivity
stat = strings(samp_size,1); %the statistic (CE/RND20/ED/EA(20))
param = strings(samp_size,1); %parameter (now: D)

s1 = table(stat,param,sens);
counter = 1;

% Dispersal:

for cd = 1:3 %CNDD distance
    for cm = 2:4 %CNDD magnitude
        for gm = 1:4 %GNDD magnitude
            s1.stat(counter) = 'CE';
            s1.sens(counter) = log(CE_all(comp_ind,cd,cm,gm) / CE_all(ref_ind,cd,cm,gm));
            s1.param(counter) = 'Dispersal distance';
            counter = counter + 1;
            
            s1.stat(counter) = 'ED';
            s1.sens(counter) = ED_all(comp_ind,cd,cm,gm) - ED_all(ref_ind,cd,cm,gm);
            s1.param(counter) = 'Dispersal distance';
            counter = counter + 1;
            
            s1.stat(counter) = 'RND20';
            s1.sens(counter) = log(RND20_all(comp_ind,cd,cm,gm) / RND20_all(ref_ind,cd,cm,gm));
            s1.param(counter) = 'Dispersal distance';
            counter = counter + 1;
            
            s1.stat(counter) = 'EA20';
            s1.sens(counter) = (EA20_all(comp_ind,cd,cm,gm) - EA20_all(ref_ind,cd,cm,gm));
            s1.param(counter) = 'Dispersal distance';
            counter = counter + 1;
        end
    end
end

% CNDD distance:

for dd = 1:3 
    for cm = 2:4
        for gm = 1:4
            s1.stat(counter) = 'CE';
            s1.sens(counter) = log(CE_all(dd,comp_ind,cm,gm) / CE_all(dd,ref_ind,cm,gm));
            s1.param(counter) = 'CNDD distance';
            counter = counter + 1;
            
            s1.stat(counter) = 'ED';
            s1.sens(counter) = ED_all(dd,comp_ind,cm,gm) - ED_all(dd,ref_ind,cm,gm);
            s1.param(counter) = 'CNDD distance';
            counter = counter + 1;
            
            s1.stat(counter) = 'RND20';
            s1.sens(counter) = log(RND20_all(dd,comp_ind,cm,gm) / RND20_all(dd,ref_ind,cm,gm));
            s1.param(counter) = 'CNDD distance';
            counter = counter + 1;
            
            s1.stat(counter) = 'EA20';
            s1.sens(counter) = (EA20_all(dd,comp_ind,cm,gm) - EA20_all(dd,ref_ind,cm,gm));
            s1.param(counter) = 'CNDD distance';
            counter = counter + 1;
        end
    end
end

% CNDD magnitude:

for dd = 1:3
    for cd = 1:3
        for gm = 1:4
            s1.stat(counter) = 'CE';
            s1.sens(counter) = log(CE_all(dd,cd,comp_ind,gm) / CE_all(dd,cd,ref_ind,gm));
            s1.param(counter) = 'CNDD magnitude';
            counter = counter + 1;
            
            s1.stat(counter) = 'ED';
            s1.sens(counter) = ED_all(dd,cd,comp_ind,gm) - ED_all(dd,cd,ref_ind,gm);
            s1.param(counter) = 'CNDD magnitude';
            counter = counter + 1;
            
            s1.stat(counter) = 'RND20';
            s1.sens(counter) = log(RND20_all(dd,cd,comp_ind,gm) / RND20_all(dd,cd,ref_ind,gm));
            s1.param(counter) = 'CNDD magnitude';
            counter = counter + 1;
            
            s1.stat(counter) = 'EA20';
            s1.sens(counter) = (EA20_all(dd,cd,comp_ind,gm) - EA20_all(dd,ref_ind,cm,gm));
            s1.param(counter) = 'CNDD magnitude';
            counter = counter + 1;
        end
    end
end

% GNDD magnitude:

for dd = 1:3
    for cd = 1:3
        for cm = 2:4
            s1.stat(counter) = 'CE';
            s1.sens(counter) = log(CE_all(dd,cd,cm,comp_ind) / CE_all(dd,cd,cm,ref_ind));
            s1.param(counter) = 'GNDD magnitude';
            counter = counter + 1;
            
            s1.stat(counter) = 'ED';
            s1.sens(counter) = ED_all(dd,cd,cm,comp_ind) - ED_all(dd,cd,cm,ref_ind);
            s1.param(counter) = 'GNDD magnitude';
            counter = counter + 1;
            
            s1.stat(counter) = 'RND20';
            s1.sens(counter) = log(RND20_all(dd,cd,cm,comp_ind) / RND20_all(dd,cd,cm,ref_ind));
            s1.param(counter) = 'GNDD magnitude';
            counter = counter + 1;
            
            s1.stat(counter) = 'EA20';
            s1.sens(counter) = (EA20_all(dd,cd,cm,comp_ind) - EA20_all(dd,cd,cm,ref_ind));
            s1.param(counter) = 'GNDD magnitude';
            counter = counter + 1;
        end
    end
end

% Transform to absolute sensitivity:
s1.sens = abs(s1.sens);

%% Plot sensitivity:

s1.stat = categorical(s1.stat);
s1.param = categorical(s1.param);
s1.param = reordercats(s1.param,{'Dispersal distance','CNDD distance','CNDD magnitude','GNDD magnitude'});

s1.stat = reordercats(s1.stat,{'CE','RND20','ED','EA20'});

boxchart(s1.stat,s1.sens,'GroupByColor',s1.param);
b = gca;
set(b,'FontSize',18,'FontWeight','bold','LineWidth',1.5);


ylabel('Sensitivity')
line([2.5 2.5],[0 2],'LineStyle','--','LineWidth',1.5,'Color',[.2 .2 .2])
lgd = legend(string(unique(s1.param)));
lgd.Title.String = 'Sensitivity to:';
text(0.6,1.9,'CSR:','FontSize',18,'FontWeight','bold')
text(2.6,1.9,'DL:','FontSize',18,'FontWeight','bold')

%%
g = gco;
set(g,'PaperSize',[11 11]); %set the paper size to what you want
print(g,'Figure5','-dpdf')
```
