## Appendix S4 for "Spatial patterns from a dispersal limitation perspective: revealing biotic interactions": generate_fig_var_Qc_ver4.m

```
%%%%%%%%%%%%%%%%%%%%%%%%%%%%%%%%%%%%%%%%%%%%%%%%%%%%%%%%%%%%%%%%%%%%%%%%%%%%
%%% Generate the figure analyzing variable CNDD
%%%%%%%%%%%%%%%%%%%%%%%%%%%%%%%%%%%%%%%%%%%%%%%%%%%%%%%%%%%%%%%%%%%%%%%%%%%%

% Author: Michael Kalyuzhny

% Date created: 10/08/21
% Date last modified: 07/11/21

%%%%%%%%%%%%%%%%%%%%%%%%%%%%%%%%%%%%%%%%%%%%%%%%%%%%%%%%%%%%%%%%%%%%%%%%%%%
%% Main text plot
%%%%%%%%%%%%%%%%%%%%%%%%%%%%%%%%%%%%%%%%%%%%%%%%%%%%%%%%%%%%%%%%%%%%%%%%%%%

%% Set up :
% Order of parameters: [D Delta_C Q_cmin Q_crange]
regs = [1 1 2 3 ; 1 3 2 3 ; 2 2 2 3 ; 3 1 2 3 ; 3 3 2 3];

min_obs_in_1D_bin = 500; %minimal number of observations

reg_inds = (sub2ind([3 3 2 3],regs(:,1),regs(:,2),regs(:,3),regs(:,4)))';
load('var_CNDD_all_analyses_results2.mat')

%remove cases with too few observations
for rr = reg_inds
    CE_sp_mean{rr}(CE_sp_samp_size{rr} < min_obs_in_1D_bin) = nan;
    ED_sp_mean{rr}(ED_sp_samp_size{rr} < min_obs_in_1D_bin) = nan;
    EA20_sp_mean{rr}(EA20_sp_samp_size{rr} < min_obs_in_1D_bin) = nan;
    RND20_sp_mean{rr}(RND20_sp_samp_size{rr} < min_obs_in_1D_bin) = nan;
end

%% First line: Distance metrics:
f1 = figure();

cols = [.9 .1 0 ; .9 .1 0 ; 0 .9 .1 ; 0 .1 .9 ; 0 .1 .9]; %dispersal distance
syms = {'x', 'o', '^', 'x', 'o'}; %CNDD distance: 7 = x, 20 = triangele, 60 = o
legend_entries = {'D=7 m., \Delta_C=7 m.', 'D=7 m., \Delta_C=60 m.', 'D=20 m., \Delta_C=20 m.', 'D=60 m., \Delta_C=7 m.', 'D=60 m., \Delta_C=60 m.'};

s1 = subplot(3,2,3);
hold on
set(s1,'FontSize',14,'FontWeight','bold','LineWidth',1.5);
counter = 1;
for rr = reg_inds
    plot(JC_all{rr} ,ED_sp_mean{rr},syms{counter}, 'color',cols(counter,:),'MarkerFaceColor',cols(counter,:),'MarkerSize',4);
    counter = counter + 1;
end

ylabel('ED')
text(1.05,0.56,'(a)','FontSize',12,'FontWeight','bold')
text(1.7,0.56,'Null: DL','FontSize',18,'FontWeight','bold')
text(1.7,-.1,'Distance metrics','FontSize',18,'FontWeight','bold','Rotation',90)
line([1 6],[0 0], 'LineStyle', ':', 'color', [.5 .5 .5], 'LineWidth', 2)

ylim([-0.02 0.6])
xlim([1 6])

s1 = subplot(3,2,4);
hold on
set(s1,'FontSize',14,'FontWeight','bold','LineWidth',1.5);
counter = 1;
for rr = reg_inds
    plot(JC_all{rr} ,CE_sp_mean{rr},syms{counter}, 'color',cols(counter,:),'MarkerFaceColor',cols(counter,:),'MarkerSize',4);
    counter = counter + 1;
end

ylabel('Clark-Evans')
text(1.05,0.95,'(c)','FontSize',12,'FontWeight','bold')
text(1.7,0.95,'Null: CSR','FontSize',18,'FontWeight','bold')
line([1 6],[1 1], 'LineStyle', ':', 'color', [.5 .5 .5], 'LineWidth', 2)
xlim([1 6])
ylim([0 1.03])

annotation('doublearrow',[.95 .95],[.4 .45],'LineWidth',2.5)
t = text(6.1,.7,'Overdispersed', 'FontSize', 11', 'FontWeight', 'bold');
set(t,'Rotation',90)
t = text(6.1,0,'Aggregated', 'FontSize', 11', 'FontWeight', 'bold');
set(t,'Rotation',90)

%% Second line: Density statistics:

s1 = subplot(3,2,5);
hold on
set(s1,'FontSize',14,'FontWeight','bold','LineWidth',1.5);
counter = 1;
for rr = reg_inds
    plot(JC_all{rr} ,EA20_sp_mean{rr},syms{counter}, 'color',cols(counter,:),'MarkerFaceColor',cols(counter,:),'MarkerSize',4);
    counter = counter + 1;
end

ylabel('EA(20)')
text(1.05,-.1,'(b)','FontSize',12,'FontWeight','bold')
text(1.7,-.1,'Density metrics','FontSize',18,'FontWeight','bold','Rotation',90)

xlabel("Species-specific added CNDD (Q_C’)")
line([1 6],[0 0], 'LineStyle', ':', 'color', [.5 .5 .5], 'LineWidth', 2)
ylim([-1.5 .03])
xlim([1 6])
legend(legend_entries,'FontSize',12,'NumColumns',2)

s1 = subplot(3,2,6);
hold on
set(s1,'FontSize',14,'FontWeight','bold','LineWidth',1.5,'YScale','log');
counter = 1;
for rr = reg_inds
    plot(JC_all{rr} ,RND20_sp_mean{rr},syms{counter}, 'color',cols(counter,:),'MarkerFaceColor',cols(counter,:),'MarkerSize',4);
    counter = counter + 1;
end

ylabel('RND(20)')
text(1.055,100,'(d)','FontSize',12,'FontWeight','bold')
line([1 6],[1 1], 'LineStyle', ':', 'color', [.5 .5 .5], 'LineWidth', 2)
ylim([0.8 110])
xlim([1 6])

annotation('doublearrow',[.95 .95],[.1 .15],'LineWidth',2.5)
t = text(6.1,1,'Overdispersed', 'FontSize', 11', 'FontWeight', 'bold');
set(t,'Rotation',90)
t = text(6.1,80,'Aggregated', 'FontSize', 11', 'FontWeight', 'bold');
set(t,'Rotation',90)

%% Export:

set(f1,'PaperSize',[10 16]); %set the paper size to what you want
print(f1,'Figure4','-dpdf')
```
