## Appendix S4 for "Spatial patterns from a dispersal limitation perspective: revealing biotic interactions": Null_run_analyze.m

```
%%%%%%%%%%%%%%%%%%%%%%%%%%%%%%%%%%%%%%%%%%%%%%%%%%%%%%%%%%%%%%%%%%%
%%% Run simulations analyzing how 4 parameters affect spatial patterns - the NULL REFERENCES
%%%%%%%%%%%%%%%%%%%%%%%%%%%%%%%%%%%%%%%%%%%%%%%%%%%%%%%%%%%%%%%%%%%
%
% Author: Michael Kalyuzhny
%
% Date created: 11/09/2020
% Date last modified: 28/12/2020
%
%%%%%%%%%%%%%%%%%%%%%%%%%%%%%%%%%%%%%%%%%%%%%%%%%%%%%%%%%%%%%%%%%%%
% Description:
%
% Here we only run the references for the 4 dispersal distances
% 
%% Set parameters:

% Set common parameters:
inp_neut = struct;

inp_neut.J = 5500; %number of individuals in the forest
inp_neut.L = 600; %landscape edge (meters)
inp_neut.S_reg = 300; %number of species in regional pool
inp_neut.b = 2; %'b' parameter of the 2DT kernel (sensu Nathan 2012), = p+1 (sensu Clark 1999), representing heavy tailedness
inp_neut.a = 12.73; %second parameter of the 2DT
inp_neut.D = 600; %quadrat edge
inp_neut.imm_prob = 0.0005; %number of immigrating seeds to add

% Time and sampling parameters (time in sweeps):
inp_neut.samps_tot = 5000; %how many samples to take?
inp_neut.samp_freq = 10; %how many sweeps between samples?
inp_neut.first_samp = 5000; %after how many sweeps to take first sample?
inp_neut.print_freq = 1000;
inp_neut.output_file = 'nan';

% Create coppies differing in dispersal distance:
range_vals = [7 20 60 3000]; %dispersal distance values 
inp_neut = repmat(inp_neut,1,4);
regimes_null = cell(1,4); 

for ii=1:length(range_vals)
    inp_neut(ii).a = range_vals(ii)*2*gamma(1)/(sqrt(pi)*gamma(0.5)); %set disperdal distance
    regimes_null{ii} = ['imm_range_' sprintf('%0.4g',range_vals(ii)) '_neut'];
    inp_neut(ii).output_file = regimes_null{ii};
end

% Set analysis parameters:
abd_bin_edges = [1 5:2:51 55:5:100 110:10:200 250:50:600 700:100:1000 1250 1500 5500];
dist_bin_edges = 10:10:200;

%% run simulations:

parfor ss = 1:length(inp_neut)
    inp = inp_neut(ss); %set input for this particular regime
    
    [com_samp, ~] = sim2_N(inp); %run and output composition
    %tmp = load([inp.output_file '.mat'],'com_samp')
    %com_samp = tmp.com_samp;
    
    Stats_regime6(com_samp, abd_bin_edges, dist_bin_edges, 0.25, 5, inp.L, 20, [inp.output_file '_analysis_results.mat']); %run analysis for this regime and save result separately
end
```
