## Appendix S4 for "Spatial patterns from a dispersal limitation perspective: revealing biotic interactions": Percentiles_to_dists.m

```
function [dists] = Percentiles_to_dists(percentiles, test_dists, a, b)
% tranforms the percentiles of distance to distances, assuming kernels is 2DT

% % Example input:
% percentiles = 0.01:0.01:0.99;
% test_dists = 0.1:0.1:2000; %distances to examine
% a = 40;
% b = 2;

% Compute the CDF of the distance distribution

DDD = @(x) 2*x.*(b - 1).*((1 + (x./(a)).^2).^(-b))./(a^2); %dispersal distance distribution
DDD_CDF = nan(1,length(test_dists));
for ii = 1:length(test_dists)
    DDD_CDF(ii) = integral(DDD,0,test_dists(ii));
end

%compute the quantile function by inverting the CDF:

dists = nan(1,length(percentiles));
for ii = 1:length(percentiles)
    [~,I] = min(abs(DDD_CDF-percentiles(ii)));
    dists(ii) = test_dists(I);
end

end
```
