## Appendix S4 for "Spatial patterns from a dispersal limitation perspective: revealing biotic interactions": plot_kernels2.m

```
%%%%%%%%%%%%%%%%%%%%%%%%%%%%%%%%%%%%%%%%%%%%%%%%%%%%%%%%%%%%%%%%%%%%%%%%%%%
%% Plot Competition Kernels:
%%%%%%%%%%%%%%%%%%%%%%%%%%%%%%%%%%%%%%%%%%%%%%%%%%%%%%%%%%%%%%%%%%%%%%%%%%%
%
% Created: 11/07/21
% Modified: 26/01/22
%
% Author: Michael Kalyuzhny

b1_1 = 4.5; %distance of GNDD
b2_1 = 5; %decay of GNDD
b1_2 = 20; %distance of one CNDD
b2_2 = 6; %decay of one CNDD

% Explore other parameters of CNDD: 
b1_3 = 20; %distance of one CNDD
b2_3 = 3; %decay of one CNDD

b1_4 = 20; %distance of one CNDD
b2_4 = 10; %decay of one CNDD

D = 0.01 : 0.01 : 40;
G1 = 1./(1 + ((1/b1_1)^b2_1).*(D.^b2_2));
C1 = 1./(1 + ((1/b1_2)^b2_2).*(D.^b2_2));
C2 = 1./(1 + ((1/b1_3)^b2_3).*(D.^b2_3));
C3 = 1./(1 + ((1/b1_4)^b2_4).*(D.^b2_4));

legend_show = {'\Delta_G = 4.5 m., \alpha_G = 5', '\Delta_C = 20 m., \alpha_G = 6', ...
                '\Delta_C = 20 m., \alpha_G = 3','\Delta_G = 4.5 m., \alpha_G = 10'};

f = figure();
axes1 = axes('Parent',f);


set(axes1,'FontSize',16,'FontWeight','bold','LineWidth',1.5);
hold(axes1,'on');
plot(D, G1,'-','LineWidth',2.5,'Color','blue');
plot(D, C1,'-','LineWidth',2.5,'Color',[1 .4 .15]);
plot(D, C2,':','LineWidth',2.5,'Color',[1 .4 .15]);
plot(D, C3,'--','LineWidth',2.5,'Color',[1 .4 .15]);
    xlim([0 D(end)])
    xticks(0:10:40);
    yticks([0 0.5 1])
    yticklabels({'0' '0.5Q' 'Q'})
    ylim([0 1.2]) %upper limit: 1.2 or the maximal R + 0.1, whichever is greater. Lower: 
    xlabel('Distance (meters)')
    ylabel('Competitive effect')
    legend(legend_show)
    %xticks([0 100 200])
    %text(2,ybounds(2)-.05, extra_labels(ss),'FontSize',16,'FontWeight','bold')
    %title(['\it' species_names{ss}])
```
