## Appendix S4 for "Spatial patterns from a dispersal limitation perspective: revealing biotic interactions": README.docx

### How to use the code appendix?

Author: Michael Kalyuzhny

Code last updated: 29/01/2022

General structure

This code appendix is aimed at reproducing all the analyses, figures and tables. The code is written in the Matlab™. As a first step, one needs to run the simulations. This is done by running the batch files within the two folders. Then, the master script ‘Full_analysis.m’ calls all other scripts, performs all analyses and produces all figures and results. Since the calculation requires hundreds of cores for hours-many days (depending on the parameters), summary of the results are included, so that the vast majority of figures can be reproduces.

**Structure of specific scripts within folders**

The hierarchy of the bullets presents which scripts call which. A ‘child’ is only presented for the first parent that calls it.

Root directory:

- ‘Full_analysis.m’ – runs all the analyses, except those run by the batch files on a cluster in the two folders 'ext1_results' and 'ext3_results'.
- ‘Null_run_analyze.m’ – runs the null models for the 4 dispersal distances.
  - ‘sim2_N.m’ – the function running the Dispersal Limitation null model.
  - ‘Stats_regime6.m’ – computes distances, densities and other statistics for a parameter regime. Saves a summary file for each regime
    - ‘sum_pop.m’ – generates a data structure of species composition by samples (time) from the samples in space and time
- ‘Analyze_var_CNDD2.m’ – computes the statistics for the simulations with variable CNDD
  - ‘Stats_regime_tables.m’ – computes species by samples tables of the statistics. These will be combined for a given parameter regime by the parent script
- ‘plot_kernels2.m’ – generate SI figure presenting the competition kernels.
- ‘generate_fig1_2.m’ – generates Figure 1, the concept of our work. Imports precalculated points.
- ‘generate_fig_EA_3.m’ – generates Figure 2, density based statistics as a function of r. Generates also examples of the distribution maps, but this requires running the simulations.
- ‘generate_fig_DD_magnitude4.m’ – generates Figure 3 and the accompanying SI figure showing the effect of CNDD and GNDD on dispersion.
- ‘generate_fig_var_Qc_ver4.m’ – generates Figure 4, variable CNDD
- ‘generate_fig_sensitivity.m’ – generates Figure 5, sensitivity to different parameters.
- ‘generate_fig_all_regs.m’ – generates the SI figures showing the four parameters over ALL regimes (of non-variable CNDD)
- ‘generate_fig_abd3.m’ – plots the relationships of abundance and dispersion.
- ‘generate_fig_DD_distance1.m’ – plots the figures of the dependence of overdispersion on CNDD and dispersal distances.
  - ‘Percentiles_to_dists.m’ – computes the percentile function of the 2DT distribution.
  - ‘Compute_survival_curve.m’ – computes the probability of survival of seeds at different percentiles of the dispersal distance distribution. Requires the simulation results to disperse the seeds around the simulated trees.

ext1_results:

- ‘ext1.sbat’ – the BASH script used to run the batch of simulations on the cluster.
- ‘Extensive_run_analyze.m’ – manages the parameter regime and runs simulations. The script runs a single simulation, getting the index of the specific regime to run from the environment. The UNIX OS should run this multiple times using the BASH script.
  - ‘sim2_1.m’ – the function running the actual simulation model.
  - ‘Stats_regime6.m’ – computes distances, densities and other statistics for a parameter regime. Saves a summary file for each regime
    - ‘sum_pop.m’ – generates a data structure of species composition by samples (time) from the samples in space and time
- Compose_pieces.m’ – service script to assemble simulation results if simulations were stopped/crashed.

Ext3_results: (STRUCTURE QUITE SIMILAR TO EXT1_RESULTS)

- ‘ext3.sbat’ – the BASH script used to run the batch of simulations on the cluster.
- ‘var_CNDD_analysis.m’ – manages the parameter regime and runs simulations. The script runs a single simulation, getting the index of the specific regime to run from the environment. The UNIX OS should run this multiple times using the BASH script.
  - ‘sim2_1.m’ – the function running the actual simulation model.
- Compose_pieces.m’ – service script to assemble simulation results if simulations were stopped/crashed.
