## Appendix S4 for "Spatial patterns from a dispersal limitation perspective: revealing biotic interactions": Stats_regime_tables.m

```
%%%%%%%%%%%%%%%%%%%%%%%%%%%%%%%%%%%%%%%%%%%%%%%%%%%%%%%%%%%%%%%%%%%%%%%%%%%%%%%%%
% Compute multiple statistics for the samples of a parameter regime IN
% TABLE FORMAT
%%%%%%%%%%%%%%%%%%%%%%%%%%%%%%%%%%%%%%%%%%%%%%%%%%%%%%%%%%%%%%%%%%%%%%%%%%%%%%%%%
%
% Author: Michael Kalyuzhny
% Date created: 22/02/2021
% Date last modified: 28/01/2022
%
% Input: 
%       1. 'com_samp' array, where first dimension is the invividual, second dimension has three elements: x,y,species ID, third dimension is the sample.
%       2. 'abd_bin_edges' array - the left edge of each abundance bin. first entry should be one, last should be
%       community size
%       3. 'min_abd' - the minimal abundance to consider
%       4. 'L' - the edge length of the landscape
%       5. 'filename' - if not NaN, the output will be saved to this location.
%
% Output:
%       1. 'mean_dist' - a table (time by species) of the mean of conspecific nearest neighbor distance in each abundance bin
%       2. 'CE' - a table of the Clark-Evans statistic
%       3. 'N20' - table of the number of neighbors within 20 meters
%       4. sp_abd - a table of abudnances
%       5. bins - a table of bins
%
%%%%%%%%%%%%%%%%%%%%%%%%%%%%%%%%%%%%%%%%%%%%%%%%%%%%%%%%%%%%%%%%%%%%%%%%%%%%%%%%%%%%%%%%%%%%

function [mean_dist, N20, CE, bins, sp_abd, abd_bin_centers] = Stats_regime_tables(com_samp, abd_bin_edges, min_abd, L, filename)
% This function computes the statistics for a parameter regime using tables

% Example input:
load('ext3_results/ext2var_dd_7_jd_7_jmin_1_jrange_0.2_ittr_1.mat', 'com_samp');
abd_bin_edges = [1 5:2:51 55:5:100 110:10:200 250:50:600 700:100:1000 1250 1500 5500];
min_abd = 5;
L = 600;
filename = 'Stats_regime_table_test.mat';

% Precalculations:
sp_abd = sum_pop(com_samp);
abd_bin_edges_use = abd_bin_edges - 0.01; %shifted edges of abundance bins, to be used
abd_bin_centers = abd_bin_edges(1:end-1) + (diff(abd_bin_edges)-1)/2; %centers of abundance bins
bins = nan(size(sp_abd));
mean_dist = nan(size(sp_abd));
N20 = nan(size(sp_abd));
CE = nan(size(sp_abd));

for sp = 1:size(sp_abd,2)
    sp
    
    for yy = 1:size(sp_abd,1)
        point_inds = com_samp(:,3,yy) == sp;
        points = com_samp(point_inds,1:2,yy);
        this_abd = size(points,1);
        
        if this_abd >= min_abd
            bins(yy,sp) = find(abd_bin_edges_use-this_abd>0,1)-1;
            
            %Analyze spatial distribution:
            
            xdists = abs(points(:,1)-points(:,1)'); %distance between every pair of points on a torus along x axis
            xdists = min(xdists,L-xdists); %torus!
            ydists = abs(points(:,2)-points(:,2)'); %distance between every pair of points on a torus along y axis
            ydists = min(ydists,L-ydists); %torus!
            dd = sqrt(xdists.^2+ydists.^2); %distances matrix

            %replace diagonal with Inf (so that your distance to yourseld does not get counted in any distance bin):
            dd(1:(this_abd+1):(this_abd^2)) = Inf;

            %find minimum in every row:
            min_dists = min(dd,[],2);

            % Now calculate statistics for this species:
            
            mean_dist(yy,sp) = mean(min_dists);
            N20(yy,sp) = mean(sum(dd<20));
            CE(yy,sp) = mean_dist(yy,sp)*2.*sqrt(this_abd/(L^2));

        end
    end
end

if ~isnan(filename) %save if required to do so
    save(filename, 'mean_dist', 'N20', 'CE', 'N20', 'bins', 'sp_abd', 'abd_bin_centers')
end

end
```
