## Appendix S4 for "Spatial patterns from a dispersal limitation perspective: revealing biotic interactions": sum_pop.m

```
function [pop_t] = sum_pop(spatial_com)
% gets as an input a community matrix with dimensions: [individuals  [x y
% sp_ind] time] and converts to [time sp_ind].
% NOTE: This code assumes that every species was sampled at least once, an assumption that makes sense when there are
% many samples and not too low immigration.

%Total numbers of samples and species:
sp_tot = max(max(spatial_com(:,3,:)));
samps_tot = length(spatial_com(1,1,:));

pop_t = zeros(samps_tot, sp_tot);
for tt = 1:samps_tot
    pop_t(tt,:) = histcounts(spatial_com(:,3,tt),0.5:(sp_tot+1));
end
end
```
