## Appendix for "Spatial patterns from a dispersal limitation perspective: revealing biotic interactions"

### Supporting information for
### Spatial patterns from a dispersal limitation perspective: revealing biotic interactions

Michael Kalyuzhny[1*], Jeffrey K. Lake[2], Annette M. Ostling[1]

1. Department of Integrative Biology, The University of Texas at Austin, 2415 Speedway, Austin, TX 78712, USA
2. Department of Biology, Adrian College, Adrian, MI, USA

#### Supplementary figures

##### Figure S1

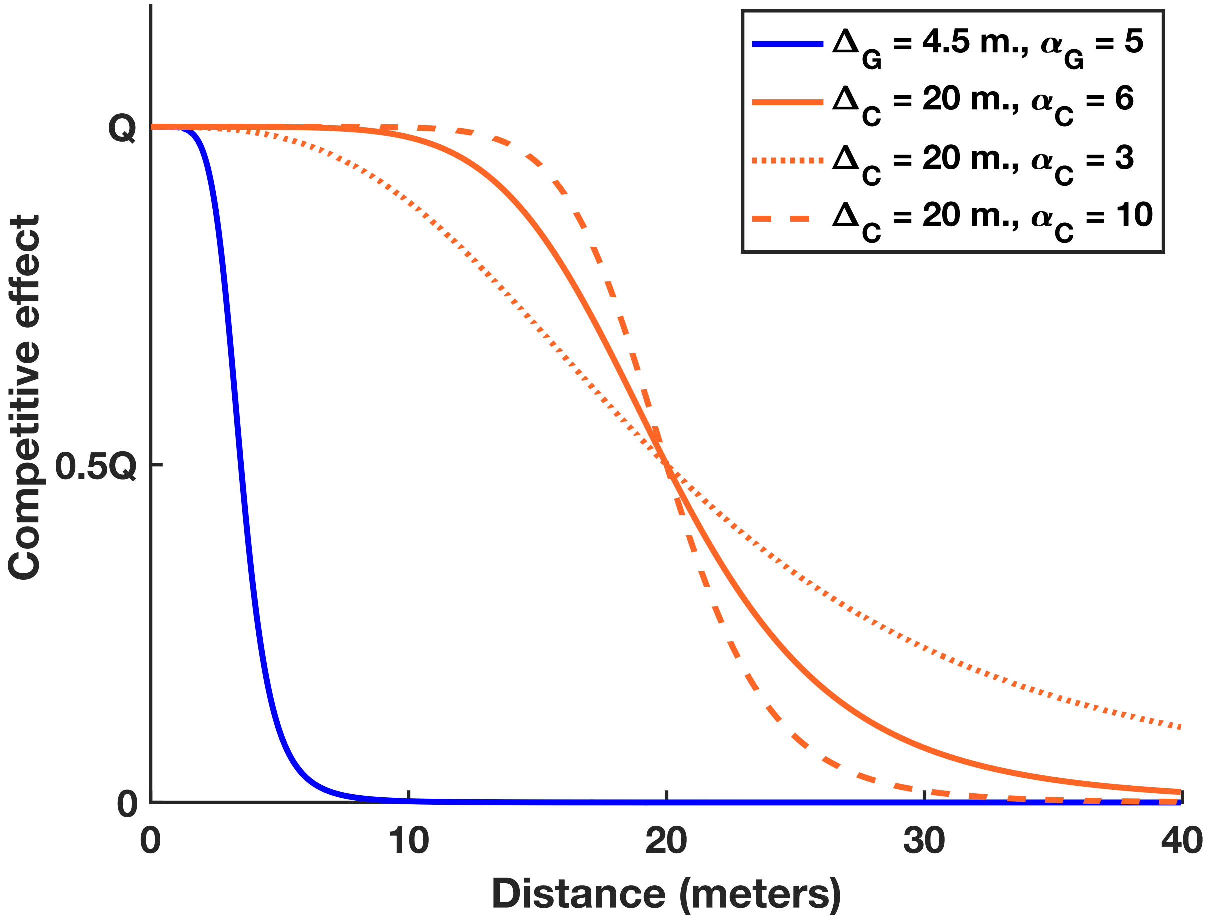


**Figure S1** Distance dependence of Negative Density Dependence (eq. 1b, 1c). The blue curve visualized eq. 1b with the parameters used for General Negative Density Dependence (GNDD): $\Delta_G = 4.5$ meters, $\alpha_G = 5$. The solid orange curve visualized eq. 1c with the parameters of the intermediate distance added CNDD: $\Delta_C = 20$ meters, $\alpha_C = 6$. The dashed orange curves visualize the effect of changing the steepness parameter, $\alpha_C$.

Figure S2

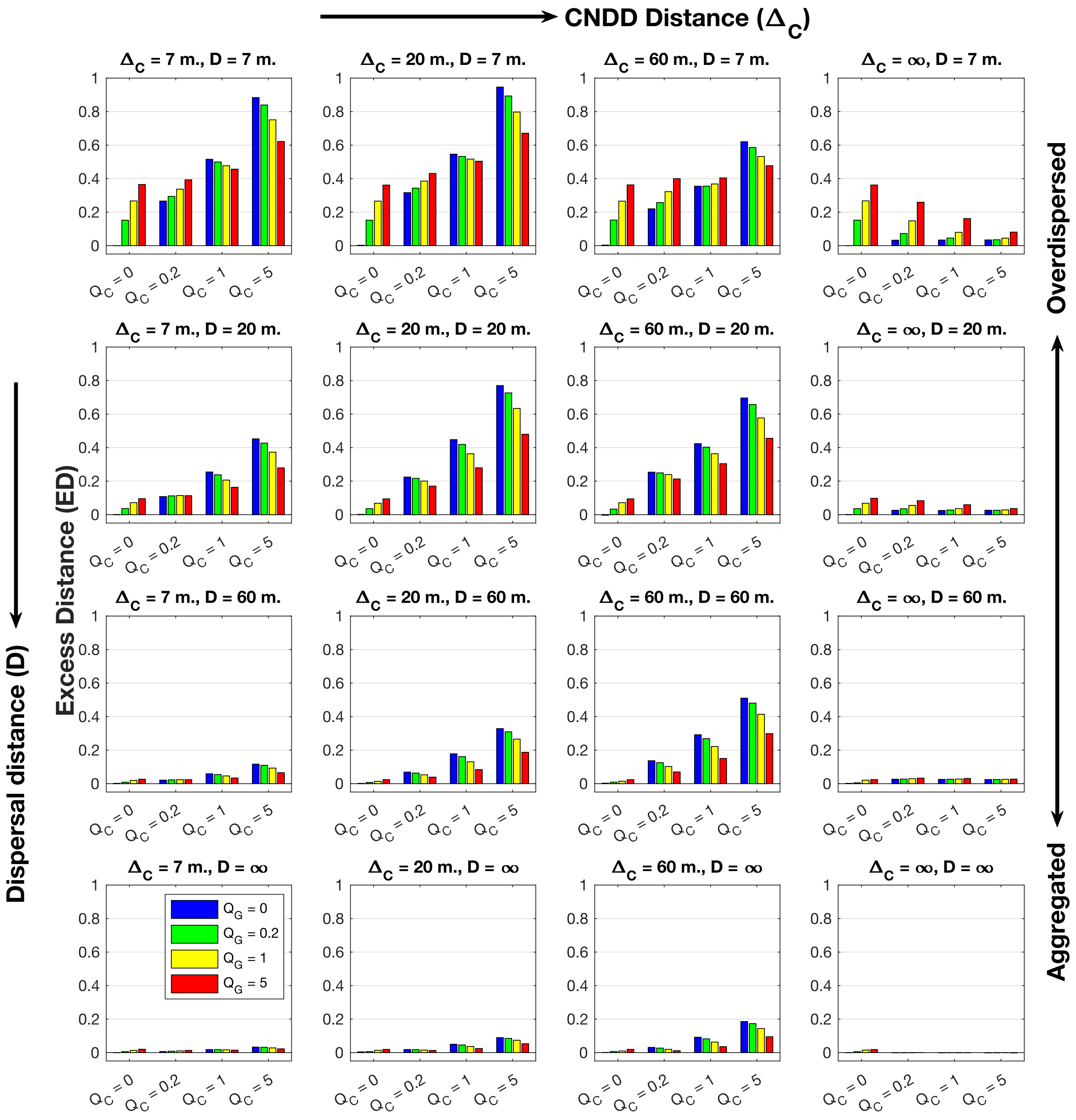


**Figure S2** The effects of dispersal distance ($D$, changes along rows), added CNDD distance ($\Delta_C$, changes along columns), the magnitude of added CNDD ($Q_C$, X axis) and GNDD ($Q_G$, color) on Excess nearest conspecific neighbor Distance (ED), which quantifies overdispersion or aggregation relative to the Dispersal Limitation (DL) null with values $> 0$ or $< 0$ respectively. Each shown value is the averaged statistic over all populations recorded in a parameter regime with abundance ≥ 5. CNDD and GNDD generate only overdispersion relative to DL.

Figure S3

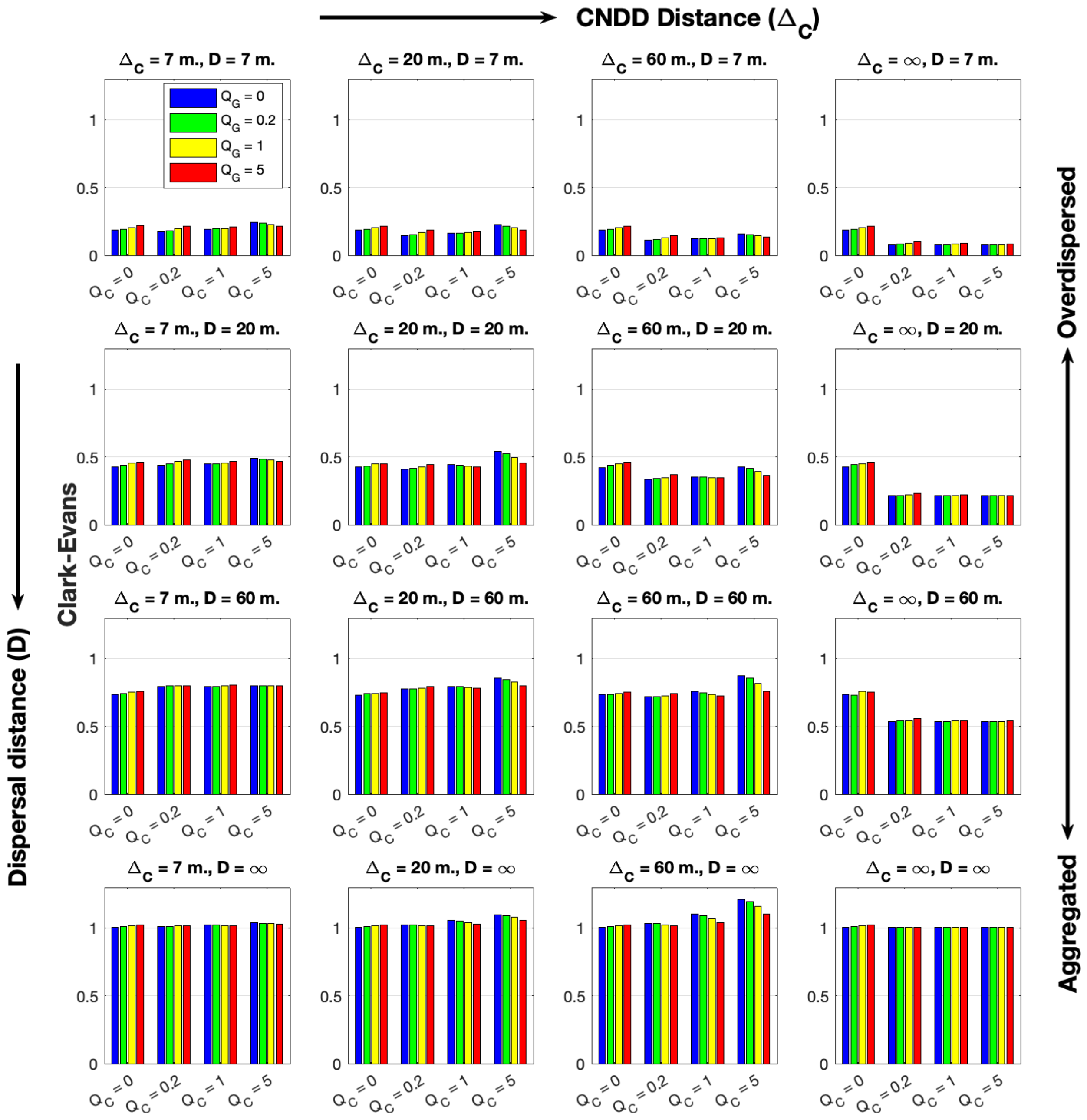


**Figure S3** The combined effect of dispersal distance (*D*, changes along rows), added CNDD distance ($\Delta_C$, changes along columns), the magnitude of added CNDD ($Q_C$, X axis) and GNDD ($Q_G$, color) on the Clark-Evans nearest neighbor statistic, which quantifies overdispersion or aggregation relative to the Complete Spatial Randomness (CSR) null with values $> 1$ or $< 1$ respectively. Each shown value is the averaged statistic over all populations recorded in a parameter regime with abundance ≥ 5. CNDD and GNDD makes populations look overdipsersed relative to CSR ONLY when they are not dispersal limited (i.e. when *D* is ∞).

Figure S4

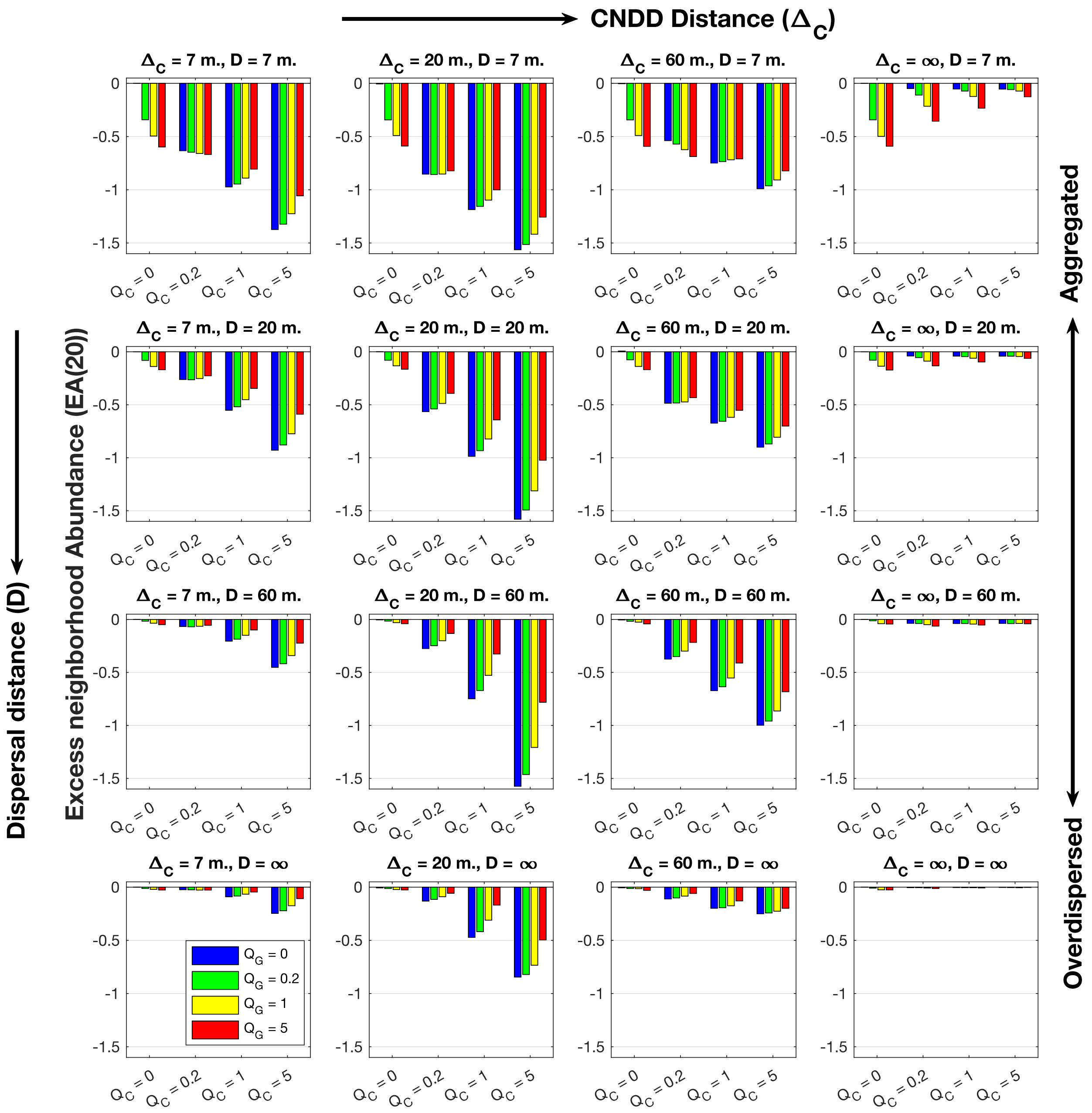


**Figure S4** The combined effect of dispersal distance (*D*, changes along rows), added CNDD distance ($\Delta_C$, changes along columns), the magnitude of added CNDD ($Q_C$, X axis) and GNDD ($Q_G$, color) on Excess neighborhood Abundance at a 20 meter radius ($EA(20)$), which quantifies overdispersion or aggregation relative to the Dispersal Limitation (DL) null with values $< 0$ or $>$ 0 respectively. Each shown value is the averaged statistic over all populations recorded in a parameter regime with abundance $\geq 5$. CNDD and GNDD generate only overdispersion relative to DL.

#### Figure S5

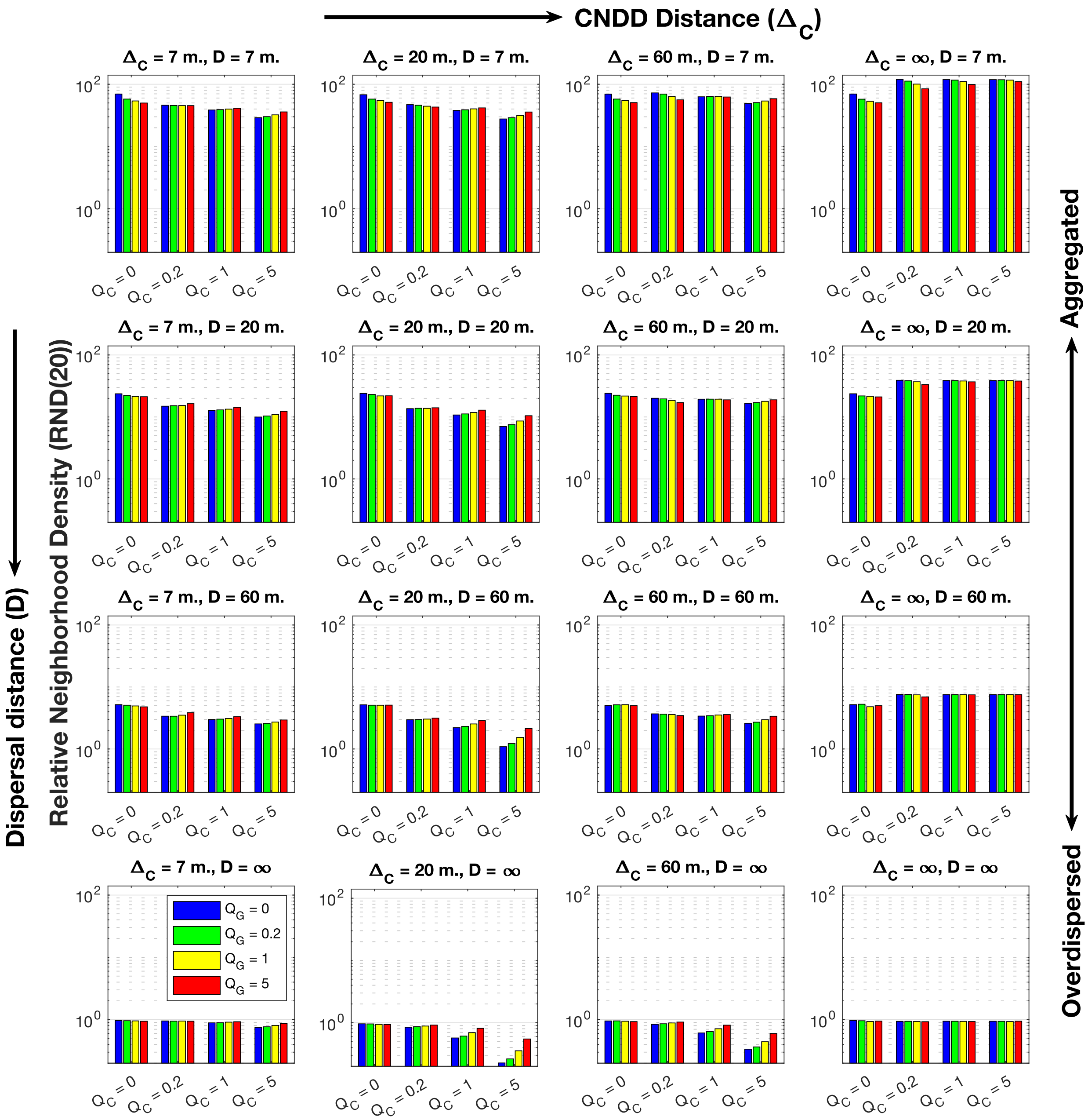


**Figure S5** The combined effect of dispersal distance (*D*, changes along rows), added CNDD distance ($\Delta_C$, changes along columns), the magnitude of added CNDD ($Q_C$, X axis) and GNDD ($Q_G$, color) on Relative Neighborhood Abundance at a 20 meter radius (*RND20*(20)), which quantifies overdispersion or aggregation relative to the Complete Spatial Randomness (CSR) null with values $< 1$ or $> 1$ respectively. Each shown value is the averaged statistic over all populations recorded in a parameter regime with abundance $\geq 5$. Note the Y axis has a logarithmic scale. CNDD and GNDD makes populations look overdispersed relative to CSR ONLY when they are not dispersal limited (i.e., when *D* is $\infty$).

Figure S6

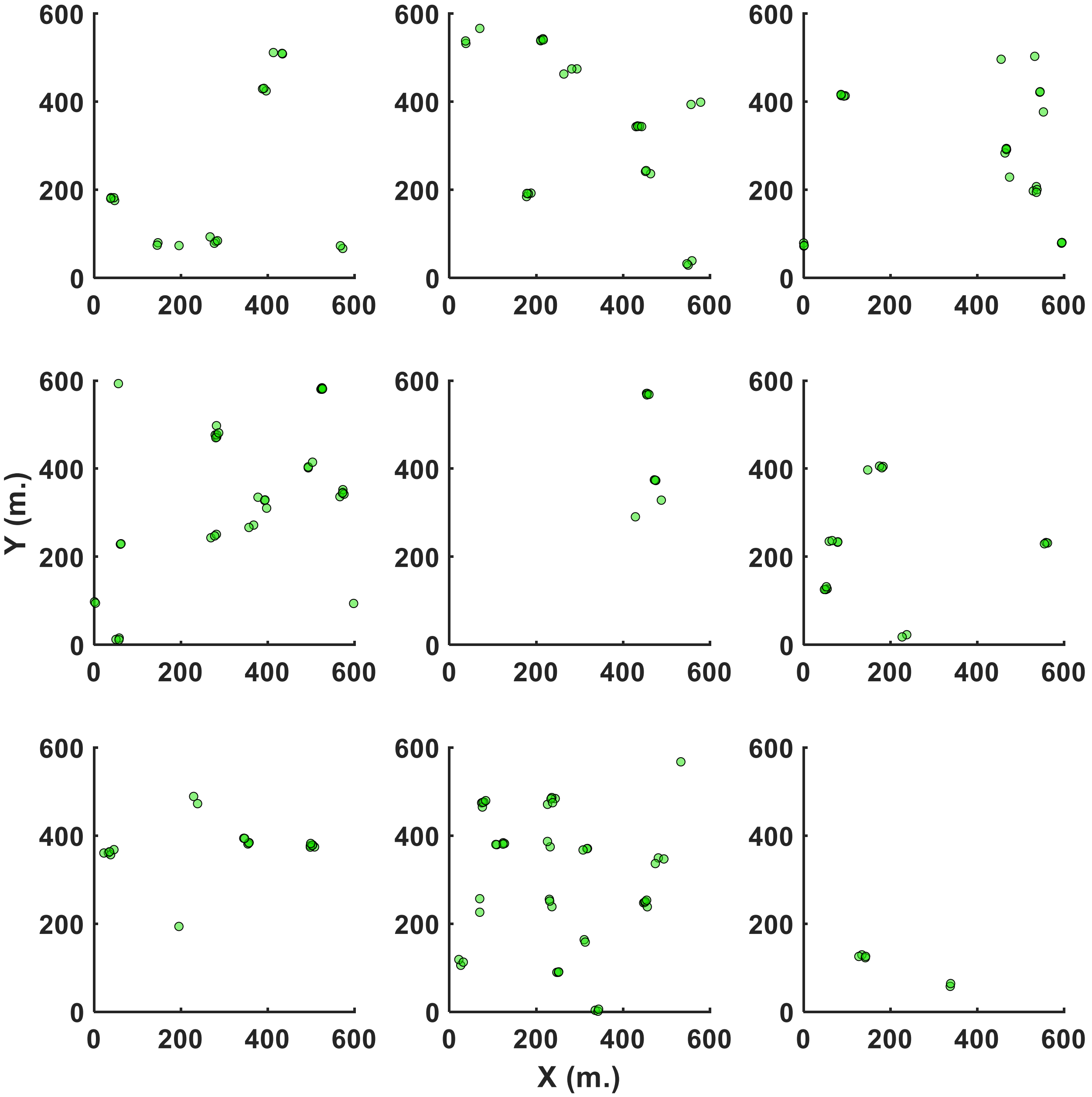


**Figure S6** Several distribution maps of simulated species when $Q_G = 0$, $Q_C = 5$, $\Delta_C = 60$ meters and mean dispersal distance (*D*) is 7 meters. These maps demonstrate the emergence of small clusters of a few individuals, separated by large distances, responsible for the 'trough' in *EA*(*r*) and *RND*(*r*) in Figure 2 of the main text. Although in Figure 2 *D* = 20 meters, a similar but less striking pattern emerges for that case.

#### Figure S7

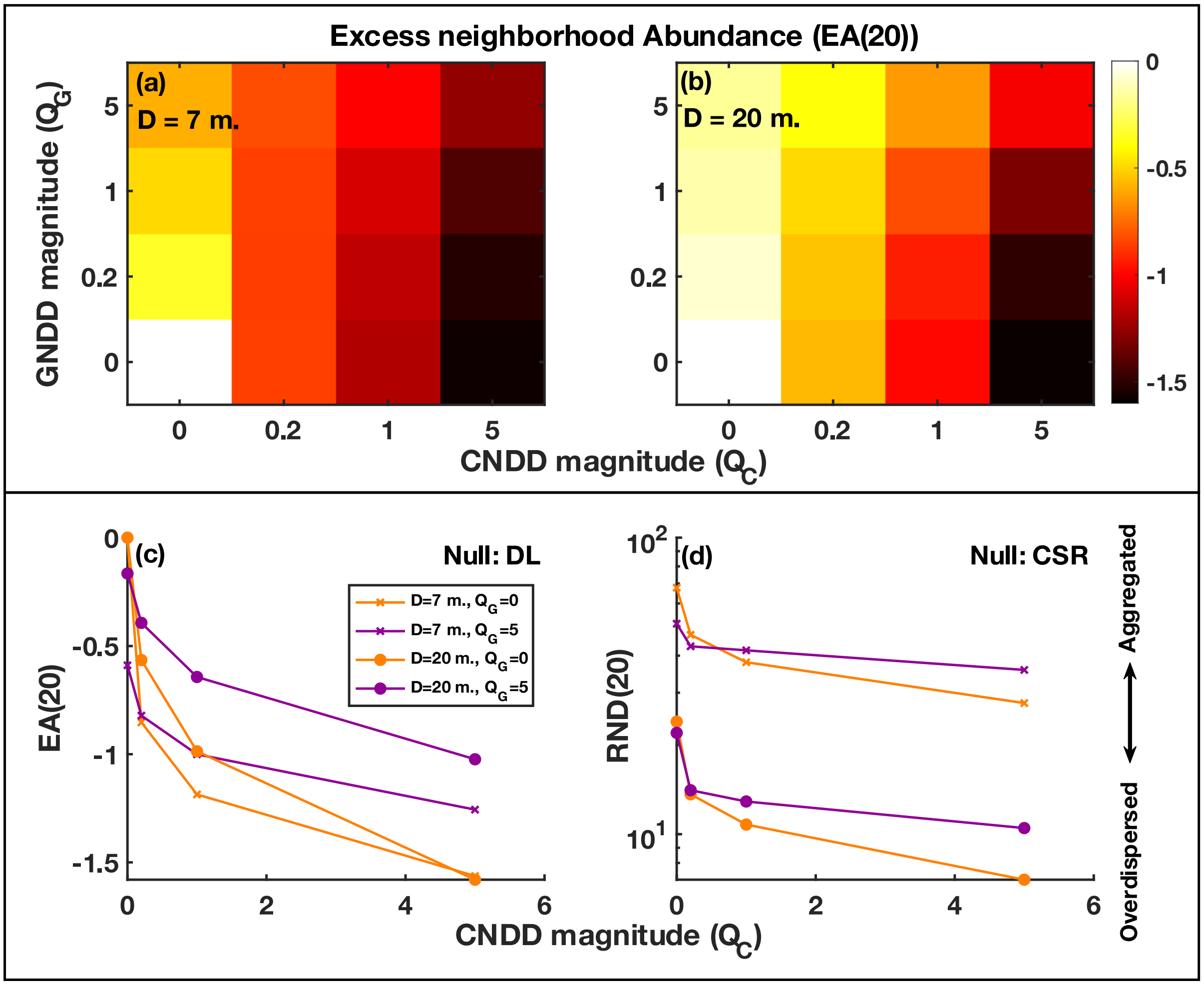


**Figure S7** Effect of the magnitude of GNDD ($Q_G$) and added CNDD ($Q_C$) on neighborhood density statistics. In all cases $\Delta_C$ = 20 meters. The colors of (**a**) and (**b**) represents the Excess 20-meter neighborhood Abundance (*EA*(20), eq. 5 of the main text) in response to a combination of GNDD and CNDD for dispersal distances (*D*) of 7 meters and 20 meters, respectively. Furthermore, the effect of $Q_C$ on (**c**) *EA*(20) and on (**d**) the Relative Neighborhood Density statistic (eq. 4 of the main text), with the latter using CSR as null, are presented when $Q_G$ = 0 (orange) or $Q_G$ = 5 (purple). GNDD alone can produce much lower overdispersion than added CNDD. GNDD counteracts the effect of added GNDD (purple versus orange curves in (**c**) and (**d**)) and the strongest overdispersion is observed when CNDD is maximal and GNDD is minimal.

Figure S8

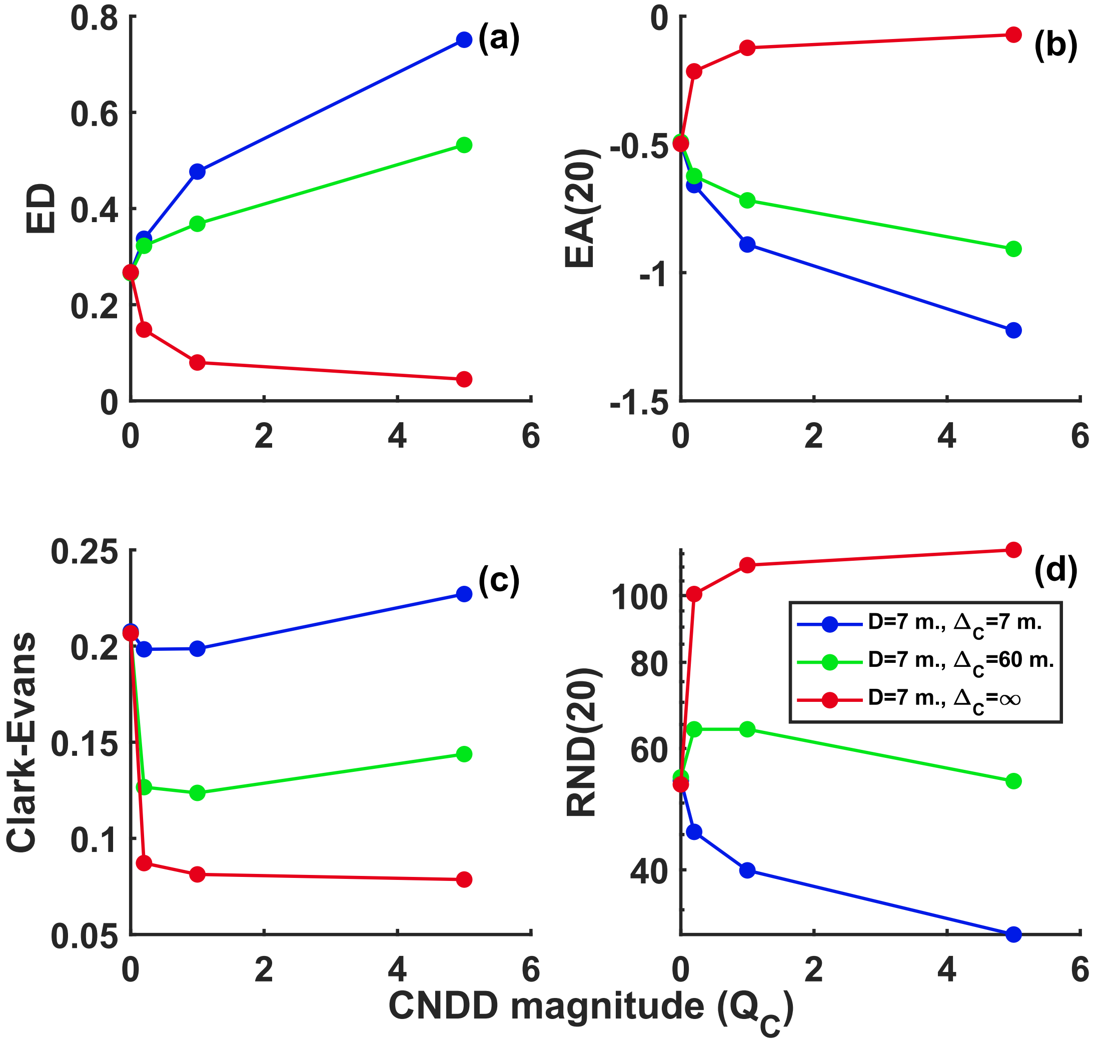


**Figure S8** Effect of the magnitude of added CNDD ($Q_C$) on spatial statistics when varying the distance of CNDD, $\Delta_C$. Blue, green and red curves correspond to $\Delta_C$ of 7 meters, 60 meters and ∞, respectively. In all cases, dispersal distance ($D$) is 7 meters and the magnitude of GNDD is $Q_G = 1$. (**a**) presents the Excess nearest neighbor Distance statistic (ED) while (**b**) presents Excess neighborhood Abundance within 20 meters ($EA(20)$), both having Dispersal Limitation (DL) as reference. (**c**) considers the Clark-Evans nearest-neighbor statistic while (**d**) examines the Relative Neighborhood Density within 20 meters ($RND(20)$) statistic, both compared to CSR. Note that when $\Delta_C$ is high, an increase in CNDD may *reduce* overdispersion.

#### Figure S9

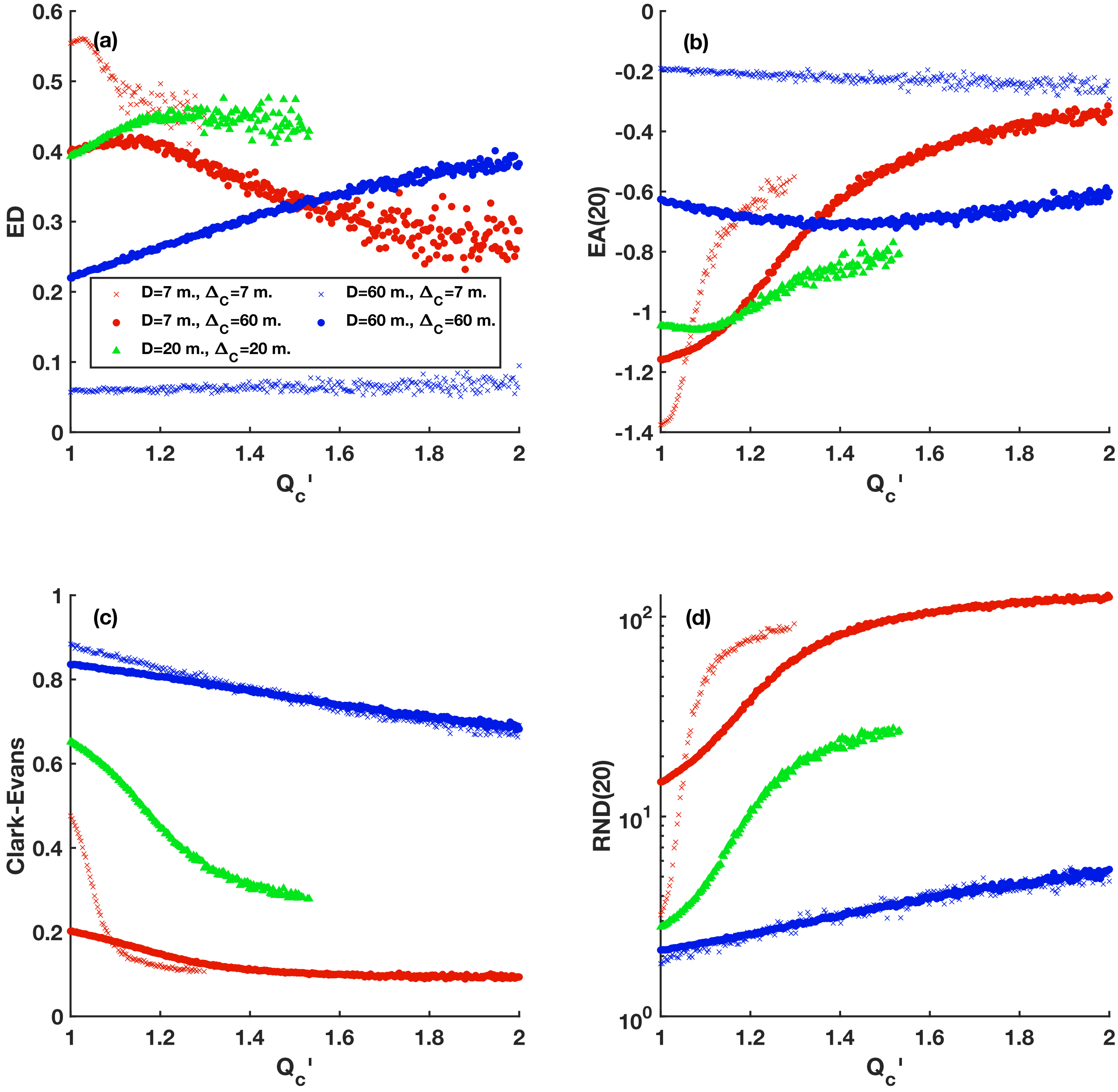


**Figure S9** Effect of the species-specific magnitude of CNDD ($Q_C$') on spatial statistics: (**a**) Excess nearest neighbor Distance (*ED*); (**b**) Excess neighborhood Abundance within 20 meters (*EA*(20), both having Dispersal Limitation (DL) as reference); (**c**) the Clark-Evans nearest neighbor statistic and (**d**) Relative Neighborhood Density within 20 meters (*RND*(20), both having CSR as reference). Different colors and symbols represent regimes with different levels of dispersal distance (*D*) and CNDD distance ($\Delta_C$), respectively. In all cases, $Q_G = 0.2$, $Q_{C\min} = 1$ and $Q_{C\text{range}} = 1$. Species with fewer then 500 observations were discarded, hence, some parameter regimes do not have observations for high $Q_C$'.

#### Appendix S1 – Glossary of important terms

| Variable / term | Meaning |
|---|---|
| **General** | |
| CSR | Complete Spatial Randomness, the standard null model for dispersion patterns, whereas the individuals appear independently and randomly across the landscape (Poisson process) |
| DL | Dispersal Limitation, the proposed null model for dispersion patterns. A spatially explicit neutral demographic model where species undergo ecological drift and disperse locally |
| CNDD | Conspecific Negative Density Dependence – reduction in the performance (in our model: recruitment) of individuals with increasing density of conspecifics |
| HNDD | Heterospecific Negative Density Dependence – reduction in the performance of individuals with increasing density of heterospecifics. |
| GNDD | General Negative Density Dependence – reduction in the performance of individuals with increasing density of all trees. HNDD represents GNDD only. |
| added CNDD | The extra negative effect that is exerted by conspecifics, on top of GNDD. CNDD = GNDD + added CNDD. |
| **Simulation models** | |
| $J$ | Total community size (value used: 5,500) |
| $m$ | Probability that a recruit will be an immigrant from the species pool (value used: $5\cdot10^{-4}$). |
| $S$ | Number of species in the pool |
| $f(r)$ | Distance distribution for seed dispersal |
| $NCI$ | Neighborhood Competition Index, measure of competition in a local neighborhood (eq. 1) |
| $N$ | Set of individuals who are the neighbors of a dispersed seed |
| $G(D_i)$ | General Negative Density Dependence (GNDD) – the competition exerted by an individual $i$ (regardless of species identity) at distance $D_i$ on seeds. |
| $C(D_i)$ | Added CNDD effect – the competition exerted by conspecific i at distance $D_i$ on seeds, on top of $G(D_i)$ (eq. 1c) |
| $Q_G, Q_C$ | The magnitudes of GNDD and added CNDD, respectively (eq. 1) |
| $\Delta_G, \Delta_C$ | The distances at which GNDD and added CNDD, respectively, fall to half of their magnitude (eq. 1) |
| $\alpha_G, \alpha_C$ | The rate at which GNDD and added CNDD, respectively, decay with distance (eq. 1) |
| $Q'_C$ | The magnitude of species-specific added CNDD in the scenario where different species have different CNDD |
| $Q_{Cmin}$ | The $Q'_C$ value of the most CNDD-resilient species (with the lowest $Q'_C$) |
| $Q_{Crange}$ | The difference between the maximal and minimal $Q'_C$ across species |
| **Spatial statistics** | |
| $\lambda$ | The density of a species in the observation window (survey plot) |
| $NND_{obs}$ | Observed distance of individuals to their nearest conspecific neighbor, averaged over individuals |
| $CE$ | The Clark-Evans statistic, comparing $NND_{obs}$ to its expectation under CSR (eq. 2). CE > 1 indicates overdispersion compared to CSR, CE < 1 indicates aggregation |
| $NND_{null}$ | Distance of individuals to their nearest conspecific neighbor, averaged over individuals, in a realization of the DL null model |
| $ED$ | Excess nearest conspecific neighbor Distance statistic, comparing $NND_{obs}$ to its expectation under DL (eq. 3). ED > 1 indicates overdispersion compared to DL. |
| $N_{obs}(r)$ | Observed number of conspecific neighbors that an average individual has between distances r and r + Δr |
| $A(r)$ | area of the annulus with inner and outer radii r and r + Δr, respectively |

| | |
|---|---|
| $N_{null}(r)$ | Number of conspecific neighbors that an average individual has between distances r and r + Δr in a realization of the DL null model |
| $RND(r)$ | Relative Neighborhood abundance, comparing $N_{obs}(r)$ to its expectation under CSR (eq. 4). RND(r) > 1 indicates aggregation compared to CSR at distance r. |
| $EA(r)$ | Excess neighborhood Abundance, comparing $N_{obs}(r)$ to its expectation under DL (eq. 5). EA(r) < 0 indicates overdispersion compared to DL at distance r. |

#### Appendix S2: Effects of dispersal and CNDD distances

The effect of added CNDD distance ($\Delta_C$) is unimodal, so that species with intermediate $\Delta_C$ are the most overdispersed. Overdispersion, as measured by *ED*, is maximized at short $\Delta_C$ if dispersal is short and at longer distances if dispersal is long (Figures S10a). From the perspective of *EA*(20), however, dispersal has a much weaker effect on the location of the peak (Figures S11a). The unimodal effect of $\Delta_C$ on overdispersion is also seen using CSR as a null, but is considerably less pronounced because the behavior of these statistics is determined almost completely by dispersal distance (Figures S10b, S11b).

CSR and DL strongly disagree over the effect of increasing dispersal distances (*D*). From the perspective of DL, increasing *D* mostly reduces overdispersion, but a unimodal effect can be seen for *ED* when $\Delta_C$ is long and the magnitude of CNDD ($Q_C$) is high (Figure S10d, S11c). On the other hand, when compared to CSR, increasing *D* quickly eliminates the aggregation of spatial patterns (Figure S10e, S11d).

These patterns are easily understood by examining the survival curves (Figure S10c,f) of seeds. These were calculated by dispersing $5 \cdot 10^5$ seeds for each percentile of the dispersal distance distribution (*f*(*r*)) from random trees observed in the simulations snapshots and recording the percentage that survived. Note that unlike previous works (e.g., Janzen 1970), our analysis considers not only the effect of the focal tree but rather the effect of all individuals in the landscape. When dispersal distance is much longer than $\Delta_C$, survival is only impacted at short distances and is otherwise uniform across space (Figure S10c. dotted red curve). As $\Delta_C$ increases (the lines in Figure S10c become more solid), survival at longer distances is reduced by CNDD (Fig S10c. dashed and solid red curves), so seeds establish farther away. Finally, as $\Delta_C$ becomes very long compared to dispersal distance, added CNDD affects all seeds similarly (and strongly), resulting in little overdispersion (Figure S10c, thick red solid curve).

Increasing dispersal distance while keeping $\Delta_C$ fixed has a similar effect (Figure S10f): if dispersal acts on much shorter distances than CNDD (Figure S10f dotted red curve) all seeds will encounter a uniform environment, generating little spatial patterns. As dispersal distances increase (the lines becoming more solid in Figure S10f), more seeds escape CNDD, reducing overdispersion. Finally, if dispersal acts on much longer distances, some seeds will randomly encounter adults and suffer from CNDD, but this will not depend on distance from the parent, making the spatial signal weaker (Figure S11f thick solid blue curve). Hence, large differences between the distances at which CNDD and dispersal operate lead to low spatial variability in mortality and a limited effect of CNDD on spatial patterns. This creates a unimodal effect of interactions distance and dispersal distance on dispersion, although for dispersal distance, a very substantial difference between the scales must occur for the unimodal effect to be observed. The unimodal effect of interaction distance is also observed when dispersion is compared to CSR. The latter unimodal effect has been observed before By Law *et al*. (2003) but only under very restrictive condition, probably due to their interaction kernel being normalized (hence, longer interactions were also weaker).

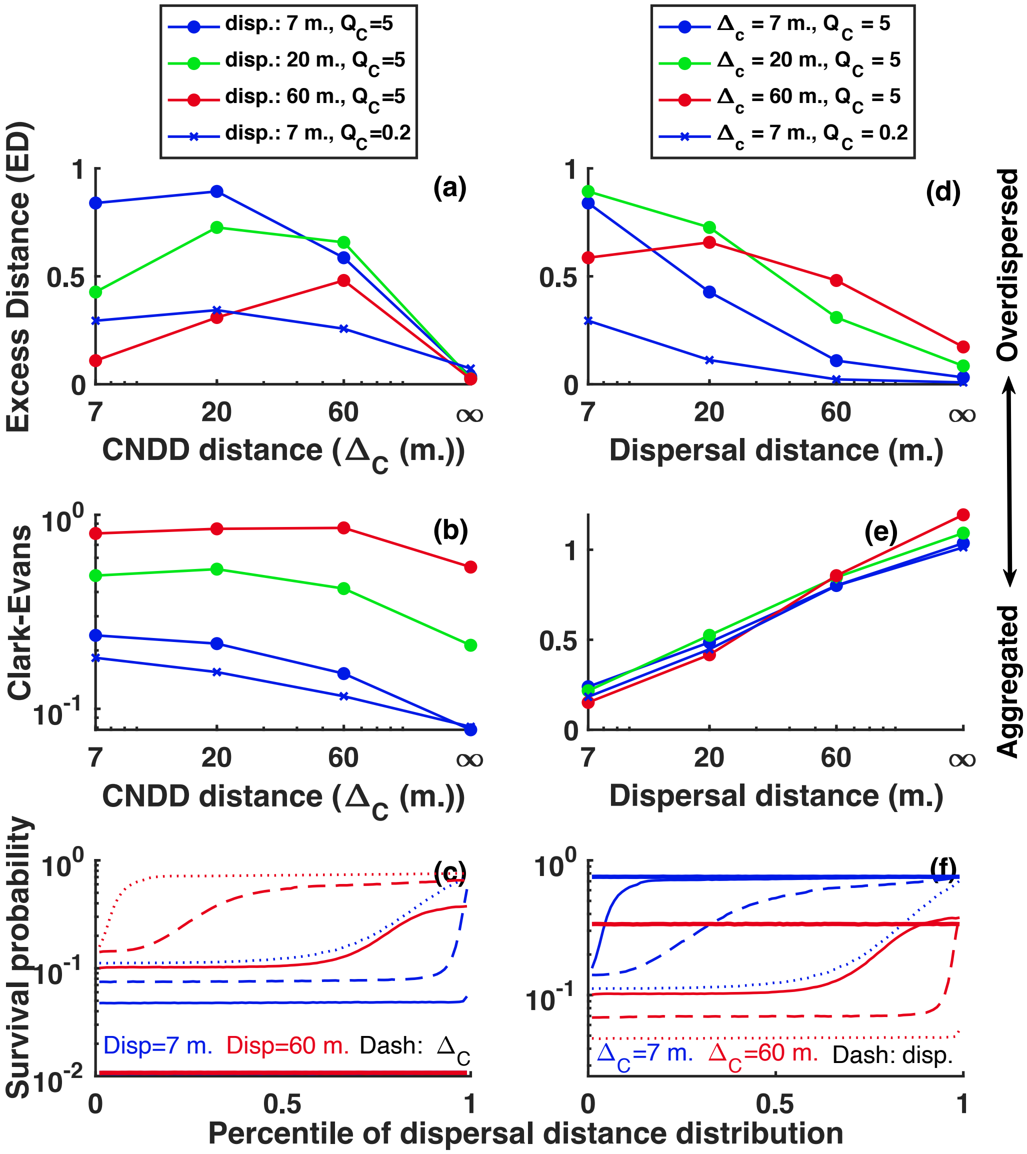


**Figure S10** Effect of dispersal and CNDD distance ($\Delta_C$) on nearest neighbor statistics. **Left side:** The effect of $\Delta_C$ on *ED* (**a**) and Clark-Evans (**b**) statistics is shown for different mean dispersal distances and CNDD magnitude ($Q_C$) when $Q_G = 0.2$. (**c**) presents the survival probability in two of the regimes from above (dispersal = 7 meters (blue) or 60 meters (red), $Q_C$ = 5) at different distances from a focal tree as $\Delta_C$ increases (dots: $\Delta_C$ = 7 meters, dash: $\Delta_C$ = 20 meters, solid line: $\Delta_C$ = 60 meters, thick solid line: $\Delta_C = \infty$). Note that the blue and red thick solid lines overlay. **Right side:** The effect of dispersal distance on *ED* (**d**) and the Clark-Evans (**e**) statistic is shown for different value of $\Delta_C$ and $Q_C$ when $Q_G = 0.2$. (**f**) presents the survival probability in two of the regimes above ($\Delta_C$ = 7 (blue) or 60 meters, $Q_C = 5$) at different distances from a focal tree as dispersal distance increases (dots: 7 meters, dash: 20 meters, solid line: 60 meters, thick solid line: $\infty$). The legend for panels (**a**) and (**b**) is above (**a**) and for panels (**d**) and (**e**) is above (**d**).

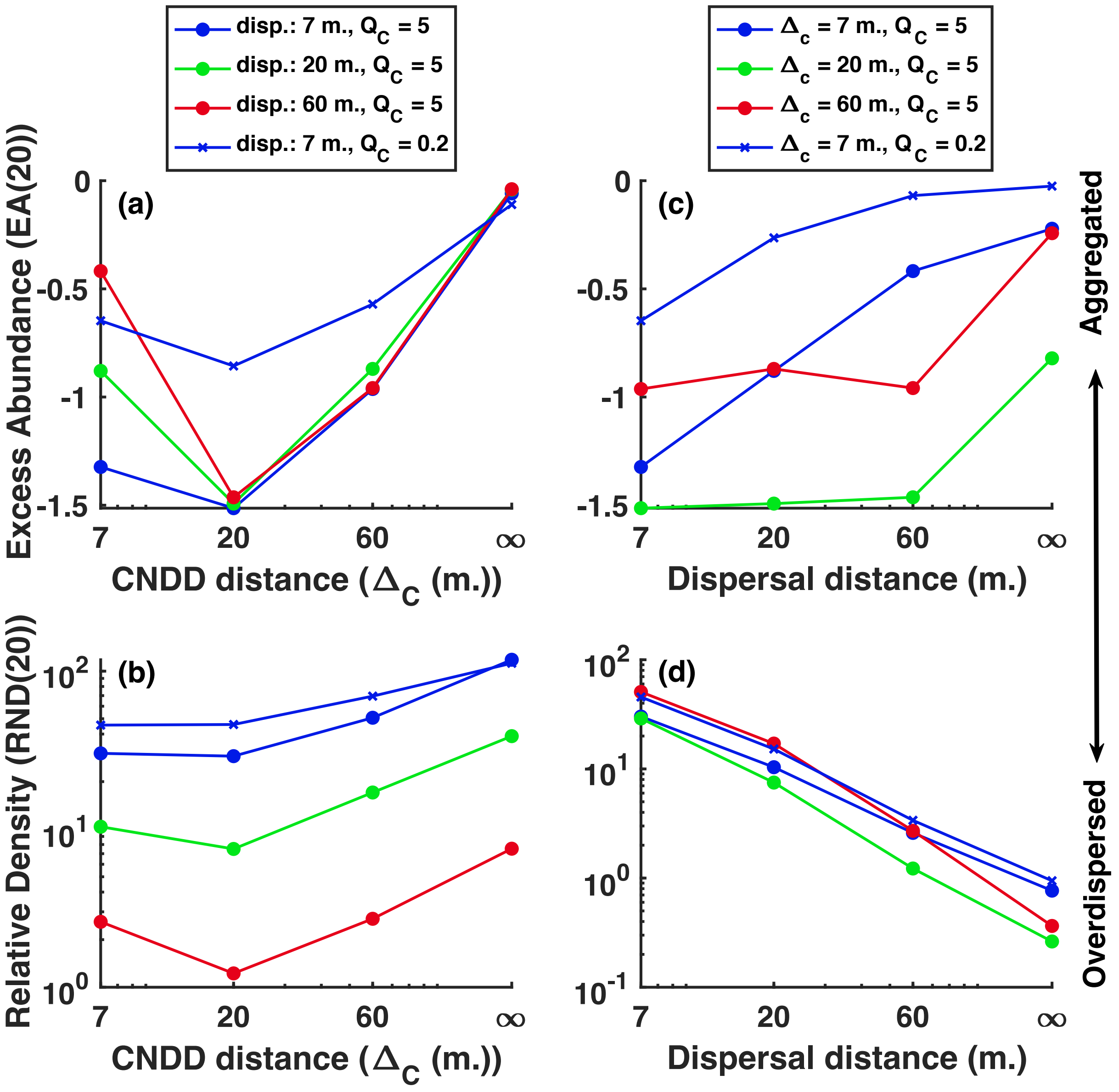


**Figure S11** Effect of Effect of dispersal and CNDD distance ($\Delta_C$) on density (second order) statistics. **Left side:** The effect of $\Delta_C$ on the $EA(20)$ (**a**) and $RND(20)$ (**b**) statistics is shown for different mean dispersal distances and CNDD magnitude ($Q_C$) when $Q_G = 0.2$. **Right side:** The effect of dispersal distance on $EA(20)$ (**c**) and the $RND(20)$ (**d**) statistic is shown for different value of $\Delta_C$ and $Q_C$ when $Q_G = 0.2$. The legend for panels (**a**) and (**b**) is above (**b**) and for panels (**c**) and (**d**) are above (**d**).

#### Appendix S3: Relationship with abundance

Almost all results in this work are obtained for the average population in each regime, but how does the abundance of populations affect their dispersion? From the perspective of the *ED* and *EA*(20) statistics, the effect of abundance is generally unimodal, with intermediately abundant species being the most overdispersed (Figures 12a, S13a). In many cases only the increasing or decreasing phase can be observed, and sometimes no relationship emerges. From the perspective of *CE* and *RND*(20), there is a strong effect of reduced aggregation with abundance, as is commonly found in tropical forests (Figures S12b, S13b, Condit *et al.* 2000).

Although it is not easy to understand the mechanisms underlying such emergent patterns, we can identify three mechanisms that we believe govern the relationship between dispersion and abundance. The first, causing a decrease in overdispersion with abundance, is the convergence of DL to CSR and approaching a grid structure. As abundance increases, the dispersion of density-independent populations, which we use as the reference for *ED* and *EA*(20), converges to CSR, as can be seen by examining nearest-neighbor distances: In Figure S12c, nearest neighbor distances in CSR are represented by the dotted blue curve, and the other curves (except the purple) approach it as abundance increases. Asymptotically, demographic stochasticity wanes with higher abundance, and the population becomes similar to a diffusion process with torus boundary conditions, having a random dispersion. This convergence to CSR with abundance explains why, compared to CSR, both density-independent and density-dependent populations become less aggregated with abundance (Figures S12b, S13b), as is often observed in nature (Hubbell 1979; Condit *et al.* 2000). Moreover, as abundance increases, the nearest-neighbor distances of dispersal-limited populations also become closer to that of a square grid (purple line in Figure S12c), which we use as a reference representing "maximal overdispersion". This creates a geometric constraint on overdispersion: for example, the mean nearest neighbor distance of a rare species with $D = 60$ m. (blue curves in Figure S12c) can be longer than the DL null (thin solid blue curve) by a factor 3, but for a common species this would lead to the nearest-neighbor distances being longer than the distances of a square grid.

The second mechanism, also causing a decrease in overdispersion with abundance, is the signature of demographic history: Because all the populations in these simulations are attracted to a similar equilibrium abundance, a rare species is the result of a recent decline, and a common species is the result of a recent increase. Since reproduction happens near parents but mortality can happen anywhere and create isolated individuals, this mechanism increases the overdispersion of rare species and reduces the overdispersion of common species. Note that under the DL null populations are equally likely to increase or decrease at a given abundance, so this effect does not happened *on average* under the null. This mechanism is general and we expect it to operate also in more realistic scenarios when different species have different equilibrium abundances.

The third mechanism is responsible for the increase in overdispersion with abundance: As a density independent population (the reference for *ED* and *EA*) increases in abundance, it both expands in total range, occupying a larger proportion of the landscape (Figure S12d, thin line), and “compresses”, having more neighbors around each individual (Figure S12e, thin line). On the other hand, populations subject to substantial CNDD expand in range rapidly upon population growth, while their local density remains low (Figure S12d,e, thick lines), as CNDD makes them “incompressible”. This “compressibility difference” between density-dependent and density-independent populations with increasing abundance generates a positive effect of abundance on overdispersion.

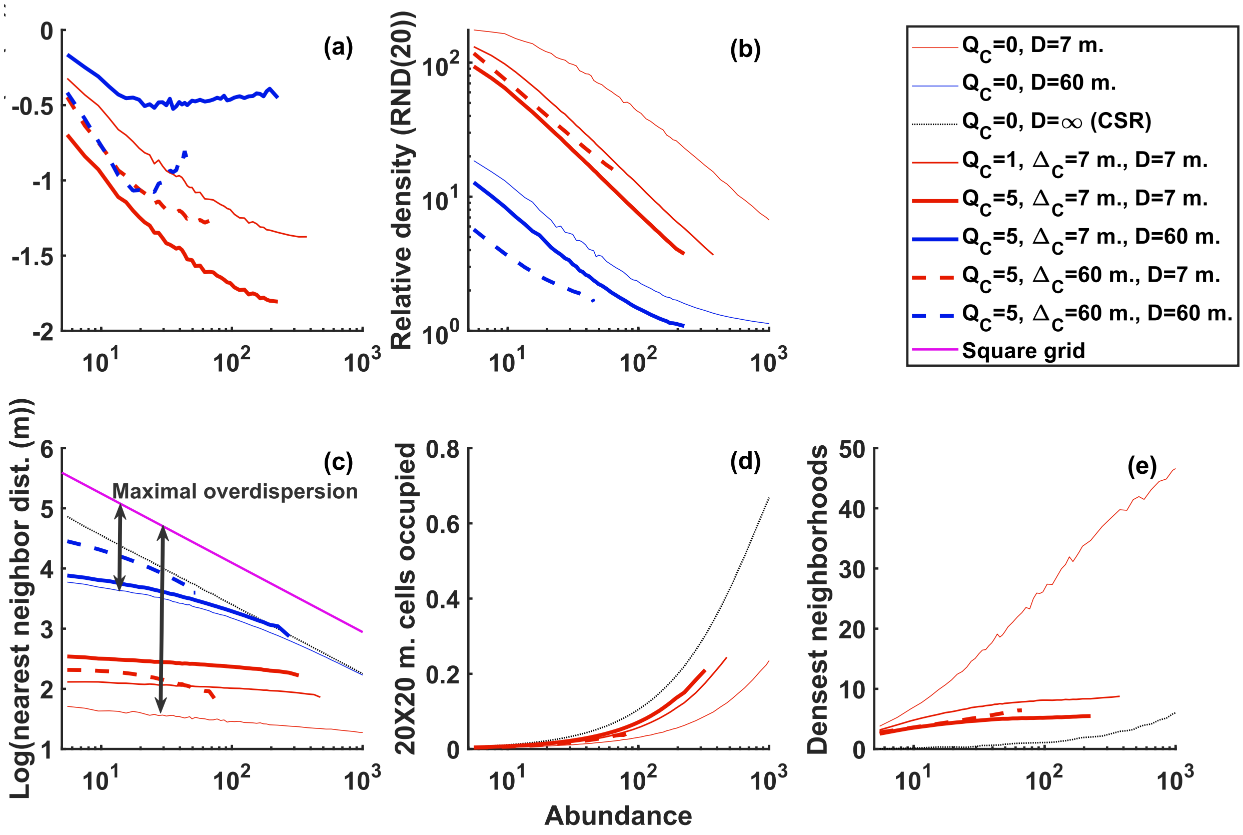


**Figure S12** Relationship between abundance and measures of dispersion and the underlying causes. In all panels, increasing line width indicates increasing added CNDD ($Q_C$), color represents dispersal distance (red: 7 meters, blue: 60 m. grey: ∞), dash represents the distance scale of CNDD ($\Delta_C$, solid: 7 m., dash: 60 m.) and in all cases there is no GNDD. The purple line represents the case of individuals distributed perfectly uniformly in space, i.e. as a grid, used as reference for maximal overdispersion, with nearest neighbor distance = $/\sqrt{N}$. (**a**) and (**b**) show the Excess neighborhood Abundance at 20 meters ($EA(20)$) and Relative Neighborhood Density at 20 meters ($RND(20)$). (**c**) compares the logarithm of mean nearest neighbor distances for several regimes, including density-independent cases, CSR and the square grid. Note that the difference between the density-independent case and the square grid (black arrow) is the maximal possible overdispersion ($ED$), which declines with increasing abundance. (**d**) and (**e**) present the proportion of 20X20 m. cells in the landscape that contain at least one individual; and 90[th] percentile of neighborhood densities within 20 m. of an individual, respectively, as measures of range and “compression”. The higher the strength of added CNDD, the weaker the relationship between abundance and the density of neighbors around the most crowded individuals, and in this sense the more “incompressible” species distributions are. The legend refers to all panels. Abundance bins with fewer than 100 observations were discarded.

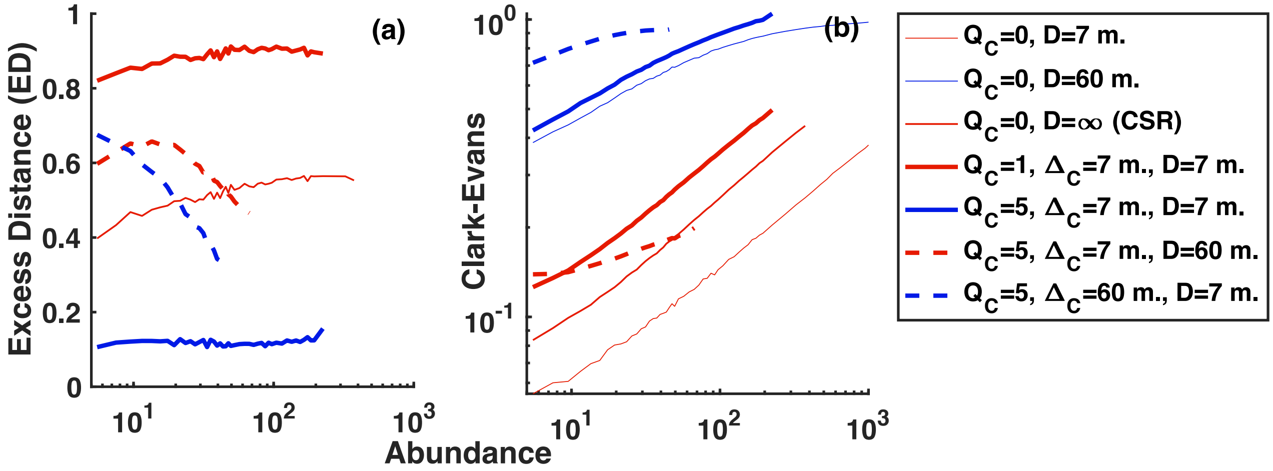


**Figure S13** Relationship between abundance and nearest neighbor statistics. In all panels, increasing line width indicates increasing $Q_C$, color represents dispersal distance (red: 7 meters, blue: 60 m. grey: ∞), dash represents $\Delta_C$ (solid: 7 m., dash: 60 m.) and in all cases $Q_G = 0$. (**a**) and (**b**) show the Excess nearest neighbor Distance (*ED*) the Clark-Evans statistic. The legend refers to all panels.
